# Decoding Immune Reconstitution Failure in People Living with HIV through Single-cell Genomics

**DOI:** 10.64898/2026.05.21.726797

**Authors:** Yi Wang, Shichen Yang, Yue Yuan, Mingli Zhu, Junjie Chen, Yong Bai, Shanshan Duan, Xuyang Shi, Tianyu Lu, Zongxing Yang, Zhuoli Huang, Dingyan Yan, Yuhui Zheng, Xuanchi Zhou, Jun Yan, Chang Liu, Wenhui Zhang, Yaling Huang, Chan Xu, Xiumei Lin, Xue Wang, Jinchuan Shi, Xiaojing Xu, Lei Ge, Jiefang Yin, Yu Feng, Hu Wan, Wangsheng Li, Bo Wang, Xin Hong, Xin Wang, Xiao Yang, Shourong Liu, Yi Zhao, Liang Chen, Yuxin Che, Wei Lyu, Bin Su, Runsheng Chen, Zhiwei Chen, Hanjie Li, Xin Liu, Ying Gu, Xin Jin, Longqi Liu, Xiangdong Wang, Jinsong Huang, Jianhua Yin, Jianhua Yu, Chuanyu Liu

## Abstract

Immune non-responders (INRs), a subset of people living with HIV (PLWH), fail to achieve full immune reconstitution and remain at increased risk of morbidity, mortality, and non-AIDS-related illnesses. As the mechanisms underlying this impaired immune recovery remain poorly understood, we performed single-cell multi-omics profiling on peripheral blood mononuclear cells from 43 INRs, 47 immune responders and 53 healthy donors. Our dataset comprises 2,744,009 transcriptomes and 1,226,658 chromatin-accessibility profiles across 58 identified immune cell types. INRs exhibited markedly elevated inflammatory signaling, increased apoptotic activity, and dysregulation of immune-activating ligand-receptor interactions. Additionally, we identified 2,996 interaction *cis*-eGenes and 5,938 *cis*-caPeaks, and validated key cell type-specific *cis*-xQTL effects associated with immune reconstitution failure. Furthermore, we identified 3,551 sc-eQTLs and 872 cell-state interaction eQTLs (ieQTLs) in CD4^+^ T cells. A genotype-specific upregulation of *OAS3* in INRs, likely regulated by STAT1, was identified based on these results. We developed scPRISM (single cell Predictive Reconstitution Immune Status Model), a multi-modal framework that integrates scRNA-seq and scATAC-seq data to perform the dual tasks of differentiating disease states and predicting gene expression from chromatin accessibility, thereby deciphering the cell type-specific *cis*-regulatory basis of the disease. Our study delineates the transcriptional and epigenomic landscape of immune dysregulation in INRs and provides new genetic insights into HIV-associated immune reconstitution failure. These findings offer a foundation for developing biomarkers and precision therapies to restore immune function in PLWH.

## Introduction

Acquired immunodeficiency syndrome (AIDS), caused primarily by the human immunodeficiency virus type 1 (HIV-1), remains a major global public health threat^1^. The disease is primarily characterized by progressive decline in CD4^+^ T lymphocyte (CD4^+^ T) counts and immune dysfunction, ultimately leading to a spectrum of immunodeficiency-associated disorders^2^. Currently, combination antiretroviral therapy (cART) is the most effective treatment strategy, suppressing viral replication and promoting immune reconstitution^3^. However, immune recovery is not achieved in all individuals. Approximately 15-30% of people living with HIV (PLWH) experience suboptimal restoration of CD4^+^ T cells, a condition termed immune non-response^2,4^. These immune non-responders (INRs) face elevated risks of hepatic diseases^5^, cardiovascular complications^6^, cytopenias^7^, dyslipidemia^8^, and increased morbidity and mortality^9^. Therefore, elucidating mechanisms underlying failed immune reconstitution in PLWH is urgently needed.

Single-cell RNA sequencing (scRNA-seq), single-cell assay for transposase-accessible chromatin sequencing (scATAC-seq) and whole genome sequencing (WGS) provide high-resolution insights into cellular diversity, enabling of gene expression and regulatory landscapes in immune cells^10–15^. Although recent advancements have identified factors influencing immune reconstitution outcomes in PLWH^11,14,16,17^, mechanisms of immune reconstitution failure remain largely unclear. Therefore, it is necessary to conduct a large-scale HIV-1 cohort study on integrating single-cell genomics.

As gene expression dynamically changes during immune cell activation and and differentiation, single-cell expression QTL (sc-eQTL) analyses have revealed context- and genotype-specific gene expression regulation^18–21^. Sc-eQTL analyses of blood cells have revealed context-specific eQTLs associated with various diseases^21–23^. To incorporates the cell state information obtained from single-cell-level data to test for interactions with the regulatory effects of genetic variants. The cell-state-interacting eQTLs or ieQTLs^21^, identifies eQTLs with an effect size that varies across different cell states, thereby capturing the state-specific gene expression regulation. Thus, integrating genetic variation and gene expression data offers a robust approach for elucidating causal mechanisms of gene regulation in immune cells, and for refining the interpretation of immune reconstruction failure-associated variants.

Peripheral blood mononuclear cells (PBMCs) serve as a representative immune cell population extensively utilized in the study of immunological responses and disease mechanisms^24^. Their changes have been associated with inflammatory conditions^25^, immune-mediated autoimmune disorders^26^, and viral pathogenesis, including HIV infection^27^. As such, PBMCs offer crucial information on systemic immune status and represent an essential resource for exploring immune reconstitution failure^28,29^.

In this study, we conducted a comprehensive single-cell multi-omics analysis, including scRNA-seq, scATAC-seq, and WGS on PBMCs from 53 healthy donors (HDs), 47 immune responders (IRs) and 43 INRs. By integrating these datasets, we constructed cell type-specific regulatory networks and characterized the impact of genetic variation on chromatin accessibility and gene expression. Our findings revealed multiple immune pathways, regulatory factors and genetic variants that influenced immune reconstitution outcomes in PLWH. Additionally, we developed scPRISM (single cell Predictive Reconstitution Immune Status Model), a multi-modal framework that integrates scRNA-seq and scATAC-seq data to simultaneously classify disease states, predict gene expression from chromatin accessibility, and infer the underlying cell type-specific *cis*-regulatory links. This study not only provides a valuable single-cell multi-omics resource for studying INRs but also offers novel insights into the molecular underpinnings of gene regulation and genetic risk in immune reconstruction failure. Our findings may also facilitate the development of personalized biomarkers and precision therapies to enhance immune recovery.

## Results

### Study design and overview for decoding immune reconstitution failure in INRs

To characterize immune reconstitution failure in PLWH, we enrolled 43 INRs, 47 IRs, and 53 HDs from Hangzhou Xixi Hospital. PBMCs were collected from each participant after obtaining ethical approval and informed consent. Multi-omics sequencing, including scRNA-seq, scATAC-seq, and WGS, was performed on PBMCs from all 143 adults. Using integrated WGS data, we mapped xQTLs associated with INRs and constructed both scPRISM-CLIN and scPRISM-EXPR models (Fig. 1a). The cohort consisted exclusively of male participants, ranging in age from 26 to 76 years (Supplementary Fig. 1a). Age distribution was comparable among INRs, IRs, and HDs, and treatment duration did not differ significantly between INRs and IRs. However, CD4^+^ T cell counts and CD4^+^/CD8^+^ T cell ratios were significantly lower in INRs than in IRs and HDs (Fig. 1b). Additional clinical characteristics are summarized in Supplementary table 1.

**Fig. 1.**
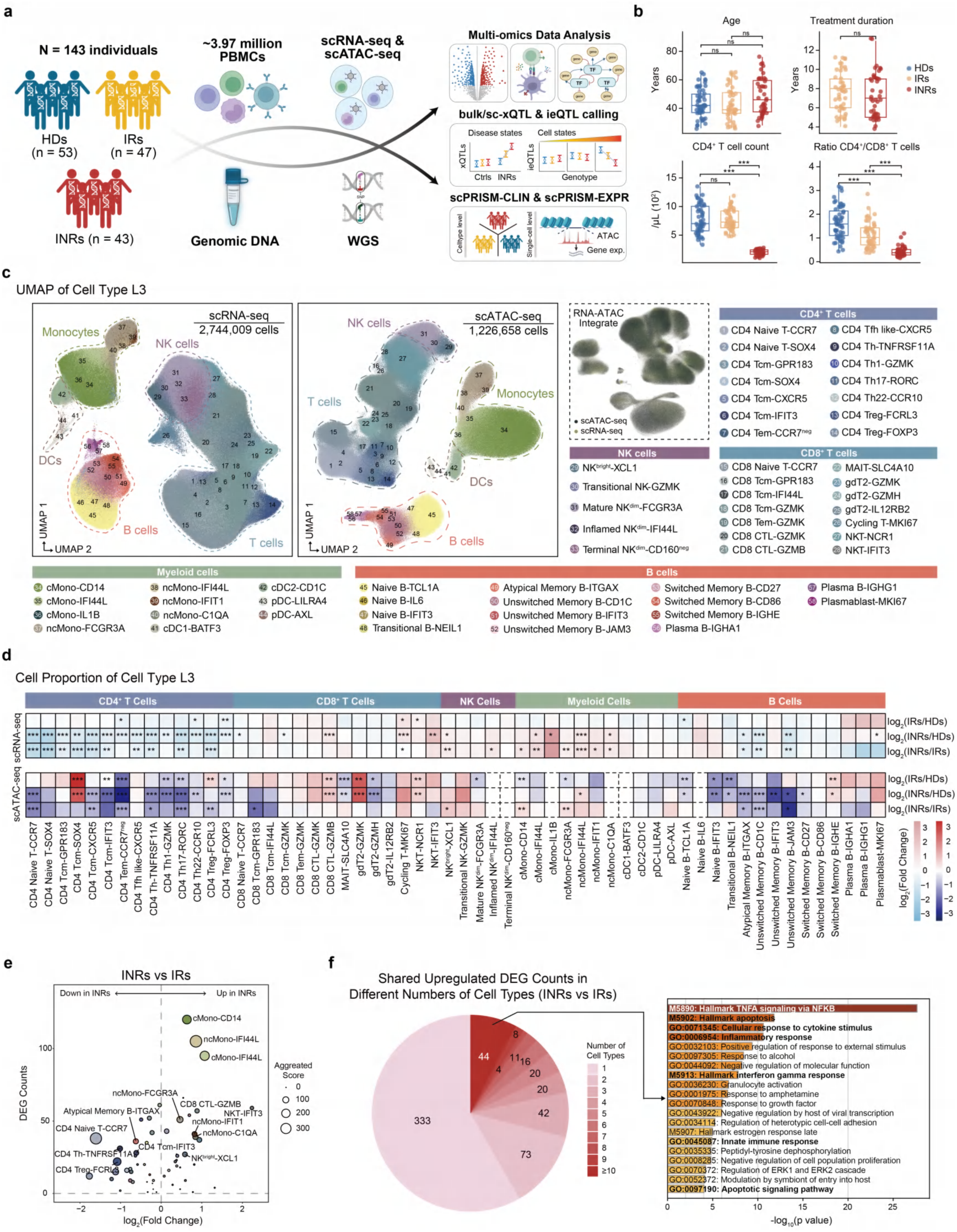
Study design and overview of single-cell genomics decoding immune reconstitution failure in HIV. **a**, Overview of the study design, including collecting peripheral blood samples from 53 HDs, 47 IRs, and 43 INRs, capturing multi-omics data and performing comprehensive analysis. **b**, Box plots showing the differences among HDs, IRs, and INRs in terms of age, treatment duration, CD4^+^ T cell count, and CD4^+^/CD8^+^ T cell count ratio. **c**, Uniform Manifold Approximation and Projection (UMAP) plots showing 2,744,009 immune cells from scRNA-seq data and 1,226,658 immune cells from scATAC-seq data that have been reduced, clustered, and annotated. At L3, cells are classified into 58 immune cell subtypes, with each cell type represented by a different color. The combined UMAP plot shows the integration of scRNA-seq and scATAC-seq. **d**, Heatmap showing the differences in cell proportions of L3 (58 cell subtypes) between HDs, IRs, and INRs in scRNA-seq data and scATAC-seq data. **e**, Bubble plot comparing changes in cell proportions and differentially expressed gene (DEG) counts between INRs and IRs. Bubble size represents a composite measure of change, defined as -log_10_(P adjust of cell proportion change) × DEG Counts. P adjust of cell proportion change was determined using the Wilcoxon rank sum test with Benjamini-Hochberg correction. **f**, The pie chart showing the number of DEGs shared up-regulated in different numbers of cell types (INRs versus IRs), and pathway enrichment of genes shared up-regulated in ≥ 10 cell types.

To ensure high-quality single-cell data, we applied a stringent quality control pipeline to PBMCs, which included the removal of doublets (Supplementary Fig. 1b, c) and batch effect correction during clustering (Supplementary Fig. 1d). For accurate immune cell phenotyping, we performed iterative dimensionality reduction and clustering. Uniform Manifold Approximation and Projection (UMAP) visualization represents 2,744,009 immune cells from scRNA-seq data after dimensionality reduction, clustering, and annotation (Fig. 1c). Guided by lineage marker gene expression, PBMCs were first categorized into five major Level 1 (L1) cell types: CD4^+^ T cells, CD8^+^ T and unconventional T cells, NK cells, myeloid cells, and B cells (Fig. 1c), following the classification framework established by Silvia Domcke et al.^30^. These broad lineages were subsequently subclustered to delineate developmental “cell states” and “cell types” based on gene expression patterns. Using canonical marker genes and cell type-specific signatures, we annotated 28 immune cell subtypes at Level 2 (L2) (Supplementary Fig. 1e) and 58 subtypes at Level 3 (L3) (Fig. 1c). The scATAC-seq datasets were annotated by integrating them with scRNA-seq data via the scglue model^31^, enabling transfer of cell type labels from transcriptomic to chromatin accessibility data. This integration yielded 27 L2 subtypes (Supplementary Fig. 1f) and 52 L3 subtypes (Fig. 1c). Collectively, our PBMC atlas establishes a unified three-tier annotation framework (Supplementary Fig. 2a) across both omics modalities, incorporating RNA and ATAC cell counts (Supplementary Fig. 2b) as well as marker gene expression and accessibility profiles (Supplementary Fig. 2c) spanning 58 immune cell types.

We analyzed immune functions using L2 and L3 cell subtypes and assessed compositional differences among INRs, IRs, and HDs. Significant disparities in the proportions of 22 (scRNA-seq) and 12 (scATAC-seq) L3 cell types were observed between INRs and IRs (Fig. 1d). Compared to IRs and HDs, INRs showed reduced proportions of CD4^+^ T cell and B cell subsets, alongside elevated proportions of NK and myeloid cell subsets at both L2 and L3 hierarchical levels (Fig. 1d and Supplementary Fig. 1g). Differential gene expression analysis in L3 transcriptomes revealed significantly more DEGs in INRs versus IRs, with myeloid cells exhibiting a notably higher number of up-regulated DEGs than other lineages (Supplementary Fig. 3a). Most immune cell types in INRs displayed more up-regulated than down-regulated DEGs (Supplementary Fig. 3a). Transcriptional profiling across 58 cell types identified pronounced changes in INR cell subsets with altered frequencies, such as CD4 Naive T-CCR7, CD4 Treg-FCRL3, cMono-CD14, ncMono-FCGR3A, CD8 CTL-GZMB, NK^bright^-XCL1, and Atypical Memory B-ITGAX (Fig. 1e). Transcriptional divergence was more extensive between INRs and HDs (Supplementary Fig. 3b), whereas only CD8 CTL-GZMB differed notably between IRs and HDs (Supplementary Fig. 3c). Gene Ontology (GO) and HALLMARK pathway enrichment analyses^11^ of commonly up-regulated DEGs in INRs versus IRs revealed significant enrichment in processes including TNF-α signaling via NF-κB, apoptosis, interferon-gamma response, and inflammatory response (Fig. 1f). Finally, we characterized the genomic distribution of differentially accessible peaks (DAPs) from five L1 cell types across INRs, IRs, and HDs. In INRs, both up- and down-regulated DAPs were predominantly located in promoter, distal intergenic, and intronic regions relative to IRs and HDs (Supplementary Fig. 3d,e).

### CD4^+^ T cells in INRs exhibited excessive activation and enhanced apoptosis

Our transcriptome analysis resolved PBMCs into 14 CD4^+^ T cell clusters (Fig. 2a and Supplementary Fig. 4a). Notably, we identified 60 immune-related DEGs that were significantly up-regulated in the CD4^+^ T cell subsets of INRs when compared to IRs and HDs (Fig. 2b). Functional annotation placed these DEGs into five key categories: antiviral response, cell activation and immune activation, inflammatory response, cytokine signaling, and apoptosis-related pathways (Fig. 2b). Strikingly, the DEGs correlated with DAPs demonstrated an enrichment pattern that aligned with the transcriptional alterations, supporting a potential regulatory relationship (Supplementary Fig. 4b).

**Fig. 2.**
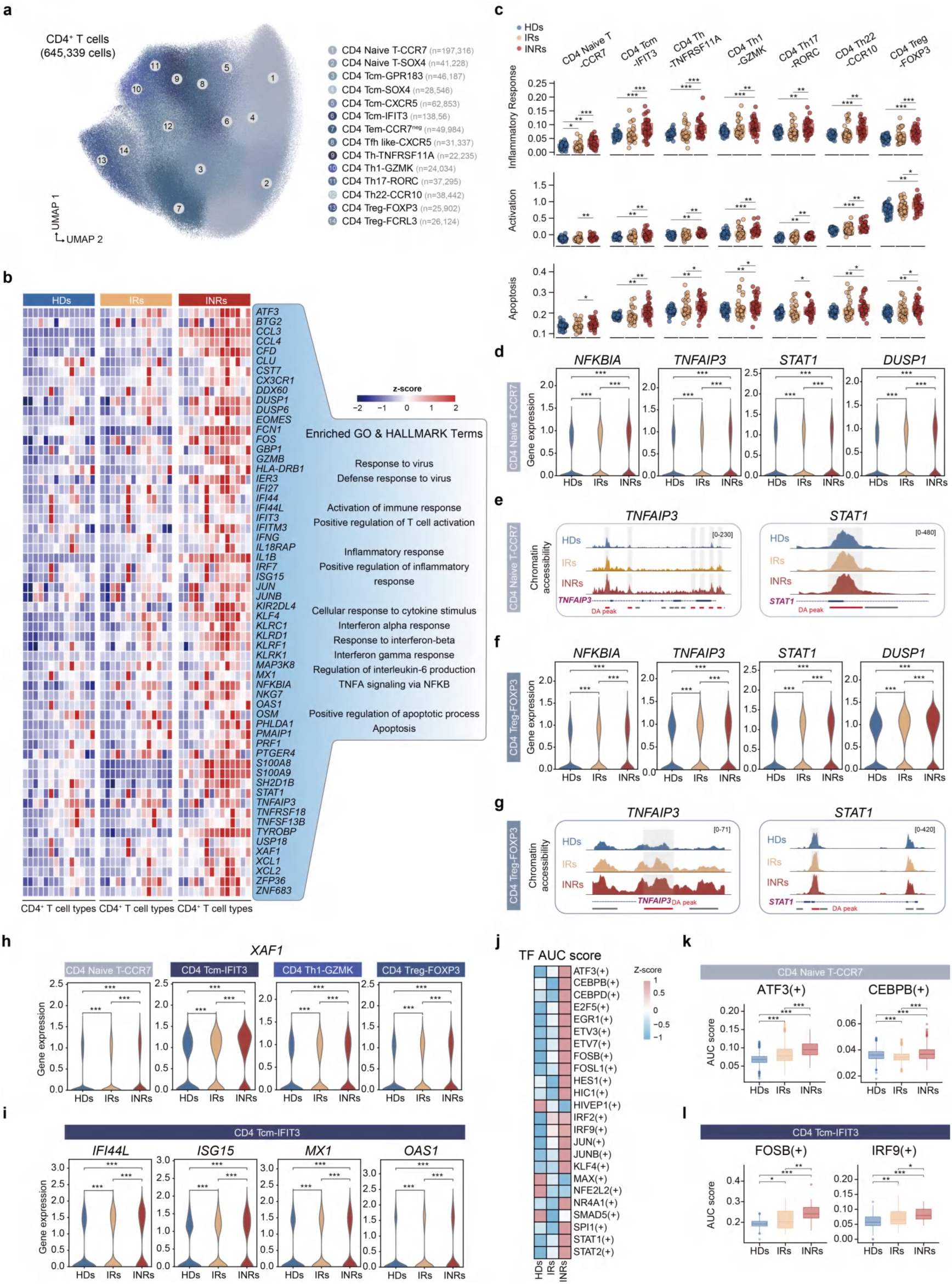
CD4+ T cells exhibited excessive activation and apoptosis in INRs. **a,** The UMAP plot showing 645,339 CD4+ T cells from scRNA-seq data, which are divided into 14 cell subtypes, with each cell type represented by a different color. **b**, Heatmaps showing the scaled mean expression of DEGs (INRs versus IRs) of 14 CD4^+^ T cell types in HDs, IRs, and INRs, and the pathways enriched for these DEGs on the right. **c**, Boxplots showing the scores of Inflammatory Response, Activation, and Apoptosis for seven CD4^+^ T cell types in HDs, IRs and INRs. Different groups are shown in different colors, the horizontal line represents the median, and the whiskers extend to the furthest data point within a maximum of 1.5 times the interquartile range. **d**, Violin plots showing gene expression of *NFKBIA*, *TNFAIP3*, *STAT1*, and *DUSP1* in CD4 Naive T-CCR7 in HDs, IRs, and INRs. Different groups are shown in different colors. **e**, scATAC-seq tracks revealing gene structure and significant differentially accessible peaks (DA peaks) for *TNFAIP3* and *STAT1* in CD4 Naive T-CCR7 in HDs, IRs, and INRs. Different groups are shown in different colors. **f**, Violin plots showing gene expression of *NFKBIA*, *TNFAIP3*, *STAT1*, and *DUSP1* in CD4 Treg-FOXP3 in HDs, IRs, and INRs. Different groups are shown in different colors. **g**, scATAC-seq tracks revealing gene structure and significant DA peaks for *TNFAIP3* and *STAT1* in CD4 Treg-FOXP3 in HDs, IRs, and INRs. Different groups are shown in different colors. **h**, Violin plots showing gene expression of *XAF1* in CD4 Naive T-CCR7, CD4 Tcm-IFIT3, CD4 Th1-GZMK and CD4 Treg-FOXP3 in HDs, IRs, and INRs. Different groups are shown in different colors. **i**, Violin plots showing gene expression of *IFI44L*, *ISG15*, *MX1*, and *OAS1* in CD4 Tcm-IFIT3 in HDs, IRs, and INRs. Different groups are shown in different colors. **j**, Heatmap showing scaled mean AUC score of differential transcription factors (TFs) in CD4^+^ T cells in HDs, IRs, and INRs. **k**, Box plots showing the AUC values of ATF and CEBPB in CD4 Naive T-CCR7 in HD, IRs, and INRs. Different groups are shown in different colors. **l**, Box plots showing the AUC values of FOSB and IRF9 in CD4 Tcm-IFIT3 in HD, IRs, and INRs. Different groups are shown in different colors.

Pathway scores for inflammatory response, immune activation, apoptosis, and cytokine signaling were significantly elevated in most CD4^+^ T cell subtypes from INRs compared to IRs or HDs (Fig. 2c and Supplementary Fig. 4c). These findings suggest that CD4^+^ T cells in INRs exist in a state of heightened activation and increased apoptosis in vivo, likely driven by aberrant inflammatory stimuli, ultimately contributing to the observed reduction in CD4^+^ T cell counts. Previous studies have indicated that dysregulated inflammation can cause T cell over-activation and aberrant apoptosis in PLWH under ART control^32^. Moreover, excessive interferon-gamma (IFN-γ) release has been implicated in the pathogenesis of chronic inflammatory conditions and is known to promote cell death via apoptosis and necroptosis^33^. To explore these interactions, we examined the relationships among the four pathways. In CD4 Naive T-CCR7 and CD4 Treg-FOXP3 subsets, inflammatory response positively correlated with both immune activation and IFN-γ response; IFN-γ response was also positively associated with activation and apoptosis. Notably, these correlations were more pronounced in INRs than in IRs or HDs (Supplementary Fig. 4d, e).

We next characterized key gene expression and chromatin accessibility signatures in CD4^+^ T cell from INRs. Expression and chromatin accessibility of *NFKBIA* and *TNFAIP3*, negative regulators of NF-κB signaling, were elevated in CD4 Naive T-CCR7, CD4 Tcm-IFIT3, and CD4 Treg-FOXP3 in INRs (Fig. 2d-g and Supplementary Fig. 4f), suggesting hyperactivation of the NF-κB pathway. Given that INRs exhibit reduced proliferation and impaired function of CD4^+^ T cells^34^, we also observed upregulation of the pro-apoptotic gene *XAF1* in multiple CD4^+^ T cell subsets (Fig. 2h), which may exacerbate CD4^+^ T cell loss.

Persistent type I interferon (IFN-I) signaling was evident in CD4^+^ T cells of INRs^35^, consistent with HIV-induced chronic immune activation that drives elevated *STAT1* and interferon-stimulated gene (ISG) expression, forming an interferon-high phenotype^36,37^. *STAT1* exhibited increased expression and chromatin accessibility in CD4 Naive T-CCR7 and Treg-FOXP3 (Fig. 2d-f and Supplementary Fig. 4f), while ISGs such as *IFI44L*, *ISG15*, *MX1*, and *OAS1* were upregulated in CD4 Tcm-IFIT3 (Fig. 2i). Additionally, *DUSP1*, a stress-responsive regulator of the MAPK pathway that may mitigate inflammatory damage^38^, was upregulated in CD4 Naive T-CCR7, CD4 Treg-FOXP3, and CD4 Tcm-IFIT3 of INRs (Fig. 2d-f and Supplementary Fig. 4f). Chronic low-grade inflammation persists in HIV despite viral suppression^39^, and sustained NF-κB and STAT1 signaling maintains high ISG expression, contributing to chronic immune dysregulation^40^.

To further explore transcriptional regulation in INRs, we used SCENIC+^41^ to integrate scATAC-seq and scRNA-seq data and construct an enhancer-driven gene regulatory network (GRN). We identified 26 TFs with differential regulatory activity between INRs and IRs in at least one CD4^+^ T cell subtype (Fig. 2j). Most showed enhanced activity in INRs, including members of the AP-1 (ATF3, JUN, JUNB, and FOSB), IRF (IRF2, IRF5, and IRF9), and STAT (STAT1 and STAT2) families. AP-1 is a central mediator of inflammatory and stress responses whose overactivation may promote chronic inflammation and immune dysregulation^42^. IRF9 regulates IFN-I signaling^43^, and its sustained activity may perpetuate an inflammatory microenvironment^44^. STAT1 drives transcription of IFN-induced genes, and its hyperactivation may sustain a pro-inflammatory state that impedes T cell functional recovery^45^. A focused regulatory network of 16 TFs and their targets revealed co-regulatory patterns, with AP-1 targets enriched in pro-inflammatory processes and IRF9/STAT1 targets linked to interferon signaling and chronic inflammation (Supplementary Fig. 4g).

At L3 resolution, area under the curve (AUC) analysis confirmed upregulated pro-inflammatory and IFN-related TFs in INRs. CD4 Naive T-CCR7 and CD4 Tcm-IFIT3 exhibited elevated AUC values for AP-1 TFs (ATF3, JUN, FOSB) and IRF9 (Fig. 2k-l and Supplementary Fig. 4h, i). In CD4 Naive T-CCR7, CEBPB, a bZIP TF associated with inflammatory responses and NF-κB activation^46^, was also upregulated (Fig. 2k). CEBPB amplifies NF-κB-dependent inflammation by regulating cytokines such as IL-6 and TNF-α, thereby contributing to sustained immune activation that correlates with IRF activity^47^.

### Sustained activation of CD8^+^ T cells and NK cells in INRs

Transcriptomic analysis of CD8^+^ T cells identified 14 distinct clusters (Supplementary Fig. 5a, b). Pathway activity scores for inflammatory response, interferon-gamma response, and cytotoxicity were significantly elevated in most CD8^+^ T cell subtypes from INRs compared to IRs and HDs (Supplementary Fig. 5c). Consistent with this, both *TNFAIP3* expression and chromatin accessibility were higher in CD8 CTL-GZMB and NKT-NCR1 subsets of INRs (Supplementary Fig. 5d, e).

Transcriptome clustering of NK cells delineated five distinct subsets (Supplementary Fig. 5f, g). Pathway enrichment revealed significant activation of TNF-α signaling via NF-κB, interferon gamma response, and inflammatory response in INRs (Supplementary Fig. 5h). *NFKBIA* expression was increased in NK^bright^-XCL1 and Mature NK^dim^-FCGR3A subsets of INRs, along with elevated expression of inflammatory genes such as *S100A9*, *DUSP1*, *IFITM3*, and *CCL4* (Supplementary Fig. 5i, j); these genes are associated with immune dysfunction^48–51^.

To examine intercellular communication, we inferred ligand-receptor interactions using CellPhoneDB^52^. While the proportions of CD8 CTL-GZMK and Mature NK^dim^-FCGR3A remained unchanged, interactions between CD8 CTL-GZMK and 10 subsets via IFNG-IFNGR1 were significantly enhanced in INRs. Similarly, IFNG-IFNGR2 interactions were observed between CD8 CTL-GZMK or Mature NK^dim^-FCGR3A and cMono-CD14, ncMono-FCGR3A, Unswitched Memory B-CD1C, and Unswitched Memory B-JAM3 (Supplementary Fig. 5k). IFN-γ is encoded by the *IFNG* and plays a crucial role in the occurrence and development of inflammatory and immune-mediated diseases^53^.

#### Myeloid cells overactivation and inflammatory microenvironment in INRs

Re-clustering of monocytes and dendritic cells based on transcriptomic data identified 11 distinct subsets, which were annotated using canonical marker genes (Fig. 3a and Supplementary Fig. 6a). Differential expression analysis revealed a consistent upregulation of inflammation-related genes in INRs compared to IRs and HDs (Fig. 3b). GO enrichment analysis across five myeloid subsets identified 22 top pathways upregulated in INRs, most of which were related to immune and inflammatory responses and were commonly shared among subsets (Supplementary Fig. 6b). Genes associated with differentially accessible peaks (DAPs) showed enrichment patterns consistent with those of differentially expressed genes (DEGs) (Supplementary Fig. 6c). Moreover, pathway scores for inflammatory response and interferon-gamma response were significantly elevated in seven myeloid subtypes from INRs relative to those from IRs and HDs (Fig. 3c).

**Fig. 3.**
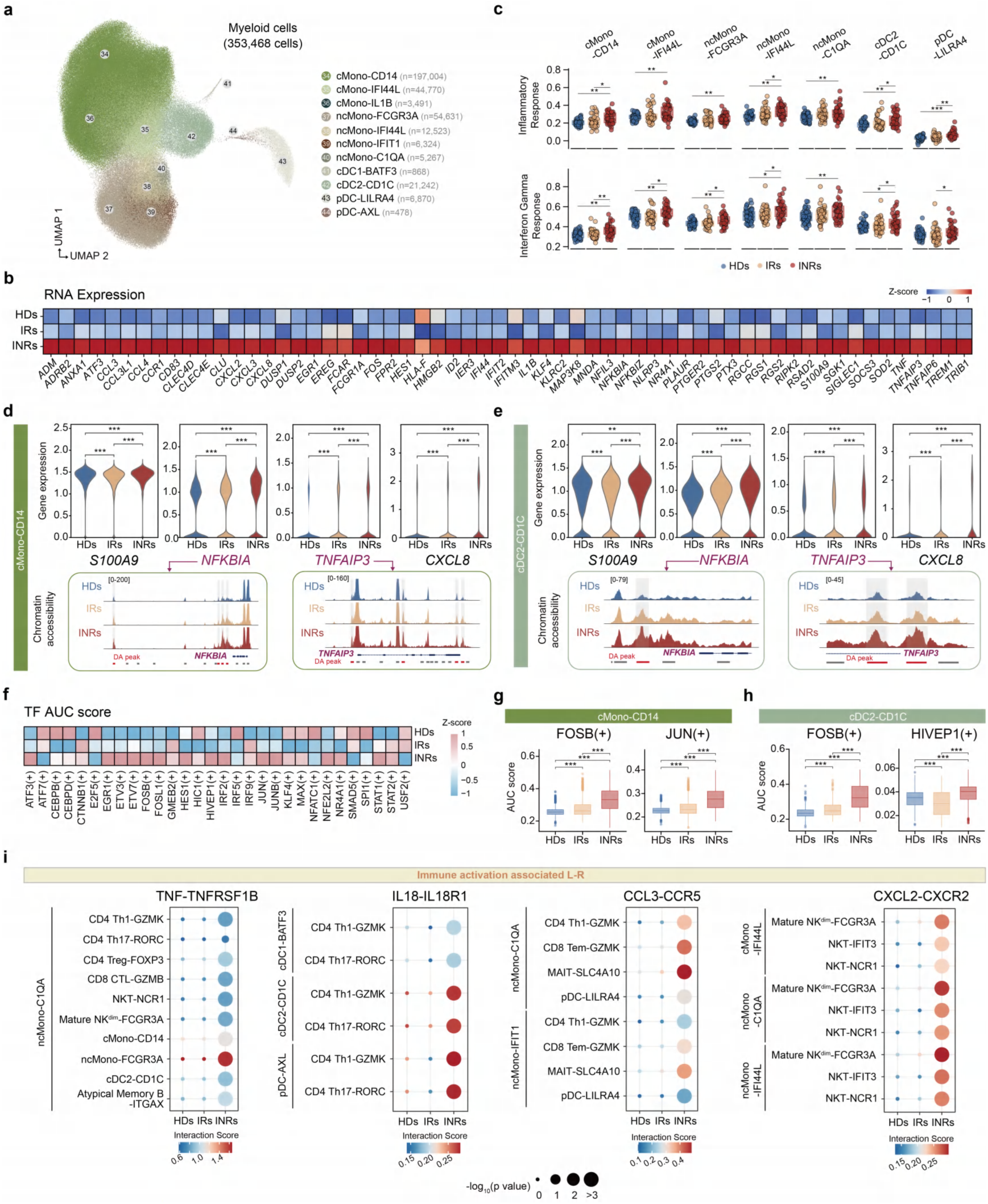
Aberrant activation of inflammatory responses in myeloid cells in INRs. **a**, The UMAP plot showing 353,468 myeloid cells from the scRNA-seq data, which are divided into 11 cell subtypes, with each cell type represented by a different color. **b**, Heatmaps showing the scaled mean expression of DEGs (INRs versus IRs) of 12 myeloid cell types in HDs, IRs, and INRs. **c**, Boxplots showing the scores of Inflammatory Response and Interferon Gamma Response for seven myeloid cell types in HDs, IRs, and INRs. Different groups are shown in different colors, the horizontal line represents the median, and the whiskers extend to the furthest data point within a maximum of 1.5 times the interquartile range. **d**, Violin plots showing gene expression of *S100A9*, *NFKBIA*, *TNFAIP3*, and *CXCL8* and scATAC-seq tracks revealing gene structure and significant DA peaks for *NFKBIA* and *TNFAIP3* in cMono-CD14 in HDs, IRs and INRs. Different groups are shown in different colors. **e**, Violin plots showing gene expression of *S100A9*, *NFKBIA*, *TNFAIP3*, and *CXCL8* and scATAC-seq tracks revealing gene structure and significant DA peaks for *NFKBIA* and *TNFAIP3* in cDC2-CD1C in HDs, IRs, and INRs. Different groups are shown in different colors. **f**, Heatmap showing scaled mean AUC score of differential TFs in myeloid cells in HDs, IRs, and INRs. **g**, Box plots showing the AUC values of FOSB and JUN in cMono-CD14 in HD, IRs, and INRs. Different groups are shown in different colors. **h**, Box plots showing the AUC values of FOSB and HIVEP1 in cDC2-CD1C in HDs, IRs, and INRs. Different groups are shown in different colors. **i**, The bubble plots showing the changes in potential immune activation-related ligand-receptor pairs between immune cell subsets in HDs, IRs, and INRs, including TNF-TNFRSF1B, IL18-IL18R1, CCL3-CCR5, and CXCL2-CXCR2. The color of the dots indicates the interaction score calculated using CellPhoneDB, and the size of the dots indicates the -log_10_(p value).

Among key regulators of the NF-κB pathway, both expression and chromatin accessibility of *NFKBIA* and *TNFAIP3* were elevated in cMono-CD14 and cDC2-CD1C from INRs. The inflammatory factor *S100A9* also showed higher expression in cMono-CD14 (Fig. 3d), cDC2-CD1C (Fig. 3e), and ncMono-FCGR3A (Supplementary Fig. 6d). Additionally, the chemokine *CXCL8* was upregulated in cMono-CD14 and cDC2-CD1C (Fig. 3d, e), a change that may promote neutrophil recruitment and contribute to a sustained inflammatory microenvironment^54,55^. Among proinflammatory cytokines, *IL1B* exhibited increased expression and chromatin accessibility in cMono-IFI44L, ncMono-IFI44L, and cDC2-CD1C (Supplementary Fig. 6e). Together, these alterations highlight a molecular landscape consistent with an elevated risk of chronic inflammation in INRs compared to IRs^56^.

We identified 30 TFs with differential regulatory activity in myeloid cells between INRs and IRs (Fig. 3f). Among these, members of the AP-1 family, including JUN, JUNB, and FOSB, displayed enhanced activity in cMono-CD14, cDC2-CD1C, cMono-IFI44L, ncMono-FCGR3A, and ncMono-IFI44L (Fig 3g-h and Supplementary Fig. 6f-h). The AP-1 complex is a key mediator of inflammatory signaling and stress responses, and its overactivation may contribute to chronic inflammation and disruption of immune homeostasis^42^. In addition, HIVEP1, a zinc finger transcription factor related to NF-κB signaling, promotes the upregulation of TNF-α, IL-6, and IL-1β via NF-κB binding sites, thereby facilitating a chronic inflammatory state^57^. HIVEP1 exhibited significantly increased activity in the same monocyte and dendritic cell subsets from INRs (Fig. 3h and Supplementary Fig. 6f-h).

Cell-cell interaction analysis further revealed enhanced communication between myeloid and other immune cells in INRs. Specifically, ncMono-C1QA showed stronger interactions with ten immune cell types via TNF-TNFRSF1B signaling (Fig. 3i), along with increased TNF-TNFRSF1A interactions involving six monocyte and four dendritic cell subsets (Supplementary Fig. 6i). These alterations suggest dysregulation of TNF-related signaling pathways, which may promote chronic inflammation and immune imbalance^58^. Moreover, cDC1-BATF3, cDC2-CD1C, and pDC-AXL exhibited enhanced IL18-IL18R1 signaling toward CD4 Th1-GZMK and CD4 Th17-RORC (Fig. 3i). As IL-18 is a pro-inflammatory cytokine known to amplify IFN-γ production in Th1 cells and promote Th17-driven autoimmune inflammation^59^, its heightened signaling in INRs may contribute to aberrant CD4^+^ T cell activation and exacerbate chronic inflammation.

### Abnormal function of B cells in INRs

Transcriptomic analysis identified 14 distinct B cell clusters (Supplementary Fig. 7a, b). Pathway enrichment analysis revealed significant activation in B cells from INRs of TNF-α signaling via NF-κB, inflammatory response, humoral immune response, and interferon-gamma response (Supplementary Fig. 7c).

Notably, *NR4A2*, a transcription factor implicated in chronic inflammation^60^, showed elevated expression and chromatin accessibility in Atypical Memory B-ITGAXand Unswitched Memory B-CD1C from INRs compared to IRs and HDs. In these B cell subsets, expression of *DUSP1*, *NFKBIA*, and *IFI30* was also upregulated (Supplementary Fig. 7d, e), and *IFI30* exhibited increased chromatin accessibility in Atypical Memory B-ITGAX (Supplementary Fig. 7e). Furthermore, INRs displayed heightened expression of the inflammatory genes *PPP1R15A*^61^ and *RGS1*^62^ specifically in Atypical Memory B-ITGAX (Supplementary Fig. 7d).

We identified 21 TFs exhibiting enhanced regulon activity in B cells from INRs relative to IRs and HDs (Supplementary Fig. 7f). Concurrently, we observed reduced proportions of specific B cell subsets in INRs (Fig. 1d). Cell-cell communication analysis revealed weakened CD86-CTLA4 ligand-receptor interactions between these B cell subsets and CD4 Tcm-IFIT3 or CD8 Tcm-IFI44L in INRs (Supplementary Fig. 7g). Given that CTLA4 serves as a critical negative regulator of T cell activation, impaired CD86-CTLA4 signaling in INRs may attenuate its suppressive effect on T cell hyperactivation, thereby exacerbating inflammation and immune dysregulation. This interpretation aligns with previous studies linking defective CTLA4 signaling to disrupted T cell homeostasis and complex immune dysregulation syndromes^63,64^.

### Identification of the relationship between clinical factors and immune reconstitution failure characteristics

The INR cohort consisted of 43 male participants aged 28-76 years, with treatment durations ranging from 4 to 13 years (Supplementary Fig. 8a). ART regimens included integrase strand transfer inhibitor (INSTI)-based therapies (n = 23) and non-INSTI regimens (n = 20). CD4^+^ T cell counts varied from 101 to 282 cells/μL (Supplementary Fig. 8a).

Correlation analyses linked four clinical factors with 12 DEGs, four signature pathways, and RNA-based cell proportions across seven immune cell subsets (Supplementary Fig. 8b). In CD4 Naive T-CCR7, *DUSP1* expression and inflammatory response scores showed a positive correlation with age, while inflammatory pathway scores were negatively correlated with CD4^+^ T cell counts (Supplementary Fig. 8c). In cMono-CD14, cDC2-CD1C, and Atypical Memory B-ITGAX subsets, *NFKBIA* expression positively correlated with age (Supplementary Fig. 8d-f). Similarly, *IL1B* (Supplementary Fig. 8d), *TNFAIP3* (Supplementary Fig. 8e), and *NR4A2* (Supplementary Fig. 8f) expression also increased with age in these respective subsets.

Levels of interferon-stimulated genes (*ISG15* and *IFI44L*) were lower in INSTI-treated individuals compared to those receiving non-INSTI regimens within cMono-CD14 and cDC2-CD1C subsets (Supplementary Fig. 8d, e). A negative correlation between RNA-derived cell proportion and age was observed in Atypical Memory B-ITGAX (Supplementary Fig. 8f).

### Identification of nominal *cis*-xQTLs in immune cell types

To assess the influence of common genetic variants on gene expression and chromatin accessibility, we performed *cis*-eQTL and *cis*-caQTL mapping across immune cell subpopulations. Our analysis identified 2,957 *cis*-eGenes across 51 cell subsets (Supplementary Fig. 9a), of which 88% were cell type-specific (detected in only one subset) and 12% were shared across multiple subsets (Supplementary Fig. 9b). Similarly, we detected 14,635 *cis*-caPeaks across 31 cell subsets (Supplementary Fig. 9c), with 68% being cell type-specific and 32% shared (Supplementary Fig. 9d). We also summarized the numbers of genes/peaks and pseudobulk samples used in the *cis*-xQTL mapping (Supplementary Fig. 9e, f).

#### Immune reconstitution failure-related *cis*-xQTLs in different cell types

We performed a systematic profiling of *cis*-xQTLs across immune cell subsets from INRs and IRs+HDs to dissect context-specific eQTLs that were unique to a particular group or exhibited divergent effect directions or magnitudes among the two groups. We identified 2,996 interaction *cis*-eGenes, with 29% restricted to a single cell type (Fig. 4a, b), and 5,938 interaction *cis*-caPeaks, of which 9% were cell type-specific (Fig. 4c, d). Notably, while nominal *cis*-xQTL numbers correlated with cell type abundance (Supplementary Fig. 10a), interaction *cis*-xQTL counts showed no correlation with cellular proportions (Supplementary Fig. 10b). Furthermore, a pronounced divergence in genomic distribution was observed: interaction *cis*-caPeaks were highly enriched in promoters (71.9%), whereas nominal *cis*-caPeaks were more evenly distributed between promoters and introns (Supplementary Fig. 10c). Functional analysis showed that these interaction *cis*-eGenes are enriched in key immunological pathways such as innate immune response, cytokine-mediated signaling, and cell activation (Supplementary Fig. 10d).

**Fig. 4.**
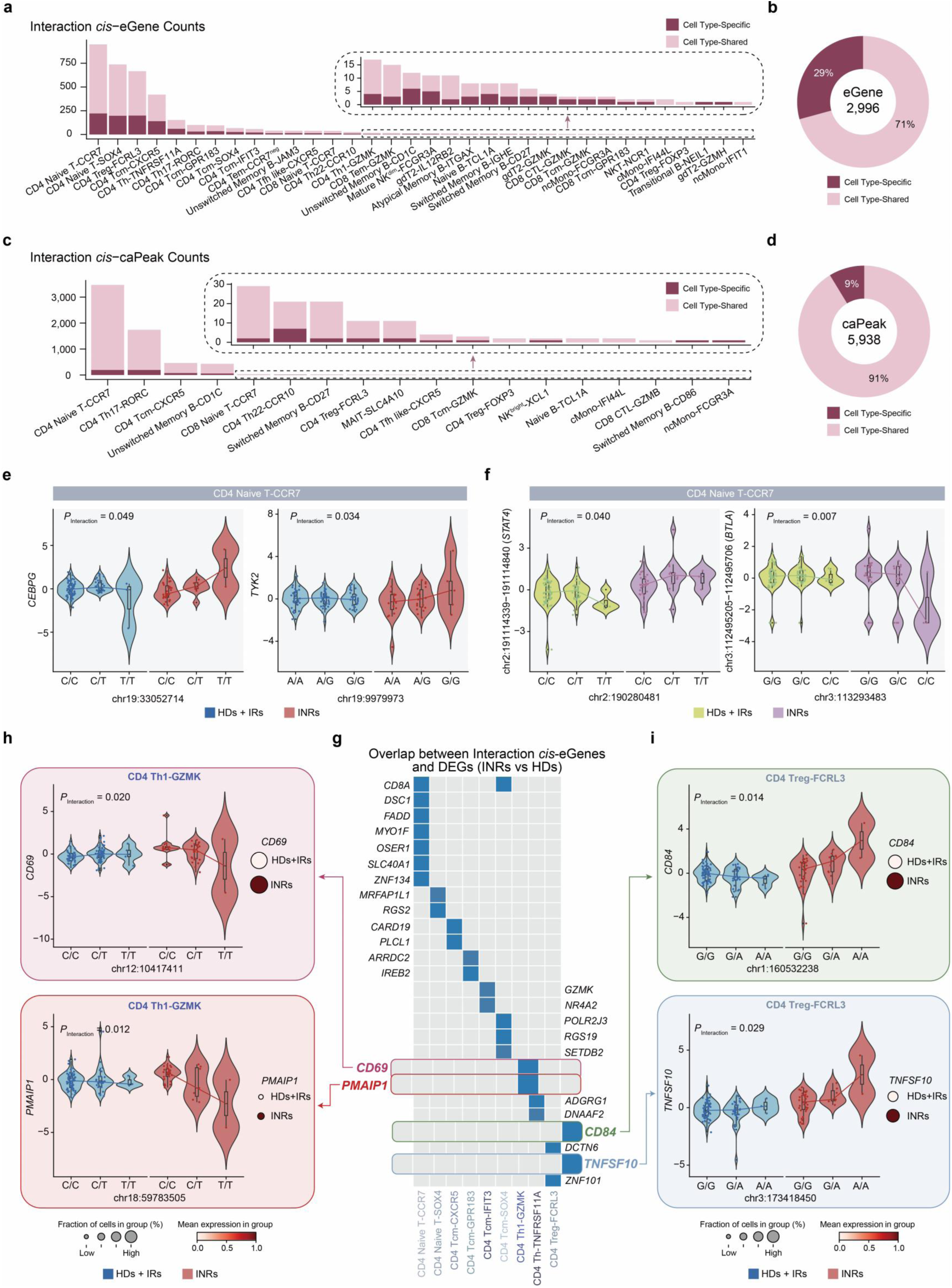
INRs-related interaction *cis*-eQTLs and *cis*-caQTLs in different immune cell types. **a**, The bar graph showing the number of cell type-shared and cell type-specific interaction *cis*-eGenes detected in the 34 cell types included in the interaction *cis*-eQTL analysis. **b**, The pie chart showing the distribution of significantly detected cell type-shared and cell type-specific interaction *cis*-eGenes. **c**, The bar graph showing the number of cell type-shared and cell type-specific interaction *cis*-caPeaks detected in the 18 cell types included in the interaction *cis*-caPeaks analysis. **d**, The pie chart showing the distribution of significantly detected cell type-shared and cell type-specific interaction *cis*-caPeaks. **e**, Violin plots showing the *cis*-eQTL effects of chr19:33052714 and chr19:9979973 on *CEBPG* and *TYK2* expression in CD4 Naive T-CCR7 in INRs and HDs+IRs control. All samples are shown as individual points. **f**, Violin plots showing the *cis*-caQTL effects of chr2:190280481 and chr3:113293483 on chr2:191114339−191114840 (*STAT4*) and chr3:112495205−112495706 (*BTLA*) accessibility in CD4 Naive T-CCR7 in INRs and HDs+IRs control. All samples are shown as individual points. **g**, Shared genes among interaction *cis*-eGenes and DEGs (INRs versus HDs). **h**, Violin plots showing the *cis*-eQTL effects of chr12:10417411 and chr18:59783505 on *CD69* and *PMAIP1* expression in CD4 Th1-GZMK in INRs and HDs+IRs control. All samples are shown as individual points. Dot plots showing the expression levels of *CD69* and *PMAIP1* in HDs+IRs and INRs. **i**, Violin plots showing the *cis*-eQTL effects of chr1:160532238 and chr3:173418450 on *CD84* and *TNFSF10* expression in CD4 Treg-FCRL3 in INRs and HDs+IRs control. All samples are shown as individual points. Dot plots showing the expression levels of *CD84* and *TNFSF10* in HDs+IRs and INRs. All samples are shown as individual points.

We identified genetic variants influencing gene expression in immune cells from INRs. In CD4 Naive T-CCR7, significant interaction eQTLs were detected in *CEBPG* and *TYK2*. Expression of both genes increased in INRs upon genotype changes, from CC to TT at *CEBPG* (chr19:33052714) and from AA to GG at *TYK2* (chr19:9979973) (Fig. 4e). *CEBPG*, a member of the C/EBP transcription factor family, has been linked to poor clinical outcomes in multiple cancers^65^. *TYK2*, involved in JAK/STAT signaling, is a known susceptibility gene for inflammatory and autoimmune diseases^66,67^.

In terms of chromatin accessibility, CD4 Naive T-CCR7 exhibited significant interaction caQTLs within the *STAT4* (chr2:190280481) and *BTLA* (chr3:113293483) genes. Accessibility at *STAT4* increased with the CC to TT genotype shift at chr2:191114339-191114840, whereas *BTLA* accessibility decreased upon GG to CC transition at chr3:112495205-112495706 (Fig. 4f). *STAT4* not only promotes inflammatory T cell responses but can also suppress Treg development by modulating the chromatin state at the Foxp3 locus^68^.

*BTLA*, an immunoregulatory receptor of the CD28 family, inhibits T and B cell activation and cytokine production^69^, and supports anti-pathogen immunity^70^.

Immune cells play a central role in the pathogenesis of various diseases, and genetic risk loci linked to autoimmune disorders may influence chromatin accessibility and gene expression in a cell type-specific manner. To explore this, we analyzed interaction effects of *cis*-xQTLs across immune cell types, with a focus on genes associated with autoimmune diseases. We identified 38 autoimmune disease-related risk genes^71–73^ across 11 immune cell types, including 19 eGenes and 44 caPeaks exhibiting INR-specific effects (Supplementary Fig. 10e). At two loci within *ETS1* (chr11:128940031 and chr11:128835058), significant caQTL interaction effects were observed in CD4 Naive T-CCR7 and CD4 Tcm-CXCR5, where chromatin accessibility increased with genotype changes from CC to GG or TT (Supplementary Fig. 10f). For *PRKCB*, an eQTL at chr16:22918168 in CD4 Tcm-CXCR5 showed elevated expression associated with the AA to GG genotype change (Supplementary Fig. 10g). At one loci within *UBE2L3* (chr22:22197535), an eQTL in Atypical Memory B-ITGAX displayed increased expression from TT to CC (Supplementary Fig. 10h).

Notably, *ETS1*, a key transcription factor in inflammatory signaling and proliferation^74^, carried a caQTL linked to inflammatory dysregulation in INRs.

The eQTL at *PRKCB*, a gene associated with severe mycobacterial infection outcomes^75^, may underpin accelerated disease progression in INRs. Furthermore, the eQTL affecting *UBE2L3*, a recognized autoimmune risk locus^71,73,76^, is associated with immune reconstitution failure in PLWH.

### Shared INRs-related *cis*-eGenes and DEGs

We investigated the relationships among *cis*-caPeaks, *cis*-eGenes, and DEGs using INR-associated *cis*-xQTL results. This analysis identified 177 genes overlapping between interaction *cis*-eGenes and interaction *cis*-caPeaks, and 27 genes shared between interaction *cis*-eGenes and DEGs (Supplementary Fig. 10i). In nine CD4^+^ T cell subtypes, 26 genes overlapped between interaction *cis*-eGenes and DEGs (Fig. 4g). DEG enrichment analysis highlighted pathways involved in T cell activation, inflammatory response, apoptosis regulation, and cytokine-mediated signaling, such as interferon-gamma (IFN-γ) signaling (Fig. 2b).

Based on these findings, we focused on four representative genes: *CD69*, involved in T cell activation^77^; the pro-apoptotic genes *PMAIP1*^78^ and *TNFSF10*^79^, and *CD84*, associated with IFN-γ signaling^80^. Significant interaction eQTL effects were detected for these genes in specific CD4^+^ T cell subsets: in CD4 Th1-GZMK for *CD69* and *PMAIP1*, and in CD4 Treg-FCRL3 for *CD84* and *TNFSF10* (Fig. 4h, i). In CD4 Th1-GZMK, a genotype shift from CC to TT correlated with reduced expression of *CD69* and *PMAIP1* (Fig. 4h). In CD4 Treg-FCRL3, genotype changes from GG to AA were associated with increased expression of *CD84* and *TNFSF10* (Fig. 4i).

In addition, significant interaction caQTLs were observed at two loci (chr21:32971387 and chr21:33897463) mapping to *IFNAR2* in CD4 Naive T-CCR7 and CD4 Th17-RORC, respectively (Supplementary Fig. 10j). In both subtypes, genotype changes (CC to AA or TT to CC) were associated with increased chromatin accessibility at two regulatory loci within *IFNAR2* (chr21:33231474-33231975 and chr21:33215493-33215994). Given the central role of *IFNAR2* in type I interferon signaling^81^, these caQTL effects may contribute to heightened interferon responses in INRs.

### sc-eQTL detects INRs-associated gene-regulatory variants

To identify genetic variants regulating CD4^+^ T cell gene expression, we mapped *cis*-eQTLs at both single-cell and bulk resolution. Following an adapted version of a previously described method^21^, we identified 3,551 *cis*-eGenes across four major CD4^+^ T cell types (Fig. 5a). To evaluate the robustness of our approach, we compared standardized effect sizes derived from bulk and single-cell eQTL analyses. As anticipated, SAIGEQTL (sc-eQTL) detected a greater number of eGenes, with shared eGenes comprising 66.5% of those identified by TensorQTL (bulk-eQTL) (Fig. 5a). Moreover, effect sizes of significant INR-associated eQTLs were highly correlated between the two platforms (*R* = 0.877; Fig. 5b). To facilitate comparative analysis, we aggregated cell types from each dataset into four CD4^+^ T cell subsets: naïve (Tn), memory (Tm), helper (Th), and regulatory (Treg) cells. Using the π₁ statistic, we quantified eQTL sharing and observed widespread concordance across subsets (Fig. 5c). Shared eQTLs generally displayed consistent effect directions, with correlation coefficients ranging from 0.895 to 0.929 (Fig. 5d).

**Fig. 5.**
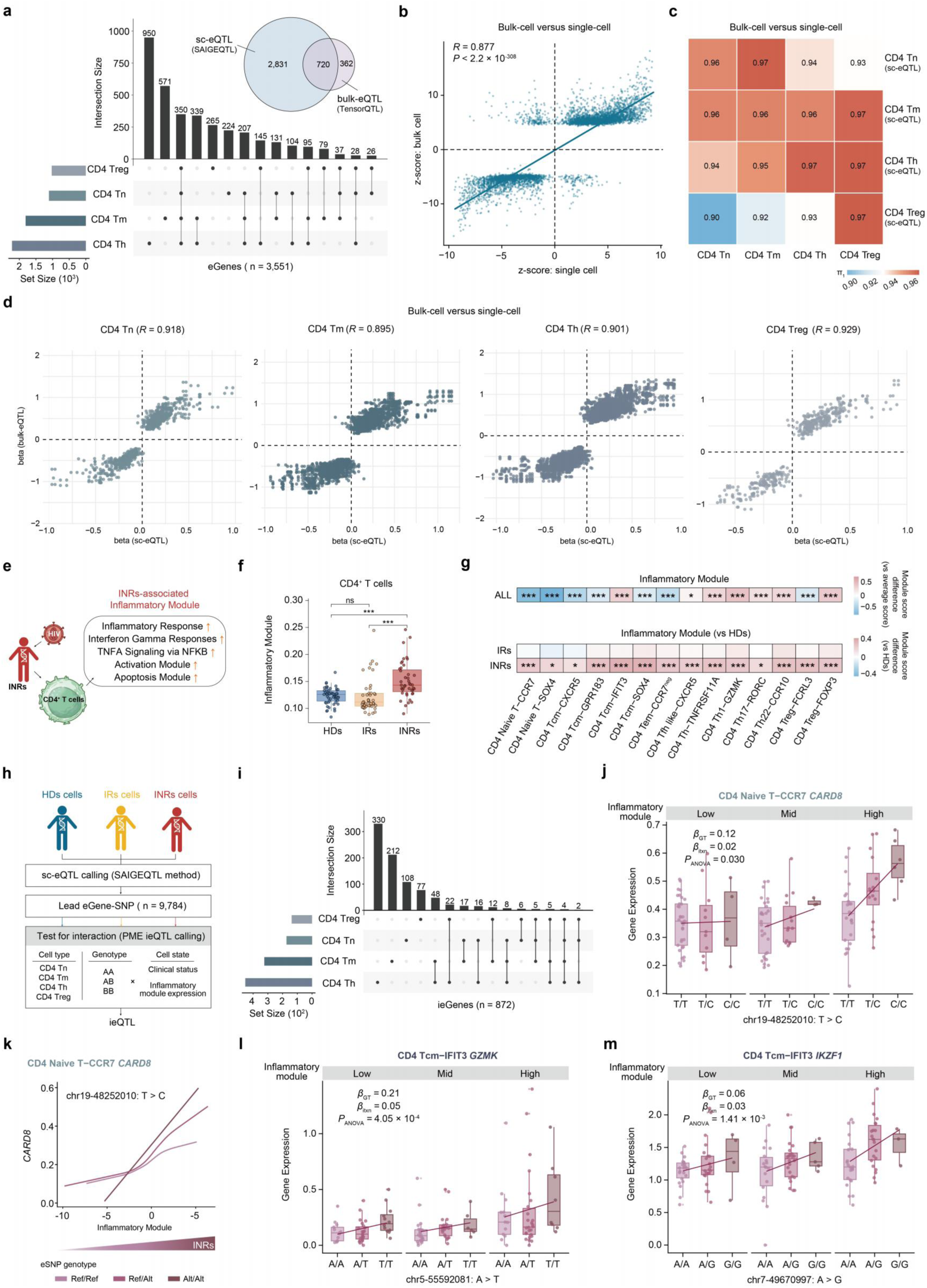
INRs-associated gene-regulatory variants detected by sc-eQTL mapping in CD4^+^ T cells. **a**, Upset plot illustrating the count of significant *cis*-eGenes across four major CD4^+^ T cell types. Venn diagram showing the number of shared *cis*-eGenes between sc-eQTL and bulk-eQTL. **b**, Replication of sc-eQTLs using bulk eQTL data. Pearson’s correlations were calculated and plotted using normalized effect sizes (z-score = β/s.e.) of significant gene-SNP pairs. Pearson’s correlation coefficient and the corresponding P values are indicated above. **c**, Heatmap showing π_1_ statistic to quantitate the extent of eQTL sharing between each pair of cell type from sc-eQTL and bulk-eQTL. **d**, The correlation of genetic effect of shared *cis*-eQTLs between sc-eQTL and bulk-eQTL. **e**, Schematic diagram of the definition of INRs-associated inflammatory module. **f**, Box plots showing the inflammatory module score in CD4^+^ T cells in HDs, IRs, and INRs. Different groups are shown in different colors. **g**, The module score of inflammatory module in CD4^+^ T cell types. Upper heatmaps depicting the difference between average scores of 14 cell types and that of all single cells. The module scores of cells in each cell type were compared with the average score of all CD4^+^ T cells. Lower heatmaps depicting the difference between average scores of IRs or INRs and those of HDs in each of 14 cell types. **h**, Scheme for identifying ieQTLs. Broad sets of sc-eQTLs were separately identified by the PME model among all cells. For significant eQTLs, interactions of genotype with cell states were tested to identify ieQTLs. **i**, Upset plot of significant ieGenes from four major CD4^+^ T cell types. **j**, Examples of CD4 Naive T−CCR7 ieQTLs in *CARD8* that exhibit significant interaction with the inferred zonal location of cells. The boxplots show gene expression according to genotype and zonal location as inferred from module expression bins. **k**, The plot shows kernel regression lines of expression by each genotype over module expression, which indicate distinct zonation patterns. **l**-**m**, Examples of CD4 Tcm-IFIT3 ieQTLs in *GZMK* (**l**) and *IKZF1* (**m**) that exhibit significant interaction with the inferred zonal location of cells. The boxplots show gene expression according to genotype and zonal location as inferred from module expression bins.

We next examined inflammatory gene modules associated with INRs. These modules were significantly enriched in five key pathways: inflammatory response, interferon-gamma response, TNFα signaling via NF-κB, immune activation, and apoptosis (Fig. 5e). Expression levels of these inflammatory modules were markedly elevated in CD4^+^ T cells from INRs compared to IRs and HDs (Fig. 5f). Across 14 CD4^+^ T cell subsets, inflammatory modules were significantly upregulated in INRs relative to HDs, whereas their expression in IRs remained comparable to that in HDs (Fig. 5g).

Leveraging the capacity of scRNA-seq to resolve cellular heterogeneity, we incorporated cell state information into our assessment of genetic variant effects on gene expression, thereby identifying ieQTLs (Fig. 5h). Cell states were quantified as module expression scores, specifically, inflammatory gene modules. To focus on variants with detectable genotype effects, we restricted interaction analyses to significant eQTLs identified across all donor cells, including those from INRs, IRs, and HDs (Fig. 5h). Among 9,784 eGene-SNP pairs detected across four CD4^+^ T cell types, 872 exhibited significant ieQTLs (Fig. 5i). A notable example was an ieQTL for *CARD8*, an inflammasome sensor of HIV-1 protease activity^82^ known to contribute to CD4^+^ T cell depletion during pathogenic infection^83^. The eQTL effect was enhanced in CD4 Naive T-CCR7 from INRs, with the highest *CARD8* expression observed in carriers of the chr19:48252010 CC genotype (Fig. 5j, k). Two additional INR-specific eQTLs were identified: one in *GZMK*, which promotes inflammatory responses^84^, and another in *IKZF1*, a recently described transcriptional repressor of HIV-1^85^. In CD4 Tcm-IFIT3, *GZMK* expression increased in INRs carrying the TT genotype, whereas *IKZF1* expression was elevated in INRs with the GG genotype (Fig. 5l, m).

### Upstream regulators of ieQTLs provide mechanistic insights

A primary mechanism by which eQTLs influence gene expression is through altering the binding affinity of upstream TFs for regulatory regions^20^. We therefore hypothesized that certain cell-state-specific ieQTLs are orchestrated by ‘master’ upstream regulators with functional relevance to disease progression (Fig. 6a). Using SCENIC+ on single-cell data, we inferred TF activity and examined whether ieSNPs (SNPs underlying ieQTLs) create or disrupt TF-binding motifs, thereby providing a mechanistic link to genotype-dependent effects (Fig. 6b). Among candidate transcriptional targets, *OAS3* expression and the rs2701615 genotype exhibited a significant interaction effect specific to INRs (Fig. 6c).

**Fig. 6.**
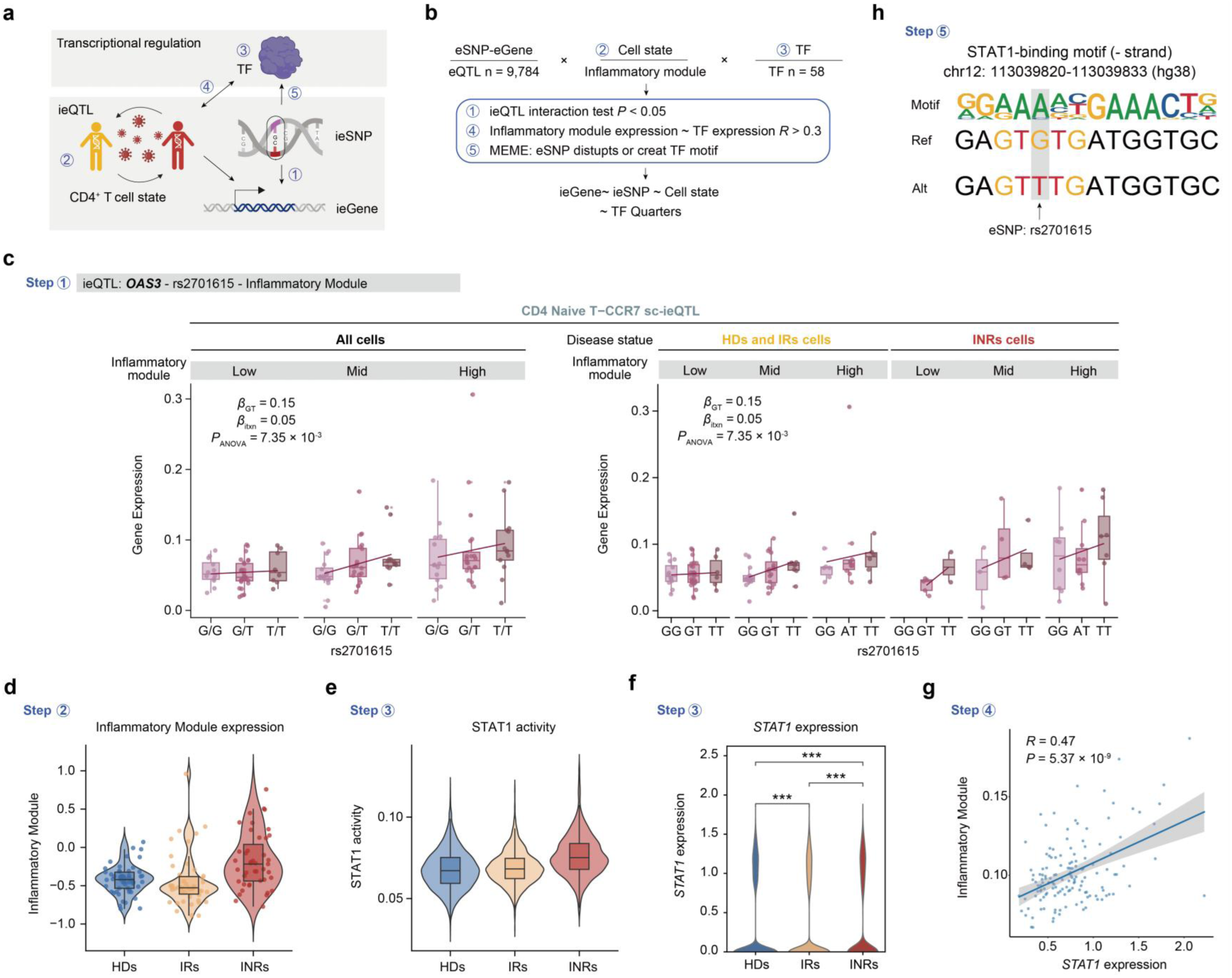
Integrative analysis of ieQTLs and upstream regulators identifies genotype-specific regulatory variants in INRs. **a**, Schematic illustration of the interrelationship of upstream TFs, ieQTL components (cell states, ieSNPs and ieGenes). **b**, Scheme of the analysis pipeline, where each step has parameters for the priority quartet (TF, cell state, ieSNP, and ieGenes). **c**-**h**, Example of an INRs-associated ieQTL in *OAS3* and its upstream regulator STAT1, displaying each step in the analysis pipeline described in (**b**). **c**, The eQTL plot for an ieQTL in *OAS3* plotted against genotype of rs2701615 (chr12:113039824:A>T) in cells derived from all donors (left), HDs and IRs (middle), and INRs (right). Regression coefficients (β_GT_ and β_itxn_ indicate PME coefficients for the genotype and interaction term, respectively) and P values (P_ANOVA_ was obtained from the interaction test) are displayed below. **d**, Expression of inflammatory module across disease stages. **e**, Violin plot of STAT1 activity across disease stages. **f**, Violin plot of *STAT1* expression across disease stages. **g**, Two-dimensional histogram showing correlation of inflammatory module expression with *STAT1* expression in CD4 Naïve-CCR7. **h**, Alignment of STAT1 motif (sequence logo) with eSNP sequences.

Inflammatory module expression, along with STAT1 activity and expression levels, were elevated in CD4^+^ T cells from INRs (Fig. 6d-f). Inflammatory module scores correlated positively with *STAT1* expression (*R* = 0.47; Fig. 6g). Notably, the rs2701615:A allele was found to generate a novel STAT1-binding motif (P < 0.01 from the FlMO software, Fig. 6h), suggesting enhanced STAT1 binding at this locus, which in turn promotes *OAS3* expression possibly.

#### Dissecting HIV immune status with single cell multi-omics deep learning

Characterizing the profound immune heterogeneity in people living with HIV and identifying its underlying *cis*-regulatory drivers remains a significant challenge. To address this, we developed scPRISM, a multi-modal deep learning framework that leveraged scRNA-seq and scATAC-seq data to link gene expression with chromatin state (Fig. 7a). The model’s inputs were defined by curating 582 differentially expressed genes from the transcriptome and profiling 5,583 associated chromatin-accessible peaks from the epigenome, located within ±250 kb window flanking these gene bodies (Fig. 7b). This resulted in a median of 6 peaks linked per gene (ranging from 1 to 93; Supplementary Fig. 11a). To align cellular profiles across unmatched scRNA-seq and scATAC-seq datasets, we established a direct cell-to-cell correspondence through an iterative minimum-weight matching algorithm framed as a bipartite-graph optimization problem (Supplementary Fig. 11b, c). The resulting cross-modally paired dataset serves as the input for two dedicated analytical modules, including (1) scPRISM-CLIN, which classifies cells into clinical phenotypes (INRs, IRs, and HDs) using the integrated profiles, and (2) scPRISM-EXPR, which predicts gene expression directly from the chromatin accessibility landscape of each paired cell (Supplementary Fig. 12a-e).

**Fig. 7.**
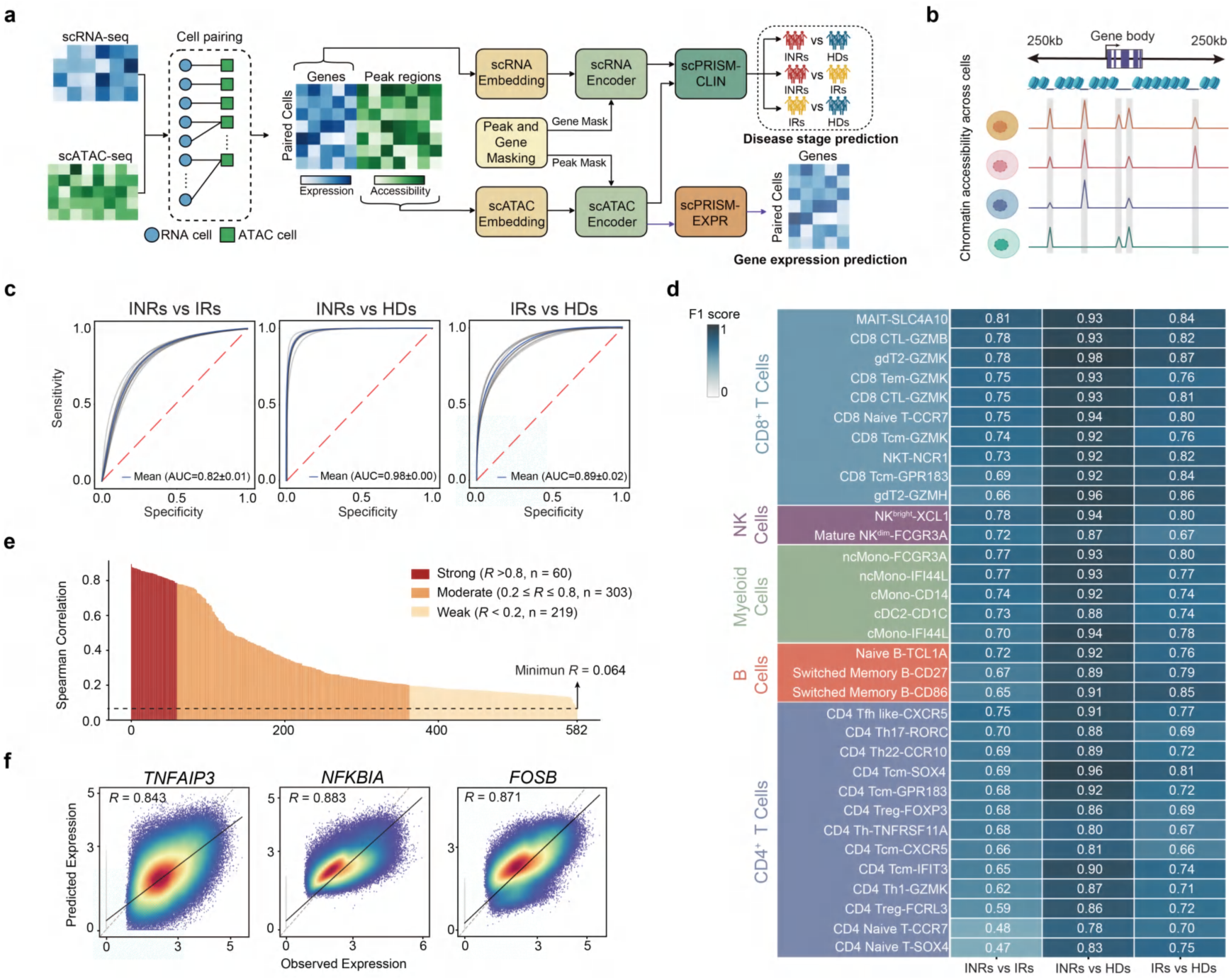
The scPRISM framework enables dual prediction of disease states and gene expression. **a**, Schematic of the scPRISM architecture, detailing the scPRISM-CLIN module for disease state classification from integrated multi-omics profiles, and the scPRISM-EXPR module for predicting gene expression from chromatin accessibility. **b**, Strategy for selecting chromatin-accessible peaks within a ±250 kb window of target gene bodies. **c**, Performance of the scPRISM-CLIN module shown by ten-fold cross-validation receiver operating characteristic (ROC) curves. Average ROC curves are highlighted, with mean area under the curve (AUC) values indicated. **d**, Pairwise classification performance (F1 scores) of the scPRISM-CLIN module across all L3-level cell subtypes. **e**, Spearman correlation coefficients between observed and scPRISM-EXPR-predicted expression for 582 genes. Genes are ranked by correlation and colored by prediction tier (strong, moderate, or weak). All correlations are significant (P < 2.2×10^-308^). **f**, Observed versus predicted expression for three representative genes (*TNFAIP3*, *NFKBIA*, and *FOSB*). Spearman correlation is shown (P < 2.2×10^-308^).

On the held-out test dataset, scPRISM demonstrated robust, dual predictive capabilities. The scPRISM-CLIN demonstrated high-fidelity discrimination of disease states, with mean AUCs of 0.82 (INRs vs IRs), 0.98 (INRs vs HDs), and 0.89 (IRs vs HDs) (Fig. 7C), and strong patient-level accuracy (Supplementary Fig. 13a). The superior discriminative performance across pairwise comparisons highlights the profound immunological divergence between these groups. This discriminative power varied by cell type (Fig. 7d and Supplementary Fig. 13b). Notably, MAIT-SLC4A10 and CD8 CTL-GZMB, previously implicated in defective immune reconstitution and mitochondrial dysfunction in INRs^10^, were highly informative for distinguishing INRs from IRs (F1 scores: 0.82 and 0.78, respectively), compared to other L2 populations. Concurrently, the scPRISM-EXPR module accurately predicted gene expression from chromatin accessibility, showing a significant global Spearman correlation between observed and predicted values (*P* < 0.05; Fig. 7e). Genes were stratified into strong (*R* > 0.8; n = 60), moderate (0.2 ≤ *R* ≤ 0.8; n = 303), and weak (*R* < 0.2; n = 219) prediction tiers. Strong-tier genes were associated with significantly higher accessible peak density compared to other tiers (*P* < 0.001; Supplementary Fig. 13c), indicating that chromatin accessibility enhances predictive accuracy. Representative genes including *NFKBIA*, *TNFAIP3*, and *FOSB* exhibited correlations exceeding 0.80 (Fig. 7f).

### scPRISM reveals cell type-specific gene signatures and regulatory circuits driving immune dysregulation

To pinpoint the cellular and molecular drivers of immune reconstitution failure, we interrogated our scPRISM-CLIN model using Integrated Gradients (IG) to quantify the contribution of 32 immune genes of interest (Fig. 8a). This attribution analysis revealed a key distinction in the immune signatures of patient cohorts. While both IRs and INRs showed broad transcriptional divergence from HDs, the critical classification separating INRs from IRs was traced to a highly specific and subtle signature (Fig. 8b). The model’s ability to distinguish these two clinical outcomes was not driven by global changes, but primarily by gene expression patterns within NK cells and specific myeloid subsets. Among the most influential features were key NF-κB pathway members (*NFKBIA*, *TNFAIP3*) and immediate-early genes (*FOSB*).

**Fig. 8.**
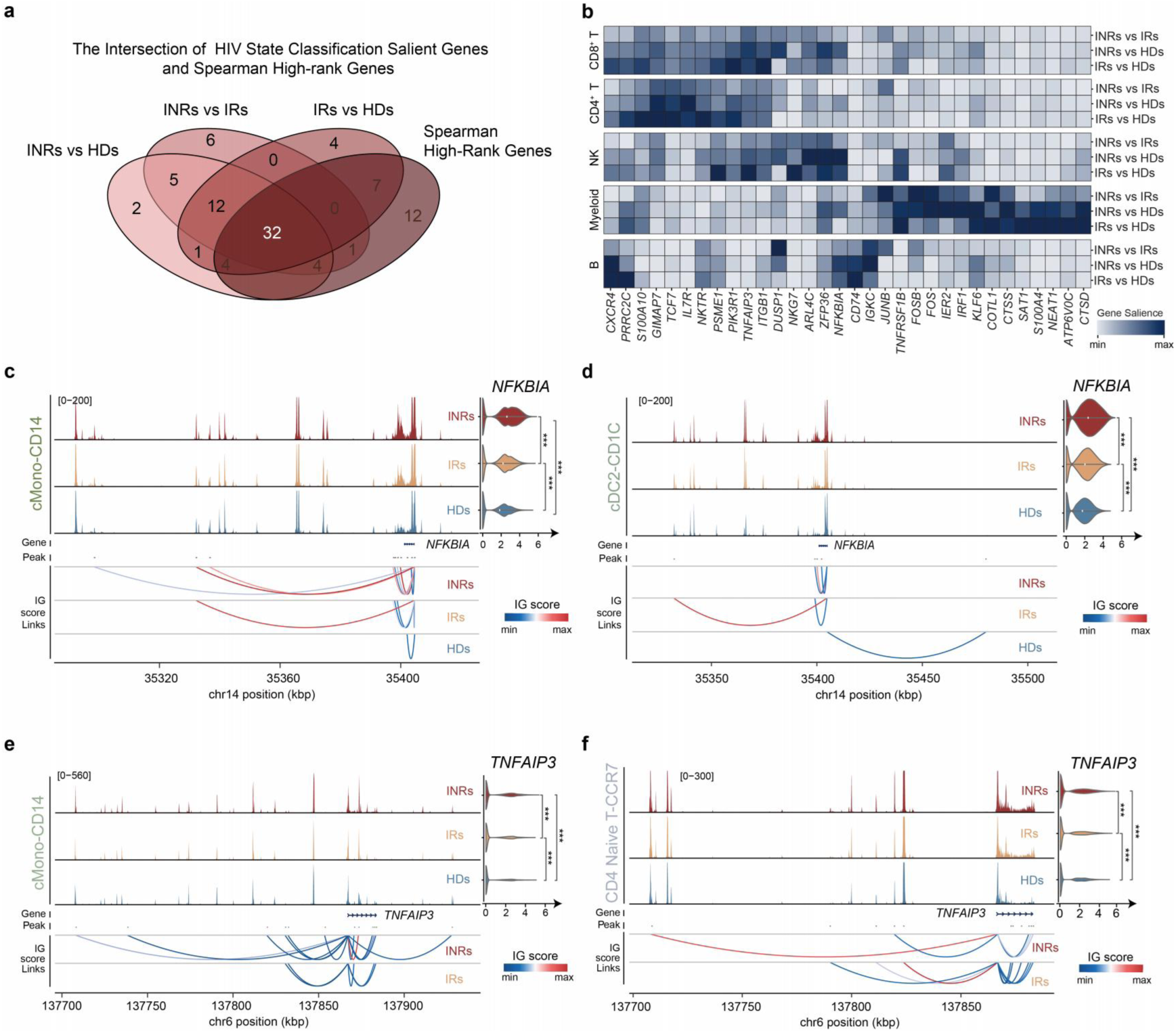
Interpretable scPRISM resolves cell type-specific gene signatures and regulatory peak-to-gene links. **a**, Venn diagram showing the overlap between the 60 genes most predictive of HIV disease state (identified by scPRISM-CLIN) and the 60 genes whose expressions are most accurately predicted from chromatin accessibility (Spearman’s R > 0.8, identified by scPRISM-EXPR). **b**, Heatmap of Integrated Gradient (IG) scores for 32 immune genes, showing their contribution to disease state classification across major (L1) cell types. **c**-**d**, Cell type-specific chromatin peaks contacting *NFKBIA* in cMono-CD14 for (**c**) and cDC2-CD1C for (**d**). **e**-**f**, Cell type-specific chromatin peaks contacting *TNFAIP3*, as identified by scPRISM-EXPR, in cMono-CD14 for (**e**) and CD4 Naive T-CCR7 for (**f**). For panels (**c**-**f**), arc diagrams illustrate significant peak-to-gene links. Violin plots show gene expression, with inset boxplots where the center line is the median, box limits are the 25th-75th percentiles, and whiskers are 1.5× the interquartile range. Significance was determined by a two-sided Mann-Whitney U-test, ***P < 0.001, **P < 0.01, and *P < 0.05.

We next leveraged scPRISM’s interpretive power to identify *cis*-regulatory interactions in a cell-subset-resolved manner. By applying the IG algorithm to the scPRISM-EXPR module, we quantified the contribution of individual chromatin peaks to the expression of the 60 genes in the strong prediction tier (Supplementary Fig. 14a). We focused on 692 chromatin-accessible regions within ±250 kb of these genes, comprising promoters (38.70%), distal enhancers (40.87%), and proximal enhancers (19.86%) (Supplementary Fig. 14b). We observed that salient peaks (IG scores > 0.005) exhibited significantly greater evolutionary conservation, validating their functional potential (Supplementary Fig. 14c). IG scoring of peak-gene interactions revealed that clinical states are defined by highly cell type-specific *cis*-regulatory circuits.

We found that the same gene could be regulated by different chromatin contacts depending on the cellular context. For example, *NFKBIA* highly expressed in INRs was associated with a greater number of long-range contacts in cMono-CD14 but fewer significant peak-gene links in cDC2-CD1C compared to other clinical groups (*P* < 0.05; Fig. 8c, d). Our analysis also identified a potentially relevant regulatory interaction with possible mechanistic implications for immune failure. We detected a chromatin peak within the promoter of *PSMA6* that showed strong predictive correlation with *NFKBIA* expression, specifically in the T and NK cell subtypes of INRs (Supplementary Fig. 14d). This observation may offer a molecular explanation for the non-responder phenotype, given that *PSMA6*-mediated proteasomal degradation of IκBα (the protein encoded by *NFKBIA*) is known to be essential for NF-κB activation. Similarly, *TNFAIP3* was upregulated in INRs in both cMono-CD14 and CD4 Naive T-CCR7 (Fig. 8e, f). However, this was driven by a dense cluster of enhancer loops in monocytes, whereas T-cells utilized a sparser set of contacts (*P* < 0.05). These findings, also observed for the transcription factor *FOSB* (Supplementary Fig. 14e, f), demonstrated that cell type-specific reliance on distinct regulatory circuits was a key feature of the immune dysregulation seen in INRs.

## Discussion

In this study, we present a comprehensive single-cell multi-omics landscape of INRs, IRs, and HDs. Our analysis reveals markedly elevated inflammatory responses, cellular activation, and apoptosis in INRs, mediated through a ligand-receptor driven multicellular signaling network. These dysregulated genes and pathways are likely contributors to impaired immune recovery in INRs. To decipher the host genetic basis of immune reconstitution failure, we integrated WGS data to map *cis*-eQTLs and *cis*-caQTLs across diverse immune cell types. Furthermore, we developed a single-cell eQLT-based analytical framework that identifies functional units comprising TFs, cellular states, ieSNPs, and ieGenes. While prior models have predicted immune reconstitution failure using clinical parameters such as age, baseline CD4^+^ T cell count, and CD4^+^/CD8^+^ T cell ratios^4,86–88^, our study advances the field by introducing two multi-omics-based predictive tools: scPRISM-CLIN for distinguishing disease states, and scPRISM-EXPR for inferring gene expression from chromatin accessibility.

We observed a reduction in CD4^+^ T cell and B cell subsets in INRs compared to IRs, alongside an expansion of inflammation-associated NK and monocyte populations. Previous single-cell studies reported a significant decrease in MAIT in INRs. In contrast, our data indicate that the proportion of MAIT marked by *SLC4A10* expression was comparable between INRs and IRs^10^. Consistent with Liu et al., we find that T follicular helper (Tfh) cell dysfunction likely undermines memory B cell activity in INRs, with Tfh subsets emerging as potential predictors of poor immune recovery^89^. Concurrently, expanded NK cell populations exhibiting heightened activation may contribute to sustained inflammation-driven immune impairment^90^.

Despite effective suppression of HIV, a state of chronic low-grade inflammation persists^39^. This sustained inflammatory environment promotes prolonged overexpression of ISGs via the NF-κB and STAT signaling pathways, leading to long-term immune dysregulation. We observed markedly elevated expression of *STAT1* and ISGs across CD4^+^ T cell subsets, collectively forming a distinct interferon-high phenotype. These findings confirm that INRs exhibit heightened responsiveness to interferons^40^. In line with this, Liu et al. demonstrated that the enhanced IFN response signature in CD4^+^ T cells of INRs strongly correlates with T cell exhaustion markers^35^. Furthermore, to counterbalance NF-κB pathway overactivation, CD4^+^ T cells appear to upregulate the negative regulators *NFKBIA* and *TNFAIP3*, thereby attenuating persistent inflammatory signaling^91^. Consistent with this mechanism, we detected significantly elevated levels of *NFKBIA* and *TNFAIP3* in CD4^+^ T cells from INRs compared to IRs and HDs. We propose that persistent inflammation may accelerate T cell aging, and aged T cells likely exhibit diminished functional capacity^92^, ultimately contributing to the sustained impairment of CD4^+^ T cell recovery.

TFs often recruit chromatin remodeling complexes to promoters and enhancers. The re-opening of chromatin upon cellular stimulation reflects the activity levels of sequence-specific TFs^93^. These functional genomic elements modulate gene expression through TF-DNA interactions. To systematically decipher transcriptional regulation, we constructed an enhancer-driven GRN by leveraging chromatin accessibility dynamics to infer TF activity. Most TFs, including the AP-1 family components FOS and JUN, which are central to inflammatory signaling and stress responses, as well as IRF and STAT family members, exhibited enhanced regulon activity in INRs. Target genes of FOS and JUN were enriched in pro-inflammatory pathways^94,95^, whereas those of IRF9, a key regulator of type I interferon signaling, and STAT1 were associated with interferon response and chronic inflammation^96–98^. Collectively, these findings indicate that key transcription factors are hyperactivated in PBMCs from INRs, where they orchestrate a pro-inflammatory and interferon-driven transcriptional program that underlies INR pathology.

Intercellular communication analysis revealed strengthened signaling through multiple ligand-receptor pairs, including IFNG-IFNGR1/2, TNF-TNFRSF1A/B, IL18-IL18R1, CCL3-CCR1/4/5, and CXCL2-CXCR2 across myeloid, CD4^+^ T, CD8^+^ T, NK, and B cells in INRs. In parallel, weakened CD86-CTLA4 inhibitory interactions may contribute to disrupted T-cell homeostasis. These findings indicate that immune reconstitution failure stems from aberrant cytokine and chemokine signaling alongside sustained inflammation. This interpretation is reinforced by the well-documented role of IFN-γ (encoded by *IFNG*) in promoting inflammatory and immune-mediated diseases^53^, as well as the recognized involvement of chemokine signaling dysregulation in infectious and inflammatory pathologies^99^.

We next investigated how genetic variation influences immune characteristics in INRs. Correlation analyses of genetic variants with gene expression and chromatin accessibility across individual cell types indicated that most interaction *cis*-eQTLs and *cis*-caQTLs exert cell type-shared effects. Importantly, we identified loci exhibiting selective effects among INRs, IRs, and HDs that have not been previously reported. This aligns with prior evidence linking chemokine receptor variants to immune recovery status in PLWH^100^. We observed that INR-associated variants in *PRKCB*, *TYK2*, and *UBE2L3* exhibited CD4^+^ T cell-specific eQTL effects. Additionally, *IFNAR2*, *STAT4*, and *BTLA* variants displayed CD4^+^ T cell-specific caQTL effects in INRs. *PRKCB* alterations have been linked to progression of T-cell leukemia/lymphoma^101^, and elevated *PRKCB* expression correlates with accelerated HIV-1 disease progression^72^. In our data, CD4 Tcm-CXCR5 showed a significant eQTL effect on *PRKCB* in INRs, leading to its increased expression. *UBE2L3* is known to promote NF-κB activation via the linear ubiquitin chain assembly complex downstream of TLR7 in SLE. Its overexpression elevates *NFKBIA* and *IFNB1* mRNA under baseline conditions^102^. Consistently, we observed significantly increased *UBE2L3* expression in Atypical Memory B-ITGAX in INRs upon SNP alteration. Type I interferonopathies, often resembling SLE, involve mutations affecting type I interferon signaling^103^. Both *TYK2* and *IFNAR2* are central to this pathway^81^. *IFNAR2* variants exert monocyte-specific eQTL effects in COVID-19 patients^104^, and *TYK2* polymorphisms are differentially associated with autoimmune diseases^105^. In INRs, CD4 Naive T-CCR7 exhibited eQTL effects on *TYK2* and caQTL effects at the *IFNAR2* locus, increasing their expression or chromatin accessibility, potentially elevating the risk of dysregulated interferon responses. Overactivation of type I IFN signaling may promote viral persistence and impair immune reconstitution. Genome-wide association studies have also linked *STAT4* polymorphisms to multiple autoimmune diseases^106,107^. In SLE patients, the *STAT4* risk allele rs7574865[T] elevates *STAT4* expression and pSTAT4 levels in response to IL-12 and IFN-α, amplifying IL-12 induced IFN-γ production in T cells^108^. Accordingly, we found that CD4 Naive T-CCR7 in INRs exhibit a caQTL effect at the *STAT4* locus, increasing its chromatin accessibility. These results suggest that *STAT4* locus variants may exacerbate inflammatory responses in INRs by enhancing IL-12-dependent IFN-γ production.

Several studies have implicated *CARD8* in inflammasome activation and the loss of CD4^+^ T cells following HIV-1 infection^82,83,109^. Additional evidence indicates that *CARD8* positively regulates inflammation and apoptosis^110^, and the WBC count exhibits an increasing trend across the three *CARD8* genotypes (AA < AT < TT)^111^. In our data, *CARD8* expression was upregulated in CD4 Naïve T-CCR7 from INRs carrying the CC genotype. These results suggest that the C allele at chr19:48252010 may contribute to more severe inflammatory responses, increased apoptosis, and lower CD4^+^ T-cell counts by elevating *CARD8* expression in INRs.

Furthermore, we propose that rs2701615 increases INR risk through allele-specific binding of STAT1, which in turn modulates *OAS3* expression. Previous work has shown that IFN-β1b-induced phosphorylation of STAT1 initiates *OAS3* transcription via direct binding to its promoter^112^. Liu et al. reported that sustained interferon-STAT1 signaling is closely associated with inflammatory autoimmune disorders^113^. Moreover, genes positively correlated with *OAS3* expression are linked to inflammatory signatures in PLWH^114^. Our study further demonstrated elevated inflammatory module expression and increased STAT1 activity and expression in CD4^+^ T cells from INRs, with inflammatory module expression correlating with STAT1 activity. Collectively, these findings suggest that elevated *OAS3* expression drives inflammatory pathology and increases INR risk in PLWH, particularly among rs2701615:TT carriers.

A fundamental challenge in immunology lies in bridging the landscape of chromatin accessibility with the gene expression programs that determine cellular identity and disease pathology. Our work presents scPRISM as a powerful computational tool designed to address this challenge in the context of HIV infection, advancing beyond generalized descriptions of immune dysregulation to uncover its precise cellular and molecular origins. We demonstrate that the key distinction between INRs and IRs is not a systemic immune alteration but arises from subtle transcriptional differences within specific cell subsets, particularly NK and myeloid cells. Importantly, scPRISM offers a mechanistic perspective for interpreting these disparities. By directly coupling chromatin state with transcriptional output, we reveal that the dysregulated expression of genes such as *NFKBIA* and *TNFAIP3* in monocytes and T cells of INRs is driven by a reconfigured network of enhancers and promoters. This finding underscores the limitations of transcriptomic-only analyses and emphasizes the critical need for epigenomic integration to uncover underlying mechanisms. The identification of a putative regulatory axis connecting *PSMA6* to *NFKBIA*, specifically active in T and NK cells of INRs, illustrates this capacity, offering a concrete, testable hypothesis for defective immune signaling. Ultimately, scPRISM establishes a robust framework for multi-omics integration, capable of translating complex single-cell datasets into precise, cell type-resolved hypotheses of gene regulatory dysfunction in human disease.

Our study has several limitations that warrant consideration. First, the single-cell multi-omics analysis was limited to PBMCs and conducted as a single-center cross-sectional study, which lacks the temporal dimension required to completely capture dynamic changes in genetic variants, gene expression, and chromatin accessibility over time. Second, the exclusive inclusion of male participants may introduce selection bias and limit extrapolation of findings to the broader INRs population. Future investigations with enhanced coverage, including multiple-center designs, greater representation of female participants, and longitudinal follow-up would improve the accuracy and generalizability of our findings.

In summary, this study offers a comprehensive analysis of immune reconstitution failure in PLWH. By integrating multi-omics approaches, we have identified key molecular characteristics underlying this condition, delineating an integrative landscape of transcriptional immune profiles, chromatin epigenetic states, and genetic influences in immune cells from INRs. Our findings highlight that immune cell inflammation, monocyte overactivation, and CD4^+^ T cell deficiency are likely contributors to immune reconstitution failure. These insights establish a theoretical foundation for developing T cell-based immunotherapies for PLWH. Furthermore, we demonstrate how specific genetic variants modulate the expression of immune reconstitution failure-related genes in a context- and genotype-dependent fashion. Further dissection of the genetic basis of immune cell states holds significant potential for stratifying PLWH and advancing precision therapeutics. Future studies should focus on validating these multi-omics features and exploring therapeutic approaches that mitigate T cell damage while fostering immune recovery.

## Methods

### Human subjects

The human subjects of this study were healthy volunteers and HIV-infected patients who received physical examinations at Xixi Hospital in Hangzhou, Zhejiang Province, China from March to June 2023. According to the definition of the Chinese guidelines for the diagnosis and treatment of human immunodeficiency virus infection/acquired immunodeficiency syndrome (2024 edition)^115^, after AIDS patients receive effective ART, some patients with good viral control cannot recover their CD4^+^ T lymphocyte counts, and are called immune non-responders (INRs). The diagnostic criteria for INRs used in this study are: receiving ART for more than 4 years, peripheral blood viral load is below the detection limit (< 50 copies/mL) for more than 3 years, and CD4^+^ T lymphocyte counts are still continuously below 350 cells/μL^35^. Based on this, we recruited 143 subjects, all male, aged from 26-76 years old, with a 1:1 age match between groups, who had not been vaccinated and had no viral infection within six months before enrollment. Among them, there were 53 healthy donors (HDs), 47 immune responders (IRs), and 43 INRs.

The studies involving human participants were reviewed and approved by the Clinical Research Ethics Committee of the Hangzhou Xixi Hospital (ethical approval No. LKYJ-2024-030) and the Institutional Review Board of BGI in accordance (ethical approval No. BGI-IRB 23041) with the tenets of the Declaration of Helsinki. Written informed consent was acquired from each participant. For additional details, see Supplementary table 1.

### Blood sample collection and processing

Peripheral blood (5-10 mL) was collected from each volunteer using EDTA anticoagulant tubes and processed within 4 hours. The blood samples were then processed using SepMate tubes (Stemcell) to rapidly isolate peripheral blood mononuclear cells (PBMCs). Isolated PBMCs were subjected to stepwise cryopreservation and stored in liquid nitrogen. Only PBMCs with post-thaw viability exceeding 80% were used for single-cell library preparation and sequencing. All samples were processed to DNA/cDNA in the P2 laboratory at Hangzhou Xixi Hospital and then transported to the P2 laboratory at BGI Research Hangzhou for subsequent library construction and sequencing.

### Generation of the genotype data from WGS

Genotyping was conducted using DNA extracted from PBMCs samples. Each sample was first subjected to single-cell droplet generation, after which the remaining cell pellets were stored in a -80°C freezer. Once all samples had completed the single-cell experiment, we used the MGIEasy Magnetic Beads Genomic DNA Extraction Kit (MGI) to extract genomic DNA. Subsequently, whole-genome library construction and sequencing (WGS) were carried out, with an average sequencing depth of 30×.

### Single-cell RNA and ATAC library preparation and sequencing

PBMCs were processed using the DNBelab C Series Single-Cell Library Preparation platform with the High-throughput Single-cell RNA Library Prep Set V2.0 (MGI, 940-000519-00) and ATAC Library Prep Set V1.0 (MGI, 940-000793-00). For scRNA-seq, approximately 22,000 cells were loaded per reaction, yielding over 10,000 captured cells. Following droplet generation, emulsion breakage, bead collection, reverse transcription, and cDNA amplification, the resulting scRNA-seq libraries were sequenced on DNBSEQ-T20 and DNBSEQ-T1 platforms at the China National GeneBank (Shenzhen, China). Sequencing was performed with 30 bp for cell barcodes and UMIs (Read 1), and 100 bp for cDNA along with a 10 bp sample index (Read 2).

For scATAC-seq, 100,000-300,000 remaining PBMCs were fixed in 0.01% formaldehyde, lysed, and subjected to nuclear transposition. Libraries were then constructed using the DNBelab C Series ATAC kit and sequenced on the DNBSEQ-T20 platform. The sequencing parameters included 70 bp (Read 1) for barcodes and insertion sites, and 50 bp (Read 2) for insertion sites plus a 10 bp sample index.

### Single-cell RNA-seq raw data preprocessing

Raw reads from oligo libraries (DNBSEQ-T1) and cDNA libraries (DNBSEQ-T20) were first processed using PISA (v0.12b)^116^ for error correction and filtering. Cleaned reads were then aligned to the human reference genome (GRCh38) using STAR (v2.7.2b)^117^, retaining only exon-mapped reads. Valid cell barcodes were identified using EmptyDrops^118^, and beads with similar oligo composition (based on cosine similarity) were merged to represent single cells. A gene-by-cell UMI count matrix was subsequently generated using the PISA Count module.

### Single-cell RNA-seq data analysis

The scRNA-seq data were processed using a unified computational pipeline incorporating several established tools, including Scrublet (v0.2.3)^119^, Scanpy (v1.9.3)^120^, rapids-singlecell (v0.9.1)^121^, and COSG (v1.0.1)^122^. This standardized workflow was applied across all scRNA-seq datasets in the cohort and comprised the following key steps:

#### Doublet detected

Doublets were identified using Scrublet (v0.2.3)^119^. For each library, raw count data were loaded into AnnData objects, and Scrublet simulated artificial doublets to compute doublet scores for each cell using a k-nearest neighbor classifier. The analysis was performed with the following parameters: expected_doublet_rate = 0.08, min_counts = 2, min_cells = 3, log_transform = True, min_gene_variability_pctl = 85 and n_prin_comps = 30. Cells with scores above 0.2 (via the call_doublets function) were flagged as potential doublets and annotated in the metadata for downstream filtering.

#### Quality control at single cell level

Quality control (QC) was performed using Scanpy (v1.9.3)^120^. For each cell, QC metrics including total UMI counts, number of detected genes, and the proportions of mitochondrial, ribosomal, and erythrocyte gene expression were calculated. Doublets identified in the previous step were excluded. Cells were retained if they met the following thresholds: 500-3,000 detected genes, 1,000-10,000 total counts, < 10% mitochondrial gene content, < 1% erythrocyte gene content, and > 1% ribosomal gene content. Additionally, genes expressed in fewer than three cells, as well as all mitochondrial, noncoding RNA, ribosomal, and erythrocyte genes, were excluded from downstream analyses.

#### Batch effect removal, dimensionality reduction, cell clustering and reclustering

Batch correction, dimensionality reduction, and clustering were performed using the Scanpy workflow (v1.9.3). First, we identified the top 2,000 highly variable genes (HVGs) using sc.pp.highly_variable_genes with the “seurat_v3” flavor. The raw count matrix was preserved in .layers[‘count’] for future use. Data were then normalized using sc.pp.normalize_total (target_sum = 1e4), log-transformed with sc.pp.log1p, and scaled (HVGs only) using sc.pp.scale with zero_center = False. Principal component analysis (PCA) was applied via sc.tl.pca, and batch effects were corrected using Harmony (sc.external.pp.harmony_integrate, batch = ‘batch’). Neighbor graph construction was based on the first 21 Harmony components (sc.pp.neighbors), followed by UMAP embedding (sc.tl.umap, min_dist = 0.3) and clustering with the Leiden algorithm (sc.tl.leiden, n_iterations = 2). DEGs were identified using COSG. Doublet-enriched clusters were removed, and distinct B, T, and myeloid populations were isolated for further sub-clustering. For reclustering, we restarted from the raw counts stored in.layers[‘count’]. After log-normalization, HVGs were re-identified using “seurat_v2” flavor (sc.pp.highly_variable_genes, batch_key = ‘stage’), followed by scaling and PCA. Batch correction was performed with the rapids-singlecell Harmony implementation (rsc.tl.harmony_integrate, batch = ‘experimental_date’). Downstream steps for dimensionality reduction and clustering remained the same. Doublet-enriched clusters and mixed populations were removed iteratively, and specific subsets were extracted using marker-based selection for further re-clustering.

In total, 2,744,009 high-quality single-cell transcriptomes were obtained, which were grouped into five major cell types: CD4⁺ T cells (645,339 cells), CD8⁺ T and unconventional T cells (1,261,644 cells), NK cells (204,859 cells), B cells (278,699 cells), and myeloid cells (353,468 cells).

#### Manual annotation of circulating immune cells

For the 2,744,009 high-quality PBMCs retained after quality control, we performed detailed cell type annotation based on canonical marker genes and DEGs between clusters. Cell type identification was guided by clustering patterns and marker gene expression, following established methodologies and previous studies^123,124^. Marker genes were selected based on prior knowledge, public databases, and relevant literature. Supplementary table 2 summarizes the key positive and negative marker genes used to define immune cell types at multiple levels. Additionally, we incorporated marker genes identified through COSG-based differential expression analysis within each fine-resolution cluster to support annotation. The most abundant population consisted of *CD3D*-expressing T cells, which were further subdivided into CD4⁺ T cells, CD8⁺ T cells, and other unconventional T cells. Iterative clustering was performed until CD4⁺ and CD8⁺

T cells were fully separated based on *CD4*, *CD8A*, and *CD8B* expression. Unconventional T cells such as NKT, gdT, and MAIT cells exhibited greater transcriptional similarity to CD8⁺ T cells and were therefore grouped together under the same level 1 (L1) category: CD8^+^ T and unconventional T cells.

In total, five major L1-level immune cell groups were defined: CD4⁺ T, CD8^+^ T and unconventional T cells, B cells, Myeloid cells, and NK cells. Each L1 group was subjected to iterative sub-clustering, with cluster resolution progressively optimized to improve granularity. The second annotation level (L2) was defined according to cell developmental states and expression of key functional markers. By further focusing on cluster-specific expression of functional genes, we ultimately defined 58 immune cell types at the third annotation level (L3). These L3-level identities, characterized by unique or highly expressed functional markers, were used in all downstream analyses.

### Single-cell ATAC-seq data preprocessing

Briefly, raw sequencing reads were aligned to the human reference genome GRCh38 using Chromap (v0.2.3-r407)^125^. To retain high-quality cells, barcode beads with fragment counts below the knee point were considered background (empty droplets) and removed. To address potential barcode collisions arising from multiple beads captured in the same droplet, we applied d2c (v1.4.7)^126^ to merge barcode beads that shared the same fragment start and end positions. After barcode merging, MACS2 (v2.2.7.1)^127^ was used to call enriched genomic regions (peaks). Finally, a cell-by-peak count matrix was generated for each library to facilitate downstream analyses.

### Single-cell ATAC-seq data analysis

The scATAC-seq data analysis pipeline was built upon several established computational tools, including scDblFinder (version 1.12.0)^128^, Signac (version 1.9.0)^129^, and SnapATAC2 (version 2.5.1)^130^. This integrated workflow was uniformly applied to all scATAC-seq samples within the cohort. The analysis proceeded through the following steps:

### Per-sample quality control and doublet removal at the single-cell level

To detect doublets, we employed scDblFinder. For each library, a SingleCellExperiment object was first created from the raw count data. We then ran the scDblFinder function with the following parameters: clusters=TRUE, aggregateFeatures = TRUE, nfeatures = 25, and processing = ‘normFeatures’. The doublet prediction results were subsequently incorporated into the cell metadata for downstream analysis. Cells identified as doublets were removed from all libraries. Additional filtering criteria excluded cells failing to meet the following thresholds: fragment count > 1,000, TSS enrichment > 5, FRiP > 0.6, and TSS proportion > 0.2.

#### Batch effect removal, dimensionality reduction and cell clustering

We utilized the SnapATAC2 workflow for batch effect removal, dimensionality reduction, and cell clustering. First, data were imported with the snap.pp.import_data function. We then selected the top 200,000 features using snap.pp.select_features and performed spectral embedding via snap.tl.spectral with default parameters. Batch correction was applied using the Harmony algorithm through the snap.pp.harmony function, specifying batch = ‘individual’, max_iter_harmony = 30, and storing the corrected embeddings in key_added = ‘X_spectral_harmony’. UMAP dimensionality reduction was performed on the top 30 Harmony-corrected dimensions using snap.tl.umap, followed by clustering with the Leiden algorithm. To ensure consistency across multi-omics samples, several sample libraries were removed to facilitate downstream analysis. Clusters exhibiting dominant batch effects were filtered out, resulting in a final scATAC-seq dataset of 1,226,658 high-quality cells. A second round of clustering was then performed using the same SnapATAC2 functions and parameters to further refine and identify distinct cell clusters.

#### Cell type annotation, peak calling and peak-to-gene linkage identification

As detailed in the section Integration of scRNA-seq and scATAC-seq Data, we employed scglue (version 0.2.3)^31^ to facilitate cell type-level cluster annotation across modalities. After GLUE integration, we leveraged the L3 annotation from the scRNA-seq dataset to identify matching cell clusters within the scATAC-seq data. The integrated datasets were embedded into a shared low-dimensional GLUE space for joint analysis.

To identify cell type-specific candidate *cis*-regulatory elements (cCREs), we performed peak calling using the snap.tl.macs3 function with the parameter groupby = ‘L3_annotation’. Following peak detection, we merged the cell type-specific peaks to create a union peak set referencing the GRCh38 genome via thesnap.tl.merge_peaks function. Finally, a peak-by-cell matrix for the scATAC-seq data was constructed using snap.pp.make_peak_matrix to enable downstream analyses of chromatin accessibility at cell type resolution.

### Integration of scRNA-seq and scATAC-seq Data

To integrate unpaired single-cell RNA-seq (scRNA-seq) and single-cell ATAC-seq (scATAC-seq) data for multi-omics analysis, we employed the scGLUE framework. Data Preprocessing: To reduce computational demands, the scRNA-seq dataset was randomly subsampled from 2,744,009 cells to 1,100,000 cells. For the scATAC-seq data, the quality-controlled fragment matrix was used to perform peak calling with the snap.tl.macs3 function, grouping by the “leiden” clustering results (groupby = ‘leiden’). A peak matrix was then constructed, and spectral embedding was generated using the snap.tl.spectral function. Data Integration: Gene location annotations were extracted from the GRCh38 GTF file using scglue.data.get_gene_annotation. A prior regulatory graph was constructed with scglue.genomics.rna_anchored_guidance_graph (rna, atac), mapping highly variable genes (HVGs) onto the scATAC-seq peak matrix.

For the GLUE model configuration, the raw peak count matrix and spectral embedding matrix of the scATAC-seq data served as ATAC inputs, configured via scglue.models.configure_dataset with parameters: prob_model = ‘NB’, use_highly_variable = True, and use_rep = ‘X_spectral’. The raw gene count matrix and PCA-reduced matrix of the scRNA-seq data were used as RNA inputs, configured similarly with prob_model = ‘NB’, use_highly_variable = True, use_layer = ‘counts’, and use_rep = ‘X_pca’. Highly variable subgraphs from both modalities were extracted using guidance.subgraph to generate guidance_hvf. Model training was performed with scglue.models.fit_SCGLUE using the scRNA-seq and scATAC-seq datasets along with guidance_hvf, specifying init_kws = {‘h_dim’: 512, ‘random_seed’: 999} and fit_kws = {‘data_batch_size’: 8192}. After training, the aligned latent representations (X_glue) were extracted for both modalities using glue.encode_data. Cell type annotations from the scRNA-seq data were then transferred to the scATAC-seq data within the integrated latent space by applying scglue.data.transfer_labels.

### DEGs and pathway enrichment analysis

DEGs between disease states and cell types were identified using the rank_genes_groups function from the Scanpy package. The Wilcoxon rank-sum test was applied, comparing groups including IRs_vs_HDs, INRs_vs_HDs, and INRs_vs_IRs. Genes with an adjusted p value (pvals_adj) less than 0.05 and an absolute log fold change greater than 0.58 (|log_2_FC| > 0.58) were considered significant DEGs.

Pathway enrichment analysis was then performed on the identified DEGs. Gene Ontology (GO) enrichment was conducted using the enrichGO function from the clusterProfiler package^131^ with parameters: OrgDb = org.Dr.eg.db, pAdjustMethod = ‘BH’, pvalueCutoff = 0.05, and qvalueCutoff = 0.2. Kyoto Encyclopedia of Genes and Genomes (KEGG) pathway enrichment was performed using the enrichKEGG function with parameters: organism = ‘human’, pAdjustMethod = ‘BH’, pvalueCutoff = 0.05, and qvalueCutoff = 0.2. For a detailed list of DEGs and enrichment results, see Supplementary table 3.

### DAPs and pathway enrichment analysis

Cell type-specific differentially accessible peaks (DAPs) between disease states were identified using the tl.diff_test function from the SnapATAC2 package. The Wald test was applied to compare groups including IRs_vs_HDs, INRs_vs_HDs, and INRs_vs_IRs. Peaks with an adjusted p value < 0.05, minimum log fold change (min_log_fc) ≥ 0.25, and direction set to “both” were considered significant DAPs. Differentially accessible peaks were annotated to corresponding genes using the ChIPseeker package^132^.

Subsequently, pathway enrichment analysis was performed on the annotated genes. GO enrichment was conducted using the enrichGO function from clusterProfiler^131^ with parameters: OrgDb = org.Dr.eg.db, pAdjustMethod = ‘BH’, pvalueCutoff = 0.05, and qvalueCutoff = 0.2. KEGG pathway enrichment was performed using the enrichKEGG function with parameters: organism = “human”, pAdjustMethod = ‘BH’, pvalueCutoff = 0.05, and qvalueCutoff = 0.2. For detailed results, see Supplementary table 4.

### Module scores for feature gene sets expression

To evaluate dynamic changes in biological processes mediated by immune cell subsets, we applied the sc.tl.score_genes function from the Scanpy package.

Parameters were set as ctrl_size = 50 and n_bins = 25. The score for each cell was calculated as the average expression of a given gene set minus the average expression of a matched control gene set. Gene sets used for scoring were derived from the Hallmark Gene Sets (50 gene sets) and the Gene Ontology Biological Process (GOBP) collection (7,608 gene sets) obtained from the MSigDB database^133^. Statistical significance of score differences among HDs, IRs and INRs was assessed using the Wilcoxon rank-sum test. For detailed results, see Supplementary table 5.

### Construction of gene regulatory networks

#### Metacell calculation

We utilized SEACells (version 0.3.3)^134^ to calculate metacells separately for scRNA-seq and scATAC-seq data. Using the SEACells.core.SEACells function, metacells were generated at an approximate ratio of 25:1 cells per metacell for scRNA-seq and 10:1 for scATAC-seq within each cell type per sample. Resulting SEACell IDs were stored in the metadata. Aggregation of raw counts from cells sharing the same SEACell ID was performed with the SEACells.core.summarize_by_SEACell function, preserving sample information and cell type annotations. This process yielded 113,487 scRNA-seq metacells and 126,491 scATAC-seq metacells.

#### Construction of pseudo-multi-omics metacell data

To generate pseudo-multi-omics metacell profiles, we randomly selected metacells from scRNA-seq and scATAC-seq datasets for each cell type per sample, matching pairs based on the smaller metacell count between the two modalities. Paired metacells were assigned a common new metacell barcode, while unpaired metacells were excluded. This resulted in a final set of 78,920 pseudo-multi-omics metacells.

#### Construction of pycisTopic object and topic modeling

A pycisTopic object was created from the scATAC-seq metacell data using the create_cistopic_object function from pycisTopic (version 2.0a0)^41^. Topic modeling was performed using the run_cgs_models_mallet function with parameters: n_topics = [5, 10, 20, 30, 40, 50, 60, 70, 80], n_iter = 200, random_state = 555, alpha = 50 (with alpha_by_topic = True), eta = 0.1 (with eta_by_topic = False), reuse_corpus = True. Model evaluation was conducted using the evaluate_models function, leading to the selection of a 50-topic model, which was then incorporated into the object via add_LDA_model.

#### Candidate enhancer region inference and motif enrichment analysis

Two complementary methods were applied to infer candidate enhancer regions from the pycisTopic analysis: Binarization of Region-Topic Probabilities: The binarize_topics function from pycisTopic was used with both the ‘otsu’ and ‘ntop’ methods. This involved either selecting all regions exceeding a threshold per topic (otsu) or the top 3,000 regions per topic (ntop). Differentially Accessible Regions (DARs): For the dataset encompassing all cell types, regional accessibility was first imputed using the impute_accessibility function, followed by normalization and scaling with normalize_scores (scale_factor = 10^4^). Highly variable regions were identified using the find_highly_variable_features function. DARs were then computed with the find_diff_features function for each L3 cell type across different stage groups. These DARs were considered candidate enhancers for subsequent analysis. Finally, the pycistarget package was utilized to refine candidate enhancer inference by integrating motif-to-transcription factor (TF) annotations. A precomputed motif score database for genomic regions was obtained from https://resources.aertslab.org/cistarget/, enabling motif enrichment analysis and TF prediction for the candidate enhancer regions.

#### Construction of enhancer-based gene regulatory networks

We constructed enhancer-driven gene regulatory networks (eGRNs) following the SCENIC+ workflow (version 1.0a1)^41^. Leveraging cisTopic results and the motif enrichment dictionary, the SCENIC+ package was applied to infer eGRNs by integrating enhancer accessibility and transcription factor binding information.

#### Identify differential transcription factors

The enrichment scores of enhancer-driven gene regulatory networks (eGRNs) were calculated using the AUCell function, which utilizes gene- or region-based rankings derived from expression or accessibility profiles per cell. Key immunity-related genes identified in this study were visualized within TF-eRegulon networks, where intermediate nodes represent target regulatory regions and edge nodes represent target genes. To identify differential eGRNs, we compared transcription factor (TFs) enrichment scores (AUC) across disease states within each cell type using the rank_genes_groups function in Scanpy. The Wilcoxon rank-sum test was employed for statistical analysis, comparing groups IRs_vs_HDs, INRs_vs_HDs, and INRs_vs_IRs. Differential eGRNs were defined by adjusted p values < 0.05 and absolute log fold changes > 0.2. Further details are provided in Supplementary table 6.

### Analysis of ligand-receptor pairs in cell communication

The cell connectome represents a network of intercellular communication mediated by ligand-receptor interactions. Extracellular signaling initiates when ligands bind their cognate receptors, activating specific signaling pathways that regulate tissue homeostasis and biological functions. The connectomeDB 2020 database^135^ provides 2,293 literature-supported ligand-receptor pairs used for this analysis. To infer ligand-receptor interactions between cell types, we leveraged single-cell gene expression data with the CellPhoneDB Python package^52^, which predicts enriched cell-cell interactions. Interaction strength was quantified based on receptor expression in one cell type and corresponding ligand expression in the interacting cell type.

For analysis efficiency and robustness, cells were grouped by disease status into HDs, IRs and INRs, using the L3 cell type annotation for granularity. CellPhoneDB’s statistical analysis was performed with the cpdb_statistical_analysis_method function, applying parameters: counts_data = ‘hgnc_symbol’, score_interactions = True, iterations = 1,000, threshold = 0.1, result_precision = 3, and pvalue = 0.05. Ligands or receptors expressed in at least 10% of cells were considered for significant interactions. Further details are provided in Supplementary table 7.

### Covariate-adjusted residual correlation analysis

To investigate the association between variables of interest (e.g., gene expression levels) and clinical variables (e.g., age), while controlling for potential confounding effects of treatment-related covariates (such as treatment duration and treatment regimen), we applied the following approach^136^: For each pair of independent (**x**, e.g., gene expression) and dependent variables (**y**, e.g., age), we fitted separate linear regression models including treatment-related covariates (treatment duration and treatment regimen) as additional predictors. Residuals were extracted from both models, representing the variation in x and y after removing the effects of the covariates. Spearman rank correlation was then performed between these residuals to assess the true correlation between the variables of interest, effectively controlling for the influence of treatment covariates.

Similarly, when evaluating the correlation between treatment-related variables or other clinical factors and a phenotypic variable, we controlled for the remaining non-target covariates using the same residual-based approach to minimize confounding bias.

### Generation and analysis of genotypes

#### Preprocessing and genotyping

The WGS FASTQ files from 143 samples underwent quality control using fastp (version 0.23.2)^137^. Read alignment, position sorting, duplicate marking, base quality score recalibration (BQSR), and variant calling were performed using ZBOLT (MegaBOLT version 2.3.0)^138^. Joint genotyping and variant quality score recalibration (VQSR) was subsequently conducted using GATK (version 4)^139^. Further quality control was performed using PLINK (version 1.9)^140^ to prepare the dataset for imputation. Specifically, biallelic variants were extracted, variants with missing genotype rates exceeding 2% were removed, and a sex check was conducted. Additional filtering criteria included a minor allele frequency (MAF) threshold of 0.01 and a Hardy-Weinberg equilibrium (HWE) p value cutoff of 1×10⁻^6^.

#### Imputation of genotyping data and principal component analysis

Genotype imputation was performed using Beagle (version 4.1)^141^ with the high-coverage 1,000 Genomes Project (1KGP) reference panel^142^ (n = 3,202 samples). Variants with an imputation quality score (DR2) greater than 0.3 were retained. Post-imputation quality control was conducted using PLINK, including filtering for biallelic variants, missing genotype rate < 2%, minor allele frequency (MAF) ≥ 0.01, and Hardy-Weinberg equilibrium (HWE) p value > 1×10⁻^6^. Heterozygosity rates were computed, and individuals with values outside ±4 standard deviations from the mean were excluded. After removing individuals lacking both scRNA-seq and scATAC-seq data, the final cohort consisted of 142 individuals. A total of 7,540,937 variants passed all filters and were retained for downstream analyses.

### Lineal model-based *cis*-xQTL detection by TensorQTL

Cell type-specific pseudobulk gene expression and peak accessibility matrices were utilized for *cis*-xQTL mapping. We applied TensorQTL (version 1.0.10)^143^ to perform linear model-based *cis*-eQTL and *cis*-caQTL analyses. After excluding cell types with fewer than 80 pseudobulk samples (each sample comprising ≥ 10 cells), 51 L3 cell types were included for *cis*-eQTL and 31 L3 cell types for *cis*-caQTL analyses. Genes and peaks expressed or accessible in fewer than 90% of samples per cell type were filtered out. The remaining pseudobulk matrices were normalized across samples by ordered quantile normalization using the QuantileTransformer and StandardScaler functions from the Python package sklearn (version 1.8.3)^144^, and used as phenotype inputs in TensorQTL.

Covariates included age, treatment duration, the first two genotype principal components (PCs), and PEER factors. PEER factors were computed using the PEER R package (version 1.0)^145^, based on the top 2,000 highly variable genes or peaks per cell type; if fewer than 2,000 genes were available, all were used. For *cis*-eQTL mapping, variants within ±1 Mb of the gene transcription start site (TSS) were tested; for *cis*-caQTL mapping, variants within ±1 Mb of the peak midpoint were included. The map_nominal function was used to obtain nominal p values for variant-gene and variant-peak pairs. Subsequently, map_cis performed Beta-approximation permutation tests to generate molecular phenotype-level summary statistics with empirical p values.

Results from all cell types were aggregated, and study-wide false discovery rates (FDR, q-values) were calculated using the calculate_qvalues function. Genes and peaks with q-value < 0.05 were annotated as *cis*-eGenes and *cis*-caPeaks, respectively. The variant with the lowest nominal p value for each *cis*-eGene or *cis*-caPeak was designated as the lead *cis*-xQTL. For additional details, see Supplementary table 8.

### INRs related *cis*-xQTL identified by interaction QTL mapping

To identify genotype-by-disease-group interactions specific to INRs, we performed interaction *cis*-xQTL mapping using TensorQTL^143^. For each cell type, pseudobulk gene expression and peak accessibility profiles were generated and formatted as BED files associating each phenotype with its genomic coordinates. To control for confounding variation, covariates including age, treatment duration, the first two genotype principal components (PCs), and PEER factors were precomputed and incorporated.

Samples were stratified into three disease status groups: HDs, IRs and INRs. For interaction analysis, HDs and IRs were combined into a single reference group, with INRs treated as a distinct group. This created a binary interaction variable used to build the interaction design matrix. We performed interaction-aware nominal *cis*-xQTL mapping considering variants within ±1 Mb of each gene transcription start site (TSS). The minor allele frequency threshold for interaction terms was set at 0.1. Multiple testing correction was conducted via empirical adjustment using eigenMT, followed by Benjamini-Hochberg (BH) FDR correction.

The fitted linear model for each gene-SNP pair included the main genotype effect (G), main disease group effect (I), genotype-by-group interaction term (G×I), and PEER covariates. Interaction terms with BH-adjusted p values < 0.05 were deemed significant, indicating SNP effects on gene expression that differ specifically in INRs compared to the combined HDs and IRs. For additional details, see Supplementary table 9.

### Poisson distribution model-based *cis*-eQTL detection by SAIGEQTL

To systematically evaluate the impact of genotype variation on single-cell gene expression, this study used the SAIGEQTL (Single-cell Aware eQTL) analysis framework. The analysis process consists of three main steps: (1) fitting a null Poisson mixed model; (2) univariate association test; and (3) gene-level p-value integration (ACAT test)^146^.

Step 1: Fitting a Null Poisson Mixed Model. In each cell type, a generalized linear mixed model (GLMM) was fitted for each gene, assuming that the expression level follows a Poisson distribution. The model simultaneously considers inter-sample correlation (random effect) and cell-level confounding factors (fixed effect), including covariates such as age, treatment time, total counts (log_total_counts), mitochondrial proportion (pct_counts_mt), genomic components (PC1-PC2), and expression principal components (PC1-PC5).

Step 2: Single-Variant Association Test. Based on the null model obtained in Step 1, SAIGEQTL performed univariate association tests for SNPs within the ±1 Mb interval of each gene to assess the relationship between genotype and gene expression. In this analysis, loci with a minor allele frequency (MAF) below 0.1 or a minor allele number (MAC) less than 20 were filtered out. Association tests were performed using the score test method under a generalized mixed model, corrected by the saddlepoint approximation (SPA) to improve the accuracy of significance estimates for rare variants.

Step 3: Gene-level significance test (ACAT). To obtain overall gene significance, the ACAT (Aggregated Cauchy Association Test) method was used to aggregate the p-values of all SNPs within a single gene. ACAT uses a Cauchy-distributed weighted summation approach to achieve fast and robust integration of multiple tests, maintaining statistical power across different association signal patterns. For additional details, see Supplementary table 10.

### Comparison of *cis*-eQTLs in TensorQTL and SAIGEQTL

A Venn diagram visualized the overlap of *cis*-eGenes in SAIGEQTL (sc-eQTL) and TensorQTL (bulk-eQTL). For this visualization, we unified the *cis*-eQTL data from SAIGEQTL and TensorQTL by grouping the cell types into four broad categories: CD4 Tn cells, CD4 Tm cells, CD4 Th cells, and CD4 Treg cells. For comparison, we designated the cell types in the TensorQTL cohort as reference cell types, while the corresponding cell types in the SAIGEQTL cohort served as query cell types. We calculated statistics and conducted independent lead *cis*-eQTL analyses using the same methods used to detect cell-type-specific genetic effects. Furthermore, we obtained the genetic effect sizes of the lead *cis*-eQTLs in the reference cell types and extracted the corresponding effect sizes of the same *cis*-eQTLs in the query cell types to assess the correlation of genetic effects.

### Defining INRs-related inflammation modules

To calculate cell-state interaction eQTLs (ieQTLs), we customized an inflammation module related to INRs. This module includes five signaling pathways: inflammatory response, interferon-γ response, TNFα signaling via NF-κB, activation, and apoptosis. We used the sc.tl.score_genes function from the Scanpy package, with parameters set to ctrl_size = 50 and n_bins = 25. The score for each cell was calculated as the mean expression of the given gene set minus the mean expression of the matched control gene set. The Wilcoxon rank sum test was used to assess the statistical significance of differences in scores between HDs, IRs, and INRs.

### ieQTL calling

To examine whether the relationship between gene expression and genotype changes depended on cell state, we added an interaction term between the genotype and the cell state. Cell states were defined as biologically interpretable module expression (continuous) or disease status of the donor (binary). The likelihood ratio test (analysis of variance (ANOVA) function in R) comparing the model with and without an interaction term was used to calculate P values. Specifically, the χ^2^ statistic was calculated as - ×2 log (likelihood ratio), which was compared against χ^2^ distribution with one degree of freedom. Such a test was performed to a broad set of significant eQTLs called from all cells including HDs, IRs and INRs cells. For additional details, see Supplementary table 11.

### Integrative analysis of ieQTLs

Significant ieQTLs were used for integrated analysis. AUC scores calculated by SCENIC+ were used as an estimate of TF activity. To calculate the correlation between TF activity and cell state, the Pearson correlation between the expression of gene corresponding to TF and the expression of inflammatory module was calculated. For TF-binding motif analysis, nucleotide sequences flanking 15 bp on each side of eSNP were submitted to the MEME suite FIMO software^147^. Motif occurrences were determined by the FIMO P < 0.01. If the TF motif occurs in the sequence with the reference allele, but does not occur in an alternative allele, or vice versa, it was designated as an eSNP that creates or disrupts the TF motif. Quartets composed of TF, cell state, ieSNP and ieGene that satisfy the following criteria were selected: (1) significant ieQTL, (2) Pearson’s correlation coefficient between expression of gene corresponding to TF and module expression >0.2 and (3) ieSNP that creates or disrupts the TF motif.

### scRNA-seq and scATAC-seq data processing for scPRISM model development

For the transcriptomic analysis, normalization of the single-cell RNA sequencing dataset was performed by adjusting for total cellular read counts with a scaling factor of 10,000, followed by a logarithmic transformation (log1p), utilizing Scanpy (v1.10.4). A feature selection step was then conducted, wherein we isolated the 2,490 most highly variable genes (HVGs) while removing genes not detected in more than 90% of the cell population. The gene set was finalized by incorporating a curated list of genes and subsequently removing any without overlap in accessible chromatin regions, yielding 582 genes for subsequent analysis.

For the chromatin accessibility analysis, the raw peak count matrix was subjected to a term frequency-inverse document frequency (TF-IDF) transformation (MUON v0.1.6) to account for technical variability. The resulting matrix then underwent normalization with a scale factor of 10,000 and a log1p transformation. Feature selection involved retaining the 50,000 most prevalent peaks, from which we excluded peaks detected in fewer than 10% of cells, resulting in a final collection of 5,583 chromatin accessible peaks for our computational framework.

### Cell pairing and multi-omics data integration

#### Sample selection and feature dimensionality reduction

Our dataset was derived from 142 patient samples with paired scRNA-seq (2,700,232 cells) and scATAC-seq (1,216,704 cells) profiles, covering 57 L3-level cell types. To create a shared feature space for cell-to-cell alignment, we focused on 582 target genes, using their transcript levels from scRNA-seq and gene activity scores from scATAC-seq. For each cell type, we performed principal component analysis (PCA) on both the expression and gene activity matrices, selecting the top 50 PCs from each to serve as the latent representation for the pairing task.

#### Cell pairing via bipartite graph

We employed a cross-modal mapping strategy to align cells from the scRNA-seq and scATAC-seq datasets. The approach is established on a mutual k-nearest neighbor (mKNN) algorithm operating in the shared principal component space. For each cell, we identified its k nearest neighbors in the opposing modality based on Euclidean distance, establishing reciprocal relationships. These relationships defined a bipartite graph, which was encoded as a cost matrix where the cost of pairing two cells was their distance. A significant advantage was conferred to robust pairings by applying a 50% cost reduction to all mutual nearest neighbor pairs. We then determined the optimal global assignment that minimized the total pairing cost by solving the linear assignment problem with the Hungarian algorithm, as implemented in SciPy (v1.13.0). We empirically set the neighborhood parameter k equal to 35.

### Iterative matching strategy

To maximize cell utilization and enhance the cross-modal integration efficiency beyond single-pass limitations, we implemented a multi-round refinement strategy to progressively match the remaining pools of unassigned cells from each modality. Following the initial pairing, the cohort of unpaired scRNA-seq cells and the corresponding cohort of unpaired scATAC-seq cells were subjected to another full iteration of the matching procedure. This iterative process was repeated until a convergence criterion was met, specifically when over 90% of the cells in the smaller of the two remaining populations were successfully paired in an iteration. Upon completion, the collection of all matched pairs from every round was consolidated to form the final dataset. This hierarchical strategy significantly increased the yield of the integration, resulting in a final dataset of 2,557,104 high-confidence cell pairings.

### Dataset splitting

We employed two distinct dataset partitioning strategies tailored to our predictive modeling objectives. For the disease state classification task, we established an independent test set by sequestering all data from a random subset of 15 donors (five from each HDs, IRs, and INRs), thereby preventing patient-specific information leakage between training and testing. Model development was then conducted on the cells from the remaining individuals using a ten-fold cross-validation scheme. Conversely, for predicting gene expression from chromatin accessibility, a paired cell-based random split was utilized. The complete dataset was partitioned into a training set (60%), a validation set (10%), and a test set (30%) to enable model training, hyperparameter tuning, and final performance assessment, respectively.

### Model architecture and development

Our scPRISM employed a dual-pathway architecture tailored for cross-modal analysis of single-cell multi-omics data. Paired cells data from both scRNA-seq and scATAC-seq modalities were first transformed through dedicated embedding layers, then processed by separate encoder networks to capture modality-specific latent representations. These encoded features were subsequently fed into two specialized prediction branches: the scPRISM-CLIN branch integrates both scRNA and scATAC representations through a fusion decoder to enable disease state classification, while the scPRISM-EXPR branch leverages the scATAC-derived features alone to reconstruct gene expression patterns.

#### Peak and gene masking

To encode whether a chromatin accessible peak was located within a ±250 kb windows of a target gene, we first defined a binary indicator matrix 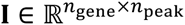, where

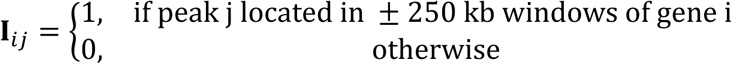

The peak masking matrix for paired scATAC-seq cells was then defined as

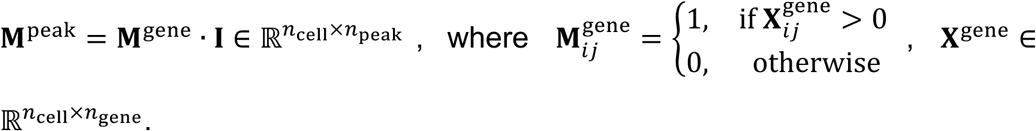

#### Numeric value embedding

Our scPRISM began with the float value embedding module that projected an input numeric feature of a paired cell 𝐱_𝑖_ ∈ ℝ^𝑑^ to an embedding matrix 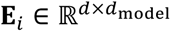. The embedding vector 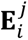 for paired cell 𝑖 with 𝑗 -th feature 𝐱_𝑖j_ was then calculated as element-wise multiplication with bias addition:

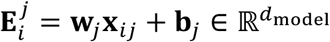

where 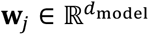 is a trainable weight vector, and 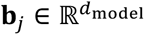 is the 𝑗-th feature bias.

Thus, the embedding matrix of paired cell 𝑖 was represented as 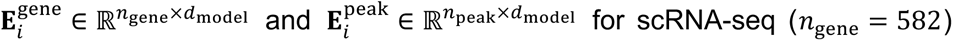 and scATAC-seq (𝑛_peak_ = 5583) data, respectively . We set 𝑑_model_ = 64 and initialized 𝐰_j_ using the Kaiming uniform distribution.

#### Encoder module

Both the scRNA-seq and scATAC-seq encoders in scPRISM were architecturally identical, each consisting of seven stacked layers. Every layer featured an eight-head self-attention mechanism and a subsequent feed-forward network. These two core components were consistently augmented with residual connections and layer normalization to facilitate deep learning.

For a paired cell 𝑖, the embedding matrix 𝐄_𝑖_ was linearly projected to query 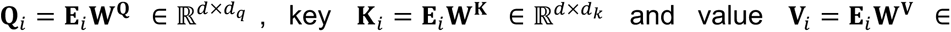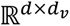. For simplicity, we set 𝑑_𝑞_ = 𝑑_𝑘_ = 𝑑_𝑣_ = 𝑑_model_. For each of the eight head ℎ, self-attention was computed as:

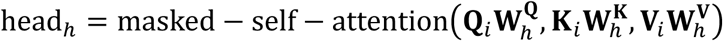

where 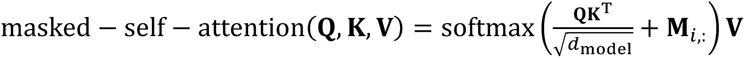, 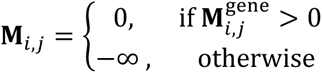 for paired scRNA-seq cell or 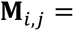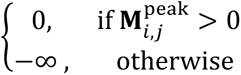 for paired scATAC-seq cell, 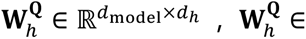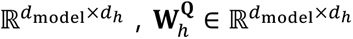. We set 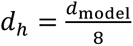. Self-attention computations were accelerated during training with the memory-efficient Flash-attention algorithm^148^.

The concatenated output was then given by

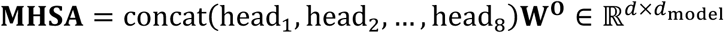

_where_ 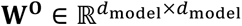.

The output from the 𝑙-th layer of multi-head self-attention module was:

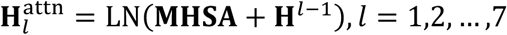

where 𝐇^0^ = 𝐄, LN is layer normalization. 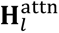 was then passed to the feed-forward network module, followed by dropout with ratio of 0.1,

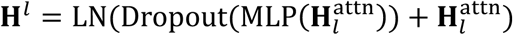

Consequently, the final encoded hidden state of a paired cell 𝑖 for scRNA-seq and scATAC-seq were 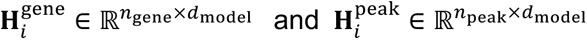.

### scPRISM-CLIN and scPRISM-EXPR modules

For a paired cell 𝑖, the cell embedding represented by hidden states of genes and peaks were repectievly obtained by

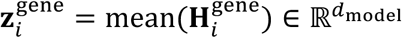

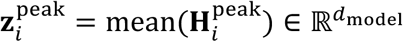

For scPRISM-CLIN module, we implemented fusion encoder for integrating both scRNA-seq and scATAC-seq by simply concating cell embedding of genes and peaks:

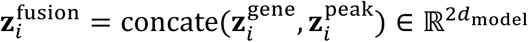

Then, 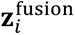 was input into a gated MLP prediction head to predict disease state.

For scPRISM-EXPR module, we took 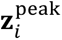 as input to an independent gated MLP prediction head to prediction gene expression.

The gated MLP was defined as:

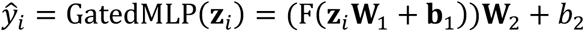

where 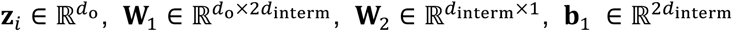 , 𝑏_2_ is a scalar bias for the final prediction, F(𝐔) = 𝐔[: , : 𝑑_model_]⊗GELU(𝐔[: , 𝑑_model_: ]), GELU is the Gaussian error linear unit. We set 𝑑_interm_ = 𝑑_o_, and 𝑑_o_ = 2𝑑_model_for scPRISM-CLIN module while 𝑑_o_ = 𝑑_model_ for scPRISM-EXPR module.

### Model training and evaluation

Our model was trained for two distinct tasks using different objective functions. For the disease state classification task, we trained three separate scPRISM-CLIN models to perform pairwise comparisons (INRs vs IRs, INRs vs HDs, and IRs vs HDs). These models were trained using a binary cross-entropy with logits loss function (BCEWithLogitsLoss). For the gene expression prediction task with the scPRISM-EXPR module, we used the mean squared error (MSE) as the loss function. All models were optimized using the AdamW optimizer with a weight decay of 0.01 and default 𝛽 parameters (𝛽_1_ = 0.9, 𝛽_2_ = 0.999). We initiated training with a learning rate of 0.0001 and implemented a step decay schedule, reducing the rate by a factor of 0.9 every 50 epochs. Models were trained with a batch size of 128 for a maximum of 100 epochs. To prevent overfitting, we employed an early stopping criterion with a patience of 15 epochs, halting training if the validation loss did not improve.

For evaluating disease state classification at patient-level prediction on the test set, all cells from a single individual were processed by the trained model to yield cell-specific labels. The final classification for the individual was then determined via a majority voting mechanism across all of their constituent cells.

### Model interpretation

#### Feature attribution

We dissected the decision-making process of our models by quantifying the importance of each input feature using Integrated Gradients (IG) (Captum v0.7.0). IG calculates feature contributions for a given prediction by accumulating the gradients relative to the input features along a path from a zero-vector baseline to the specific input of interest, which we approximated over 50 uniform steps. We used the absolute attribution scores to measure the strength of each feature’s influence. By applying this procedure to every cell in our independent test set, we generated a comprehensive map of cell-level feature importance. These granular attributions were then aggregated to derive both global feature rankings (by averaging across all test cells) and cohort-specific feature signatures (by averaging across cells within a given disease or control group) for our gene expression and disease state prediction tasks.

#### Distilling a core set of disease-associated genes

To distill a robust signature of genes central to the disease phenotype, we established a two-pronged selection strategy based on both model influence (via scPRISM-CLIN) and predictive accuracy (via scPRISM-EXPR). First, we leveraged the IG method to identify the top 60 genes with the highest mean absolute attribution scores for each of the three disease state comparisons (INRs vs IRs, INRs vs HDs, and IRs vs HDs). Concurrently, we identified the 60 genes whose expression levels were most accurately predicted by the scPRISM-EXPR module, selecting for genes with a Spearman correlation coefficient greater than 0.8 between predicted and observed values. The intersection of the gene sets derived from these four independent analyses, three from model attribution and one from predictive correlation, resulted in a core set of 32 genes poised to be significant drivers of the disease state.

#### Identifying significant peak-gene interactions

We identified significant peak-gene interactions using a two-stage approach. First, we used the scPRISM-EXPR model to perform an initial screening, retaining candidate peaks within a ±250 kb window of a gene if their IG score was greater than 0.005. Second, we validated these candidates with a permutation test. For each candidate peak, we generated 100 background peaks matched for GC content and signal intensity using pychromvar software (v0.0.4). We then computed the Spearman correlation between the gene’s expression and the candidate peak’s accessibility ( 𝜌_obs_ ). A null distribution of correlations was generated by correlating the gene’s expression with the 100 background peaks. The significance of the observed correlation was assessed by calculating a 𝑍-score relative to this null distribution’s mean (𝜇_bg_) and standard deviation (𝜎_bg_), from which a two-sided p value was derived.

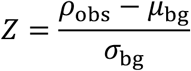

The resulting set of statistically significant interactions (P < 0.05) served as a benchmark to evaluate the regulatory links predicted by our model.

### Computational environment

Our model was implemented in Python (v3.10.8), leveraging the PyTorch (v2.4.1) and PyTorch-Lightning (v2.2.5) frameworks for development and training. All computational tasks were performed on a high-performance workstation featuring four NVIDIA A100 GPUs with 40 GB of VRAM each, 128 Intel Xeon Gold CPUs, and 250 GB of system RAM.

## Data availability

All data of this research have been deposited to CNGB Nucleotide Sequence Archive, OMIX and GVM databases, including: processed genomic VCF files (CNSA, GVM), scRNA-seq matrix TXT files (CNSA, OMIX), scATAC-seq matrix MTX files (CNSA, OMIX) and *cis*-xQTL summary statistics files (Supplementary table 8 and 9). All data were analyzed with standard programs and packages, as detailed below.

## Code availability

Details of publicly available software used in the study are given in the Methods. All complete analytical and visualization codes were uploaded on GitHub. Any additional information required to reanalyze the data reported in this paper is available from the lead contact upon request.

## Acknowledgements

We sincerely thank all participants for their invaluable contributions and our team members for their support. This work was supported by the National Key Research and Development Program of China (2025YFC3409300 to Chuanyu Liu), National Natural Science Foundation of China (32300769 to Y.W.), China Postdoctoral Science Foundation (2023M732369 to Y.Y.), Guangdong Basic and Applied Basic Research Foundation (2024B1515230003 to Chuanyu Liu), Hangzhou Biomedical and Health Industry Development Support Science and Technology Special Project (2022WJC174 to J.Yu), Zhejiang Science and Technology Department (2025C01092), Hangzhou Science and Technology Department (2024SZD1B09), Construction Fund of Key Medical Disciplines of Hangzhou (2025HZZD13 and 2025HZGF09) and Research Grants Council (C4024-22GF, R4007-23 to Xin Wang) of the Hong Kong Special Administrative Region, China. We would like to thank DCS Cloud (https://cloud.stomics.tech/) for providing the computational resources and software support necessary for this study.

## Author contributions

Chuanyu Liu, J.Yu, Y. W. and S.Y. conceived the idea; Chuanyu Liu, J.Yu, Jianhua Yin, J.H. and Xiangdong Wang supervised the work; Chuanyu Liu, J.Yu, Y.W. and S.Y. designed the experiment; Y.Y., J.C., M.Z., S.D., X.S., Z.Y., D.Y., J.Yan, Z.W., W.H., L.S. and B.W. performed the majority of the experiments; S.Y., J.C., L.Z. and Y.W. performed data preprocessing and quality evaluation; S.Y. and J.C. analyzed the data; Y.B., T.L. and X.Z. developed the prediction model; S.Y., Y.Z., Z.H., Xue Wang and X.Lin. provided technical support; Y.Y., Chang Liu, Y.H., J.S. L.G., Jiefang Yin, Y.F., C.X., Xin Wang, Y.Z., J.X., Y.C., W.L., B.S., R.C., Z.C., H.L., L.C., X.Y., X.Liu, Y.G. and L.L. gave the relevant advice; Y.W., S.Y., Y.B., Chuanyu Liu, Jianhua Yin and J.Yu wrote the manuscript.

## Competing interests

The employees of BGI have stock holdings in BGI.

---

**Supplementary Figure 1.**
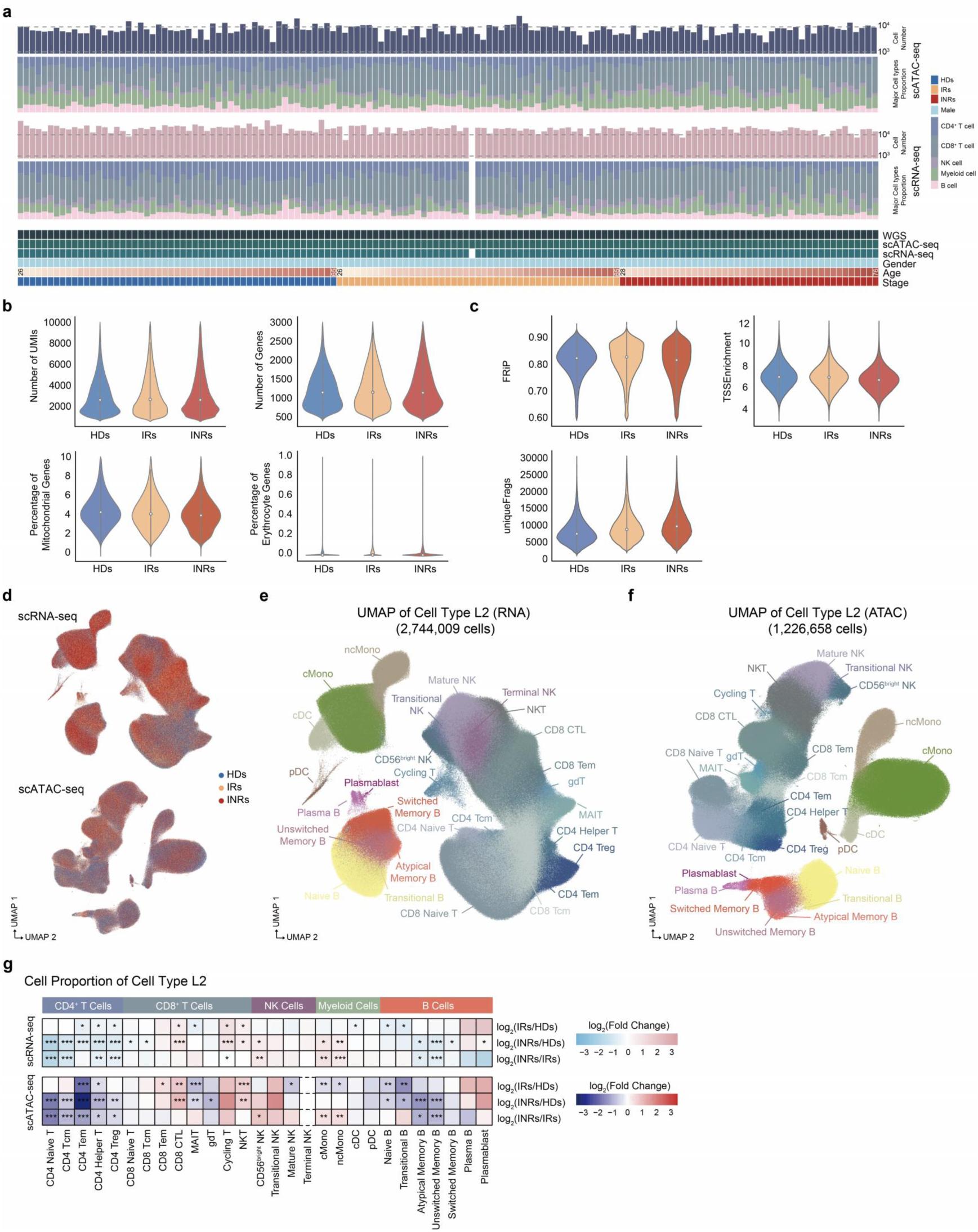
Summary of sample information, quality control results, clustering and annotation. **a**, The 143 individuals are arranged in ascending age order by different groups, showing whether different types of data were collected for each individual, the number of cells detected by scRNA-seq and scATAC-seq and the relative proportion of the L1 compartments identified in the study. **b**, The violin plots showing the number of UMIs, number of genes, proportion of mitochondrial genes and proportion of erythrocyte genes after quality control of HDs, IRs and INRs in scRNA-seq data. **c**, The violin plots showing the number of fractions of all mapped reads that fall into peak regions (FRiP), TSSEnrichment and uniqueFrags after quality control of HDs, IRs and INRs in scATAC-seq data. **d**, UMAP plots showing the batch from scRNA-seq and scATAC-seq data, with each group represented by a different color. **e**, UMAP plot showing 2,744,009 immune cells from scRNA-seq data that have been reduced, clustered, and annotated. At L2, cells are classified into 28 immune cell subtypes, with each cell type represented by a different color. **f**, UMAP plot showing 1,226,658 immune cells from scATAC-seq data that have been reduced, clustered, and annotated. At L2, cells are classified into 27 immune cell subtypes, with each cell type represented by a different color. **g**, Heatmap showing the differences in cell proportions of L2 (28 cell subtypes) between HDs, IRs and INRs in scRNA-seq data and scATAC-seq data.

**Supplementary Figure 2.**
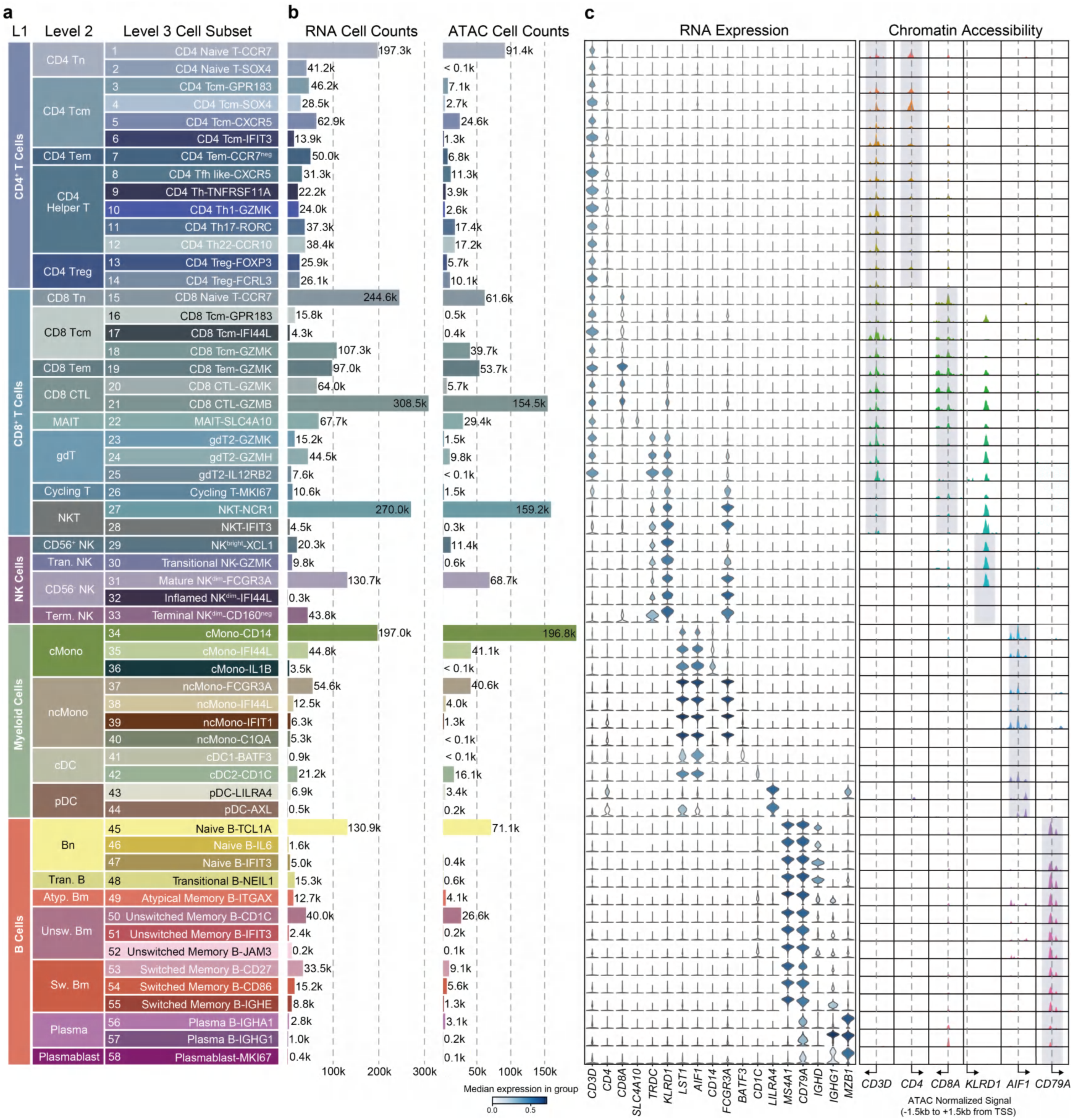
Classical marker gene and cell counts of 58 immune cell subsets in the atlas hierarchy. **a**, The bar chart showing the hierarchical structure of immune cell annotation based on the expression of specific marker genes in scRNA-seq data. L1 represents the five major immune cell types. L2 (28 subtypes) represents immune cell types at different granularities. L3 contains 58 immune cell types marked by specific genes, represented by numbers and colors. **b**, The bar graphs showing the cell counts for each L3 cell type captured by scRNA-seq and scATAC-seq. **c**, Violin plots and track plots showing RNA expression and chromatin accessibility of classical marker genes used to annotate L1 immune cells.

**Supplementary Figure 3.**
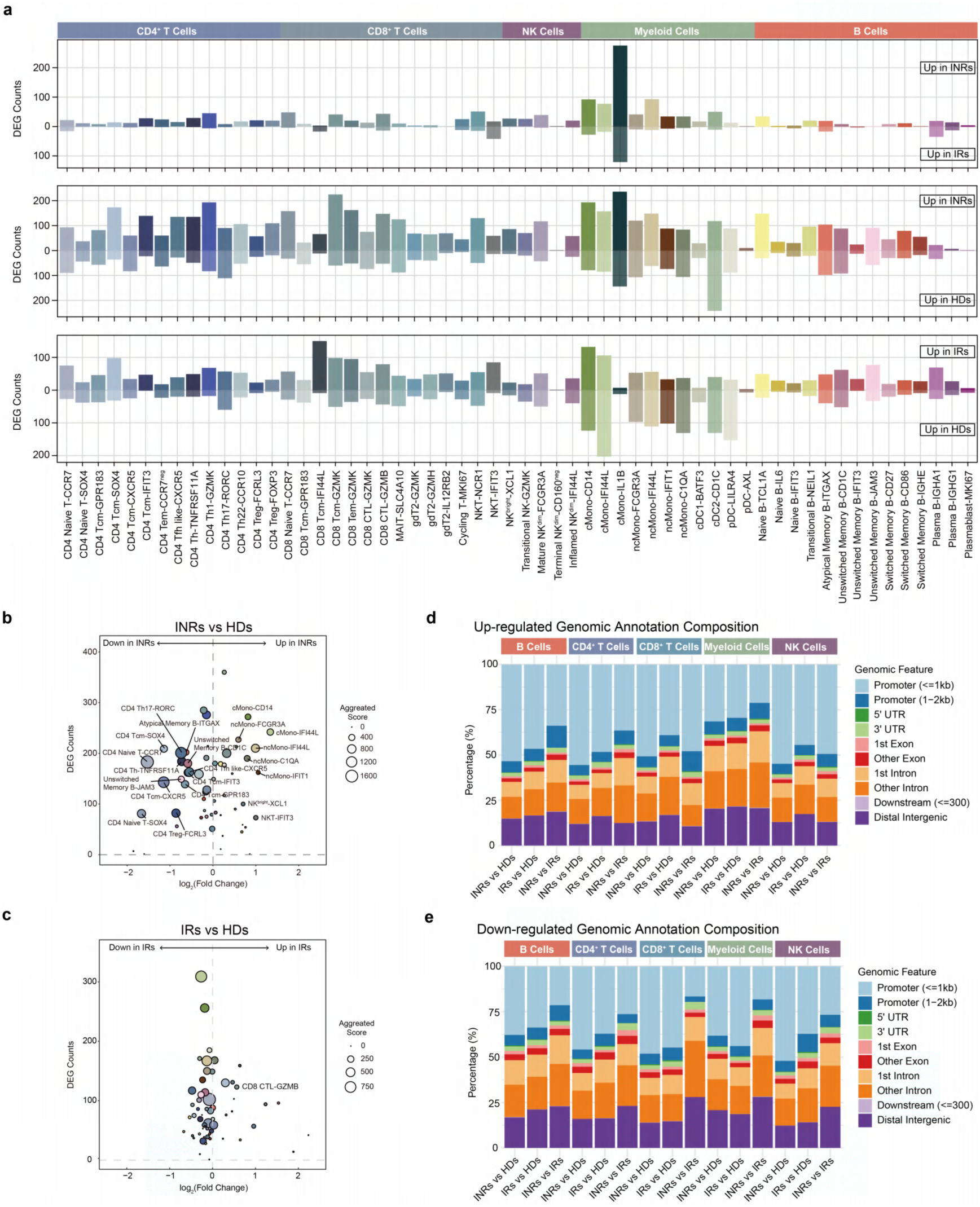
Differential gene statistics, key cell type identification and differential peak region annotation. **a**, Bidirectional bar graphs showing the number of DEGs in immune cell subsets, comparing HDs, IRs and INRs. All compared DEGs were defined as |log_2_foldchange| > 0.58 and P adjust < 0.05. **b**, Bubble plots comparing changes in cell proportions and differentially expressed gene (DEG) counts between INRs and HDs. Bubble size represents a composite measure of change, defined as -log_10_(P adjust of cell proportion change) × DEG Counts. P adjust of cell proportion change was determined using the Wilcoxon rank sum test with Benjamini-Hochberg correction. **c**, Bubble plots comparing changes in cell proportions and DEG counts between IRs and HDs. Bubble size represents a composite measure of change, defined as -log_10_(P adjust of cell proportion change) × DEG Counts. P adjust of cell proportion change was determined using the Wilcoxon rank sum test with Benjamini-Hochberg correction. **d**, Stacked bar chart showing up-regulated genomic annotation composition in different cell types. **e**, Stacked bar chart showing down-regulated genomic annotation composition in different cell types.

**Supplementary Figure 4.**
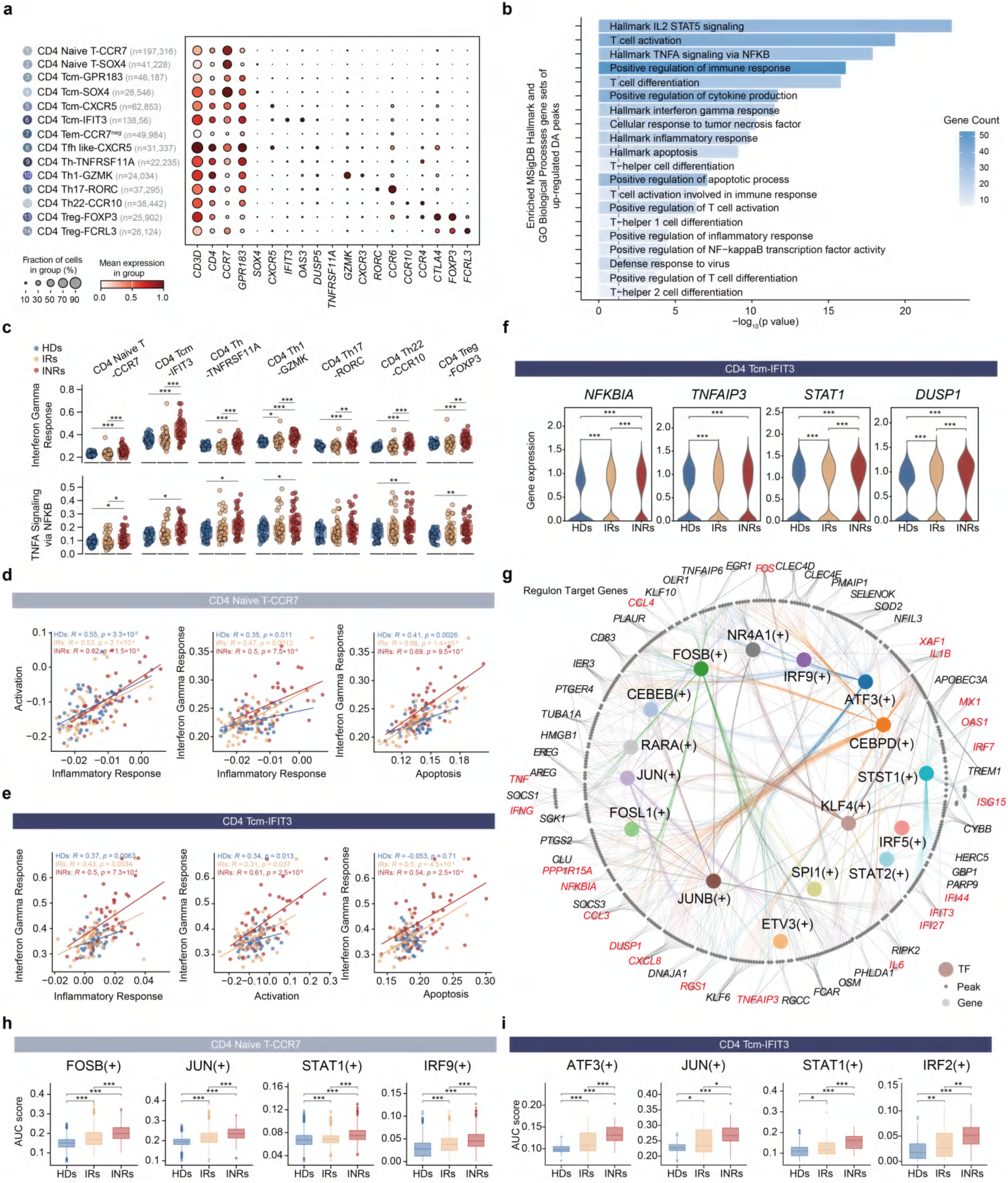
CD4^+^ T cells in INRs undergo apoptosis due to overactivation. **a**, Bubble plot showing the expression of marker genes used to annotate L3 CD4^+^ T cells. **b**, The bar graph showing the top 20 pathways enriched by genes annotated with up-regulated DA peaks. **c**, Boxplots showing the scores of Interferon Gamma Response and TNFA Signaling via NFKB for seven CD4^+^ T cell types in HDs, IRs, and INRs. Different groups are shown in different colors, the horizontal line represents the median, and the whiskers extend to the furthest data point within a maximum of 1.5 times the interquartile range. **d**, The scatter plot showing the correlation between Inflammatory Response, Activation, Inflammatory Response and Interferon Gamma Response in CD4 Naive T-CCR7 in HDs, IRs, and INRs. Different groups are shown in different colors. **e**, The scatter plot showing the correlation between Inflammatory Response, Activation, Inflammatory Response and Interferon Gamma Response in CD4 Tcm-IFIT3 in HDs, IRs, and INRs. Different groups are shown in different colors. **f**, Violin plots showing gene expression of *NFKBIA*, *TNFAIP3*, *STAT1*, and *DUSP1* in CD4 Tcm-IFIT3 in HDs, IRs, and INRs. Different groups are shown in different colors. **g**, The Cytoscape diagram showing the immune function regulatory network composed of TFs, peaks and target genes. For ease of visualization, only differentially expressed target genes in immune cells were selected for each TF, and important target genes were indicated in red. **h**, Box plots showing the AUC values of FOSB, JUN, STAT1, and IRF9 in CD4 Naive T-CCR7 in HD, IRs, and INRs. Different groups are shown in different colors. **i**, Box plots showing the AUC values of ATF3, JUN, STAT1, and IRF2 in CD4 Tcm-IFIT3 in HD, IRs, and INRs. Different groups are shown in different colors.

**Supplementary Figure 5.**
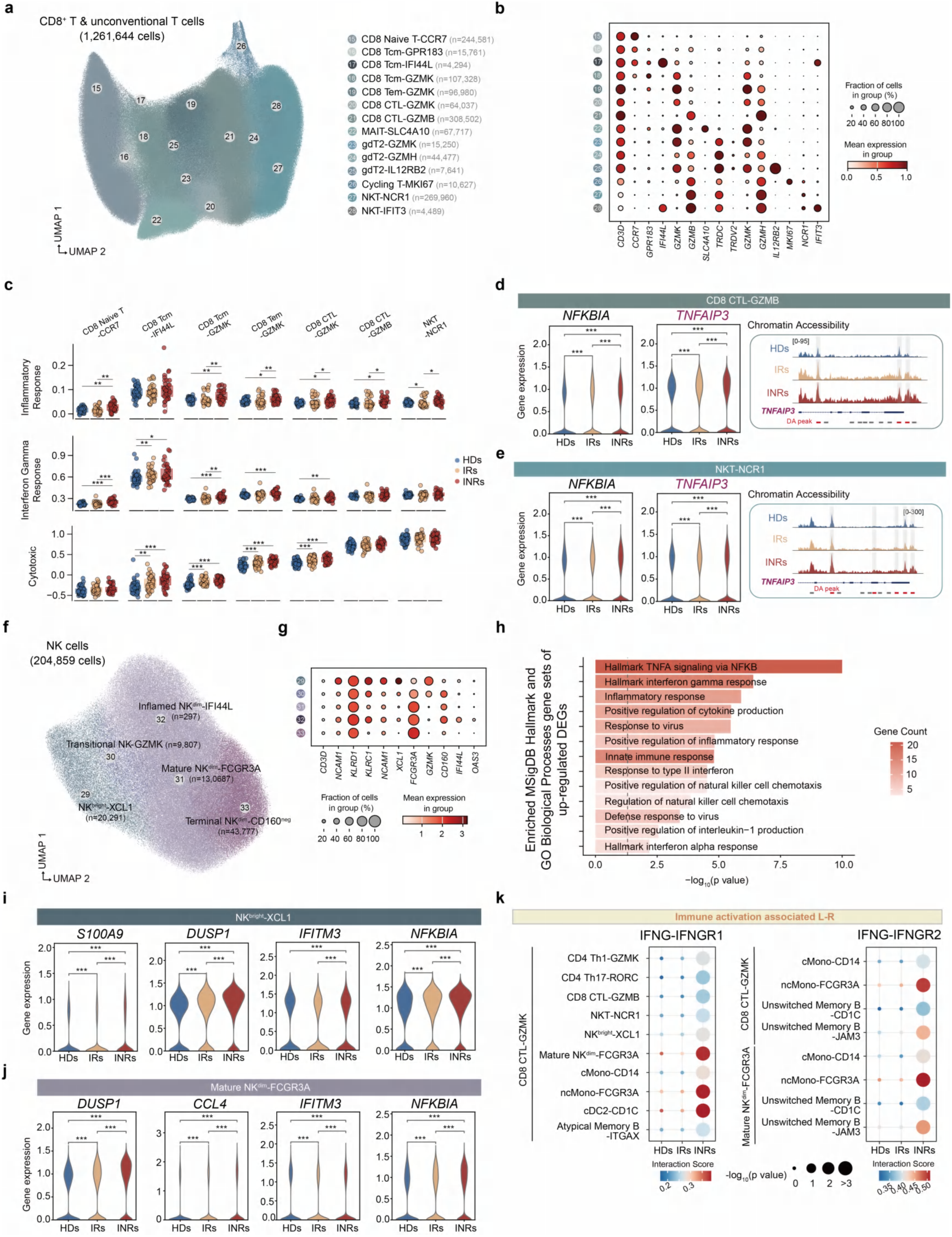
Sustained activation of CD8^+^ T cells and NK cells in INRs. **a**, The UMAP plot showing 1,261,644 CD8^+^ T cells from scRNA-seq data, which are divided into 14 cell subtypes, with each cell type represented by a different color. **b**, Bubble plot showing the expression of marker genes used to annotate L3 CD8^+^ T cells. **c**, Boxplots showing the scores of Inflammatory Response, Interferon Gamma Response and Cytotoxic for seven CD8^+^ T cell types in HDs, IRs, and INRs. Different groups are shown in different colors, the horizontal line represents the median, and the whiskers extend to the furthest data point within a maximum of 1.5 times the interquartile range. **d**, Violin plots showing gene expression of *NFKBIA* and *TNFAIP3* and scATAC-seq tracks revealing gene structure and significant DA peaks for *TNFAIP3* in CD8 CTL-GZMB in HDs, IRs, and INRs. Different groups are shown in different colors. **e**, Violin plots showing gene expression of *NFKBIA* and *TNFAIP3* and scATAC-seq tracks revealing gene structure and significant DA peaks for *TNFAIP3* in NKT-NCR1 in HDs, IRs, and INRs. Different groups are shown in different colors. **f**, The UMAP plot showing 204,859 NK cells from scRNA-seq data, which are divided into 5 cell subtypes, with each cell type represented by a different color. **g**, Bubble plot showing the expression of marker genes used to annotate L3 NK cells. **h**,The bar graph showing the top 13 pathways enriched by up-regulated DEGs. **i**, Violin plots showing gene expression of *S100A9*, *DUSP1*, *IFITM3*, and *NFKBIA* in NK^bright^-XCL1 in HDs, IRs and INRs. Different groups are shown in different colors. **j**, Violin plots showing gene expression of *DUSP1*, *CCL4*, *IFITM3*, and *NFKBIA* in Mature NK^dim^-FCGR3A in HDs, IRs, and INRs. Different groups are shown in different colors. **k**, The bubble plots showing the changes in potential immune activation-related ligand-receptor pairs between immune cell subsets in HDs, IRs, and INRs, including IFNG-IFNGR1 and IFNG-IFNGR2. The color of the dots indicates the interaction score calculated using CellPhoneDB, and the size of the dots indicates the - log_10_(p value).

**Supplementary Figure 6.**
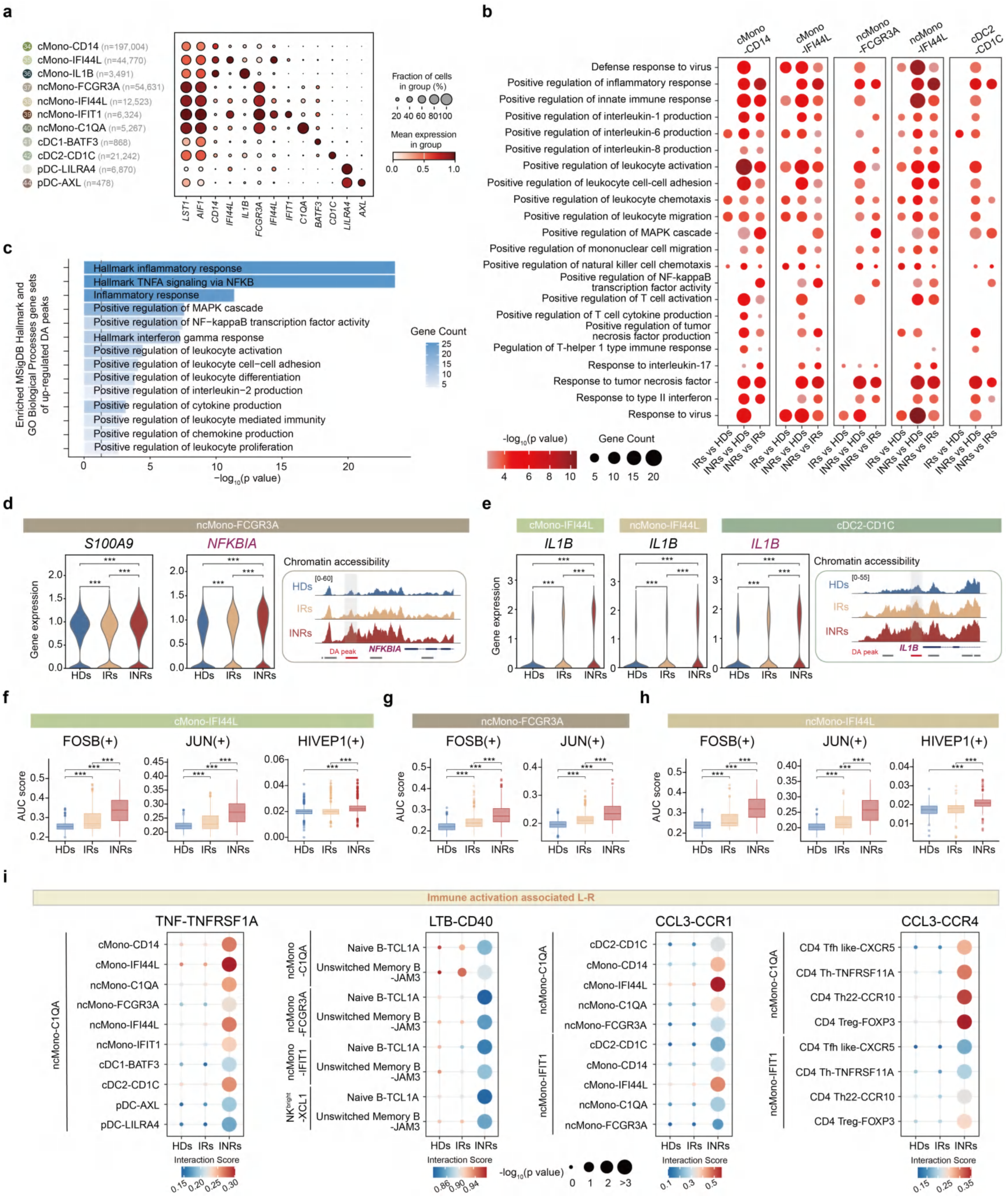
Abnormal activation of inflammatory responses in myeloid cells in INRs. **a**, Bubble plot showing the expression of marker genes used to annotate L3 Myeloid cells. **b**, The bubble plot showing the biological processes enriched by GO analysis for up-regulated DEGs in each group of five cell types. The color of the dots indicates the statistical significance of the enrichment, and the size of the dots indicates the number of genes annotated for each term. **c**, The bar graph showing the top 14 pathways enriched by genes annotated with up-regulated DA peaks. **d**, Violin plots showing gene expression of *S100A9* and *NFKBIA* and scATAC-seq tracks revealing gene structure and significant DA peaks for *NFKBIA* in ncMono-FCGR3A in HDs, IRs, and INRs. Different groups are shown in different colors. **e**, Violin plots showing gene expression of *IL1B* and scATAC-seq tracks revealing gene structure and significant DA peaks for *IL1B* in cMono-IFI44L, ncMono-IFI44L and cDC2-CD1C in HDs, IRs, and INRs. Different groups are shown in different colors. **f**, Box plots showing the AUC values of FOSB, JUN, and HIVEP1 in cMono-IFI44L in HD, IRs, and INRs. Different groups are shown in different colors. **g**, Box plots showing the AUC values of FOSB and JUN in ncMono-FCGR3A in HD, IRs, and INRs. Different groups are shown in different colors. **h**, Box plots showing the AUC values of FOSB, JUN, and HIVEP1 in ncMono-IFI44L in HD, IRs, and INRs. Different groups are shown in different colors. **i**, The bubble plots showing the changes in potential immune activation-related ligand-receptor pairs between immune cell subsets in HDs, IRs, and INRs, including TNF-TNFRSF1A, LTB-CD40, CCL3-CCR1, and CCL3-CCR4. The color of the dots indicates the interaction score calculated using CellPhoneDB, and the size of the dots indicates the -log_10_(p value).

**Supplementary Figure 7.**
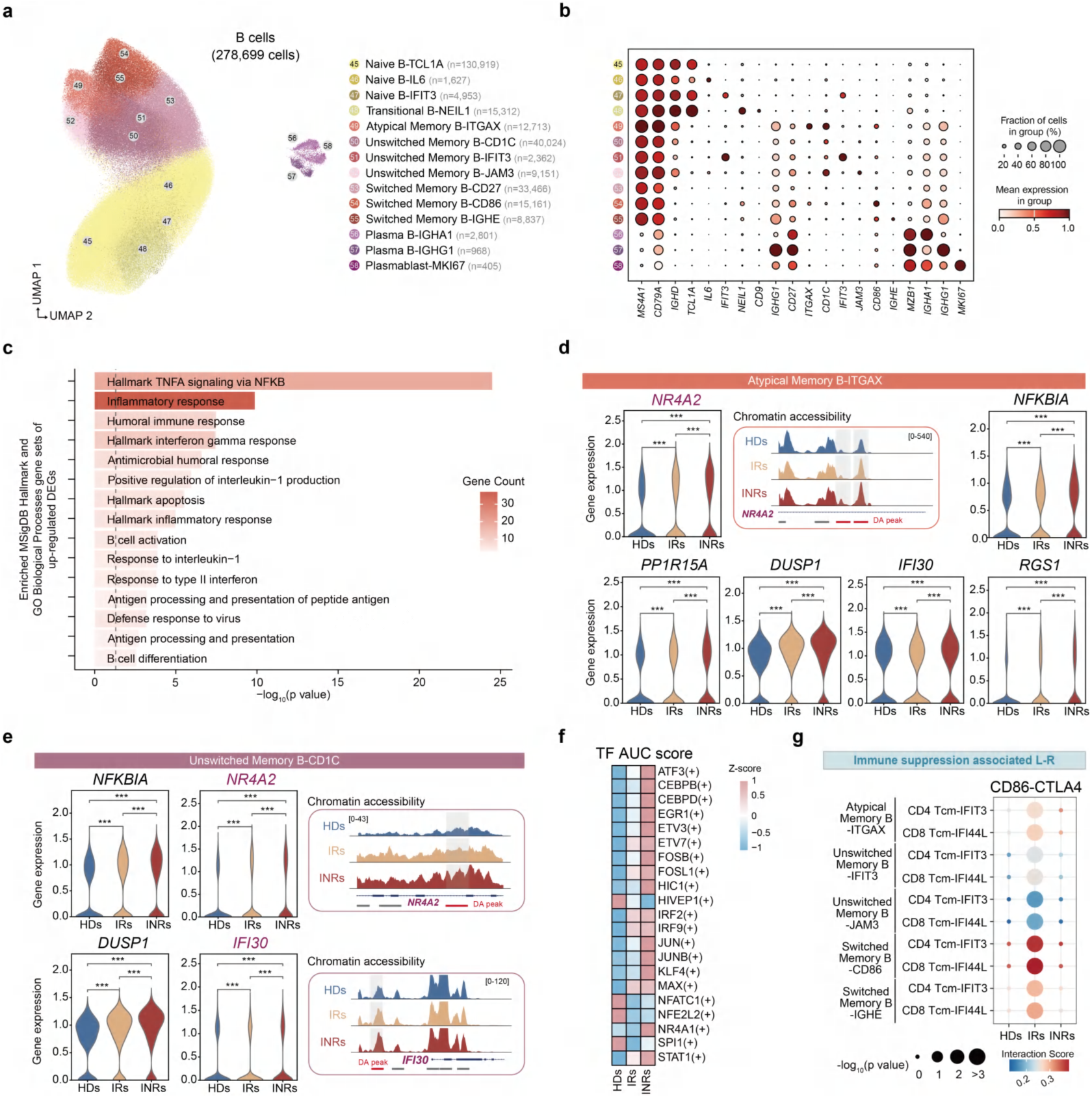
Abnormal function of B cells in INRs. **a**, The UMAP plot showing 278,699 B cells from scRNA-seq data, which are divided into 14 cell subtypes, with each cell type represented by a different color. **b**, Bubble plot showing the expression of marker genes used to annotate L3 B cells. **c**, The bar graph showing the top 15 pathways enriched by up-regulated DEGs. **d**, Violin plots showing gene expression of *NR4A2*, *NFKBIA*, *PP1R15A*, *DUSP1*, *IFI30*, and *RGS1* and scATAC-seq tracks revealing gene structure and significant DA peaks for *NR4A2* in Atypical Memory B-ITGAX in HDs, IRs, and INRs. Different groups are shown in different colors. **e**, Violin plots showing gene expression of *NR4A2*, *NFKBIA*, *DUSP1*, and *IFI30* and scATAC-seq tracks revealing gene structure and significant DA peaks for *NR4A2* and *IFI30* in Unswitched Memory B-CD1C in HDs, IRs, and INRs. Different groups are shown in different colors. **f**, Heatmap showing scaled mean AUC score of differential TFs in B cells in HDs, IRs, and INRs. **g**, The bubble plots showing the changes in potential immune suppression-related ligand-receptor pairs between immune cell subsets in HDs, IRs, and INRs, including CD86-CTLA4. The color of the dots indicates the interaction score calculated using CellPhoneDB, and the size of the dots indicates the -log_10_(p value).

**Supplementary Figure 8.**
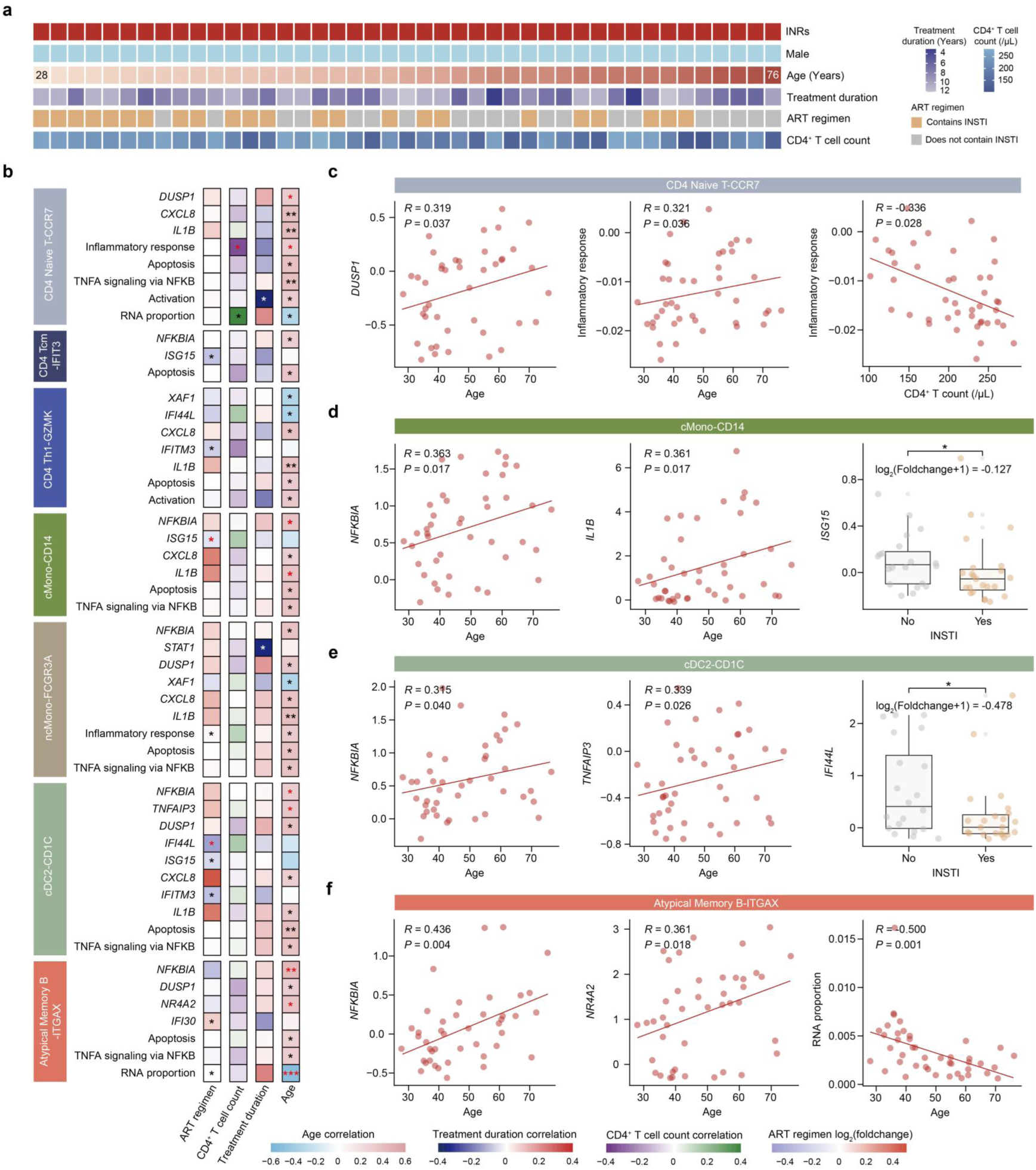
Clinical factors affecting immune reconstitution in INRs. **a**, The 43 INRs are arranged in ascending order by age and show whether different types of data were collected for each individual including sex, treatment duration, ART regimen, and CD4^+^ T cells count. **b**, Heatmaps showing the residual correlations of key gene expressions, pathway scores, and RNA cell proportions in eight cell types of INRs with four clinical information. **c**, Scatter plot showing genes and pathways significantly associated with age and CD4^+^ T cell count in CD4 Naive T-CCR7 of INRs. **d**, Scatter plot showing genes significantly associated with age and ART regimen in cMono-CD14 of INRs. **e**, Scatter plot showing genes significantly associated with age and ART regimen in cDC2-CD1C of INRs. **f**, Scatter plot showing genes significantly associated with age and RNA proportion in Atypical Memory B-ITGAX of INRs.

**Supplementary Figure 9.**
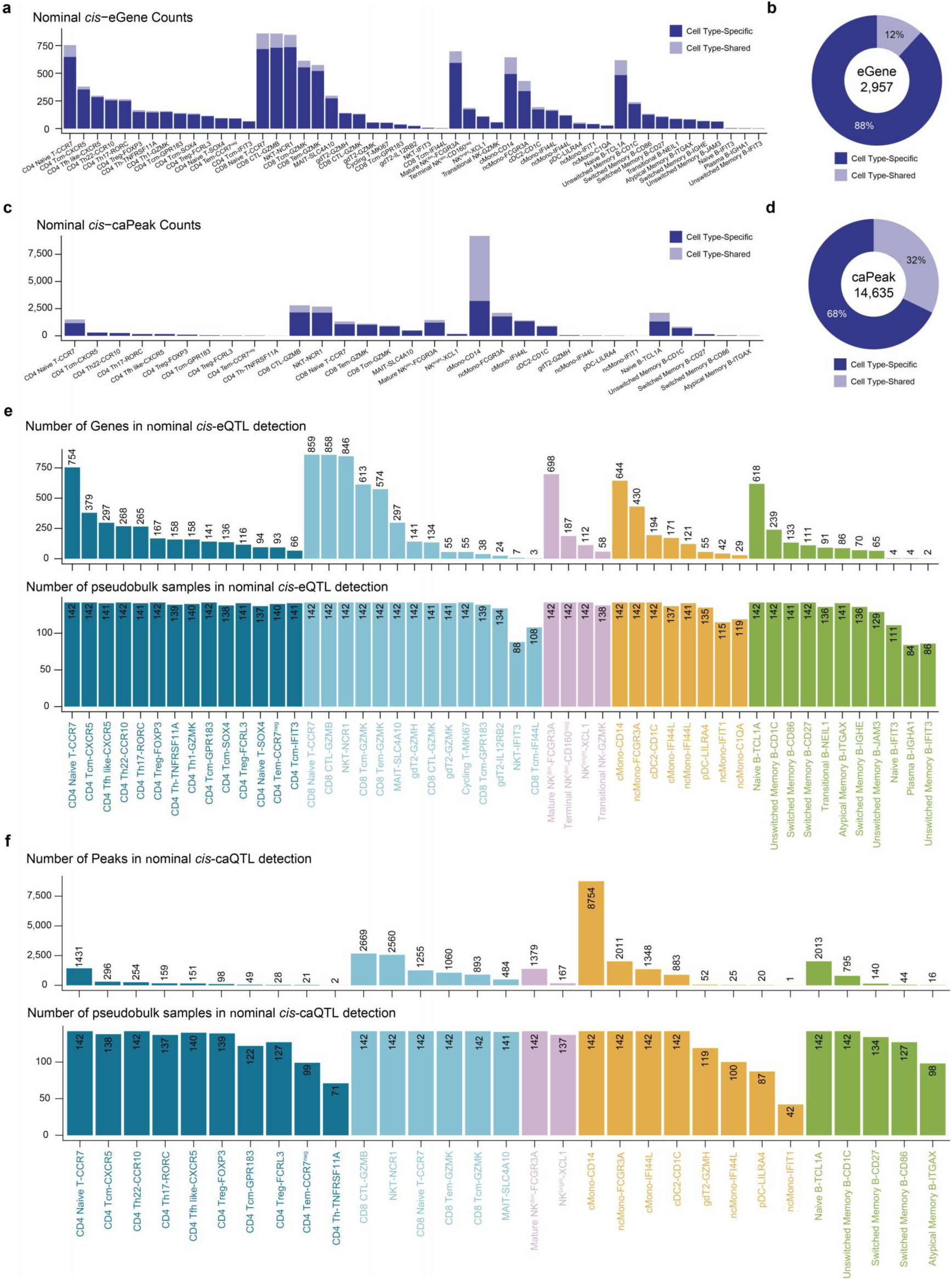
Summary of nominal *cis*-eQTL and *cis*-caQTL in immune cell types. **a**, The bar graph showing the number of cell type-shared and cell type-specific nominal *cis*-eGenes detected in the 51 cell types included in the nominal *cis*-eQTL analysis. **b**, The pie chart showing the distribution of significantly detected cell type-shared and cell type-specific nominal *cis*-eGenes. **c**, The bar graph showing the number of cell type-shared and cell type-specific nominal *cis*-caPeaks detected in the 31 cell types included in the nominal *cis*-caQTL analysis. **d**, The pie chart showing the distribution of significantly detected cell type-shared and cell type-specific nominal *cis*-caPeaks. **e**, Numbers of genes and pseudobulk samples used for nominal *cis*-eQTL analysis. **f**, Numbers of peaks and pseudobulk samples used for nominal *cis*-caQTL analysis.

**Supplementary Figure 10.**
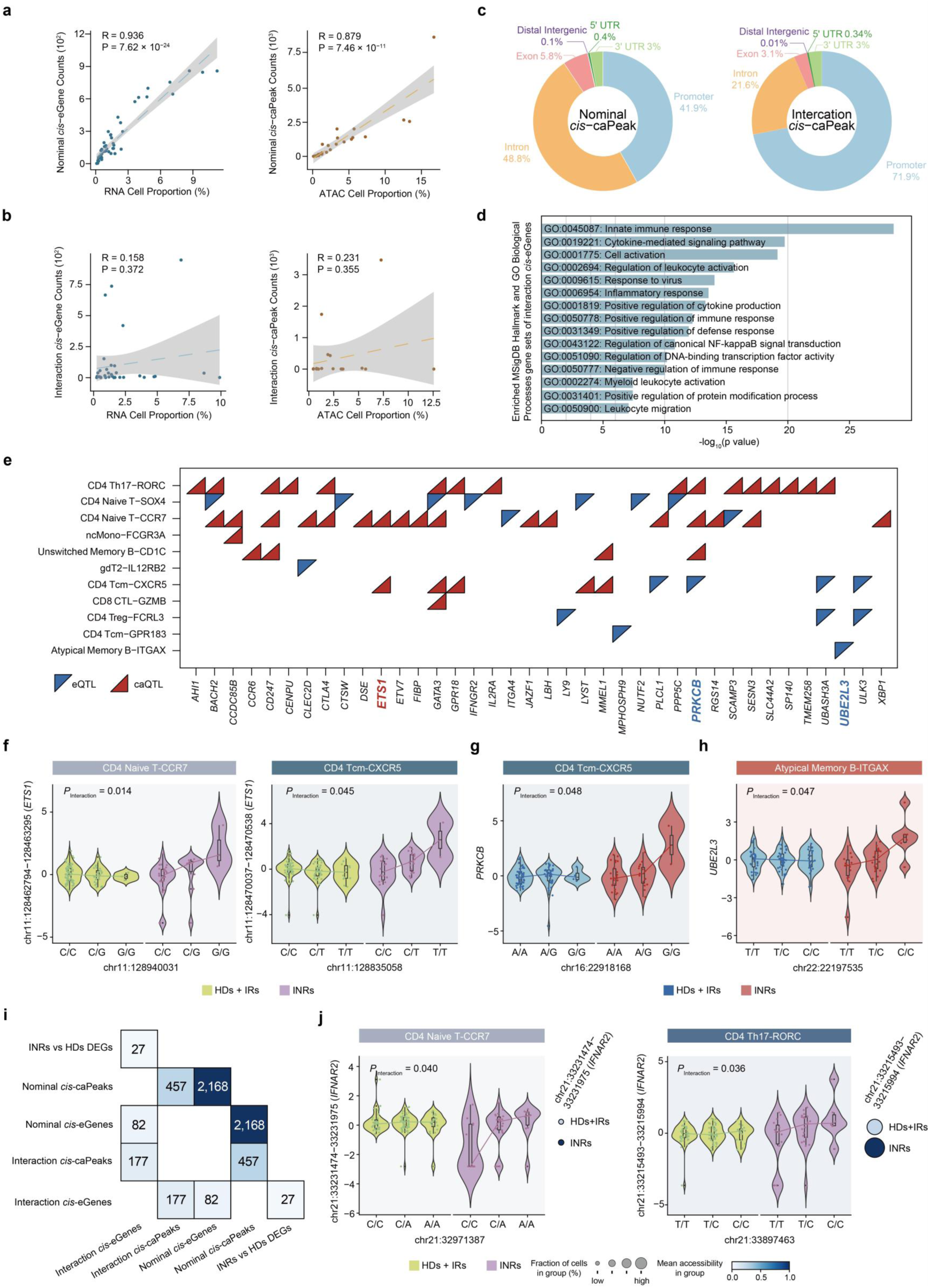
The supplement information of interaction *cis*-xQTLs analysis. **a**, The scatter plots showing the correlation between the nominal *cis*−eGene Counts and the corresponding RNA Cell Proportion and the correlation between the nominal *cis*−caPeak Counts and the ATAC Cell Proportion for each cell type. **b**, The scatter plots showing the correlation between the interaction *cis*−eGene Counts and the corresponding RNA Cell Proportion and the correlation between the interaction *cis*−caPeak Counts and the ATAC Cell Proportion for each cell type. **c**, The circle diagrams showing the proportion of regions annotated by the nominal *cis*-caPeaks and interaction *cis*-caPeaks on the genome. **d**, The bar graph showing the top 15 pathways enriched by interaction *cis*−eGenes. **e**, Shared genes among leading interaction *cis*-xQTLs and 76 risk genes associated with autoimmune diseases. **f**, Violin plots showing the *cis*-caQTL effects of chr11:128940031 and chr11:128835058 on chr11:128462794−128463295 (*ETS1*) and chr11:128470037−128470538 (*ETS1*) accessibility in CD4 Naive T-CCR7 and CD4 Tcm-CXCR5 in INRs and HDs+IRs control. All samples are shown as individual points. **g**, Violin plots showing the *cis*-eQTL effect of chr16:22918168 on *PRKCB* expression in CD4 Tcm-CXCR5 in INRs and HDs+IRs control. All samples are shown as individual points. **h**, Violin plots showing the *cis*-eQTL effect of chr22:22197535 on *UBE2L3* expression in Atypical Memory B-ITGAX in INRs and HDs+IRs control. All samples are shown as individual points. **i**, Heatmap showing the overlap between *cis*-eGenes, *cis*-caPeaks and DEGs. **j**, Violin plots showing the *cis*-caQTL effects of chr21:32971387 and chr21:33897463 on chr21:33231474−33231975 (*IFNAR2*) and chr21:33215493−33215994 (*IFNAR2*) accessibility in CD4 Naive T-CCR7 and CD4 Th17-RORC in INRs and HDs+IRs control. All samples are shown as individual points. Dot plots showing the peak accessibility of chr21:33231474−33231975 and chr21:33215493−33215994 in HDs+IRs and INRs.

**Supplementary Figure 11.**
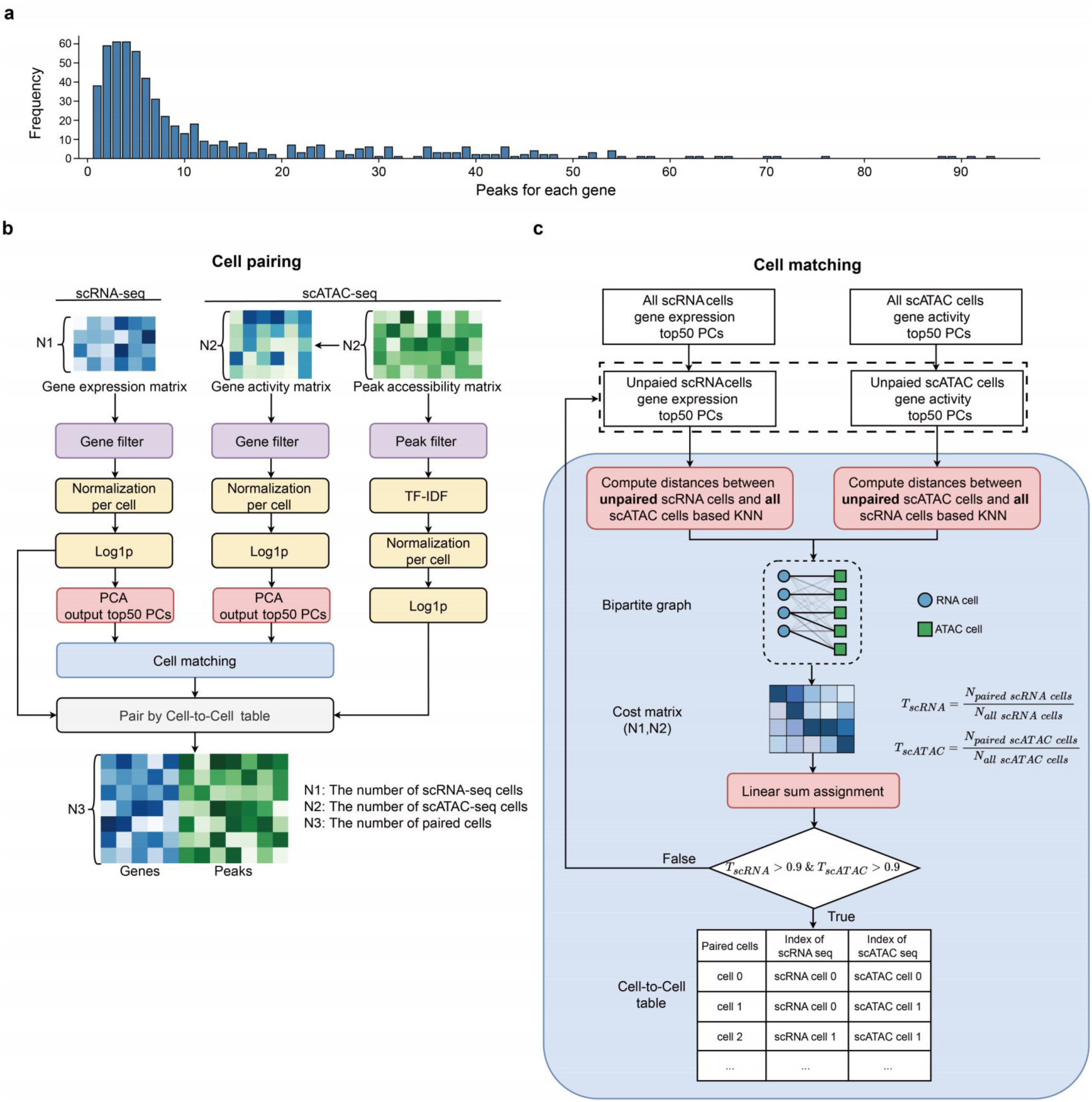
Workflow for processing and pairing of single-cell multi-omic data. **a**, Distribution of the number of chromatin accessibility peaks associated with each gene. **b**, Schematic overview of the workflow used to pair individual cells between the scRNA-seq and scATAC-seq modalities. **c**, Detailed flowchart illustrating the implementation of the cell pairing module introduced in (**b**).

**Supplementary Figure 12.**
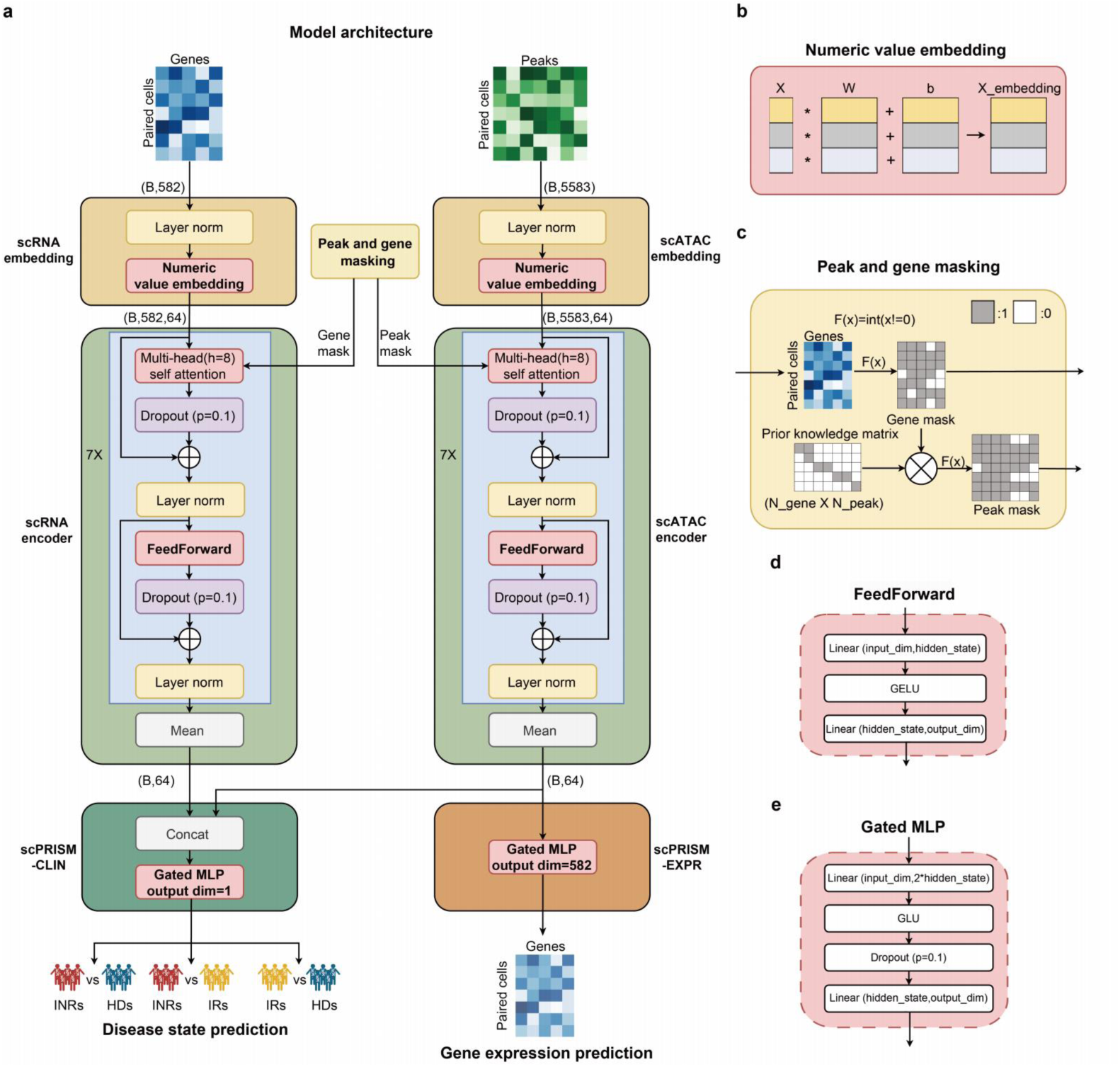
Architectural details of the scPRISM framework. **a**, Overview of the scPRISM architecture. Separate seven-layer masked self-attention encoders process the scRNA-seq and scATAC-seq data to generate distinct embeddings for each modality. These embeddings are then passed to two specialized modules: scPRISM-CLIN for disease state classification and scPRISM-EXPR for gene expression prediction. B: batch size during mode training. **b**, The Numeric Value Embedding module. This block transforms scalar input values (i.e., gene expression or peak accessibility) into high-dimensional vectors using feature-specific (per-gene and per-peak) learnable weights. **c**, The Peak and Gene Masking module. This component generates attention masks that restrict interactions to expressed genes and nearby chromatin peaks, defined as those within a ±250 kb window of the gene body. These masks are then applied within the multi-head attention layers to constrain the model’s focus to biologically relevant features. **d**, The Feed-Forward Network (FFN) block. Positioned after each multi-head attention layer. **e**, The Gated Multilayer Perceptron (MLP) module. This block, which utilizes a Gated Linear Unit (GLU), serves as the final prediction head for both the scPRISM-CLIN and scPRISM-EXPR modules, mapping the final embeddings to the desired output.

**Supplementary Figure 13.**
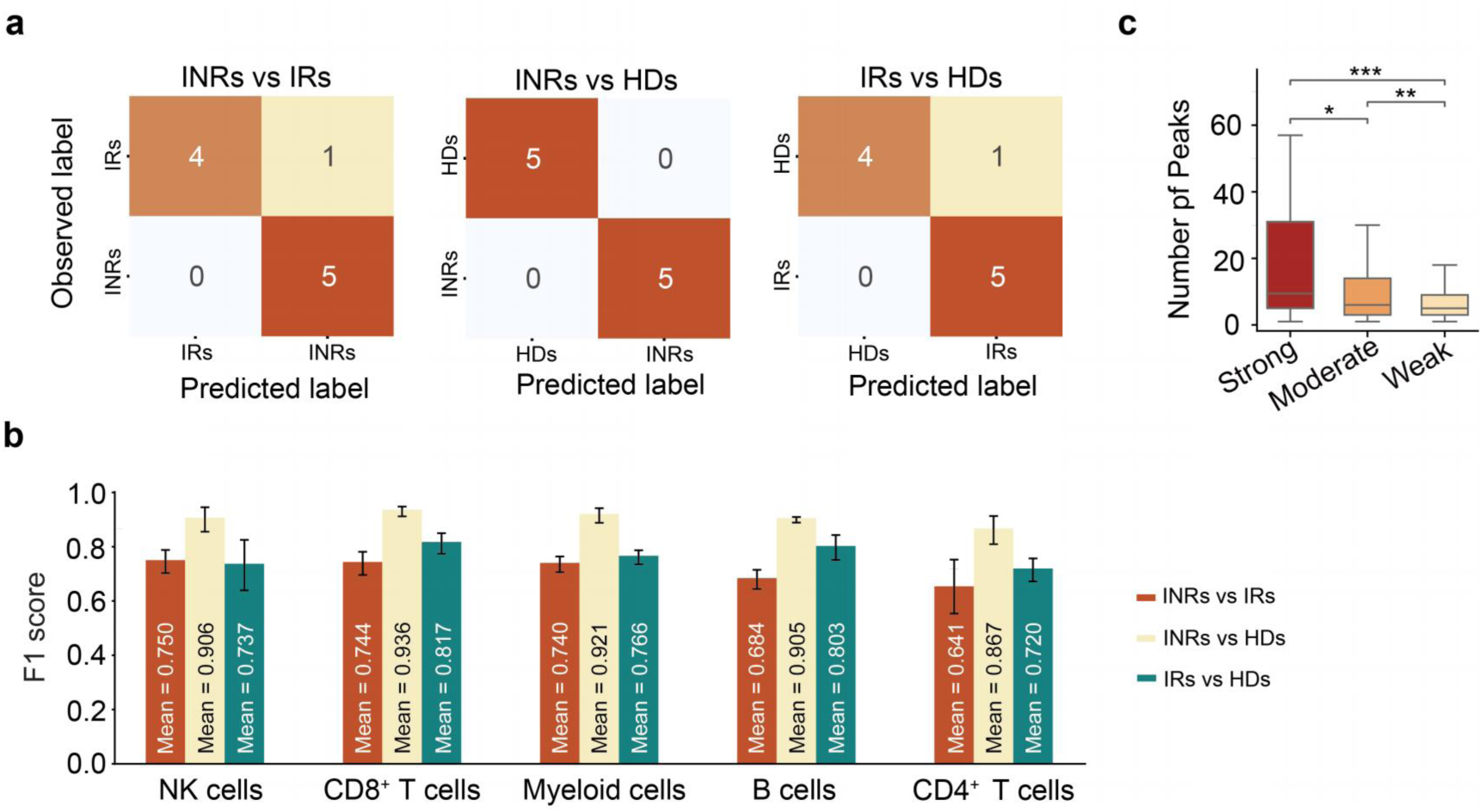
Performance evaluation of the scPRISM-CLIN and scPRISM-EXPR modules. **a**, Confusion matrix depicting the classification performance of the scPRISM-CLIN model on the held-out test set for discriminating between HIV disease states. **b**, Mean F1 scores for pairwise classification (INRs vs IRs, INRs vs HDs, and IRs vs HDs) by the scPRISM-CLIN module across major (L1) cell types. Error bars represent standard deviation. **c**, Comparison of the number of associated chromatin peaks for genes, stratified by the prediction accuracy (strong, moderate, or weak Spearman correlation) of the scPRISM-EXPR module. Significance were determined by a two-sided Mann-Whitney U-test with Benjamini-Hochberg correction, *P < 0.05, **P < 0.01, ***P < 0.001.

**Supplementary Figure 14.**
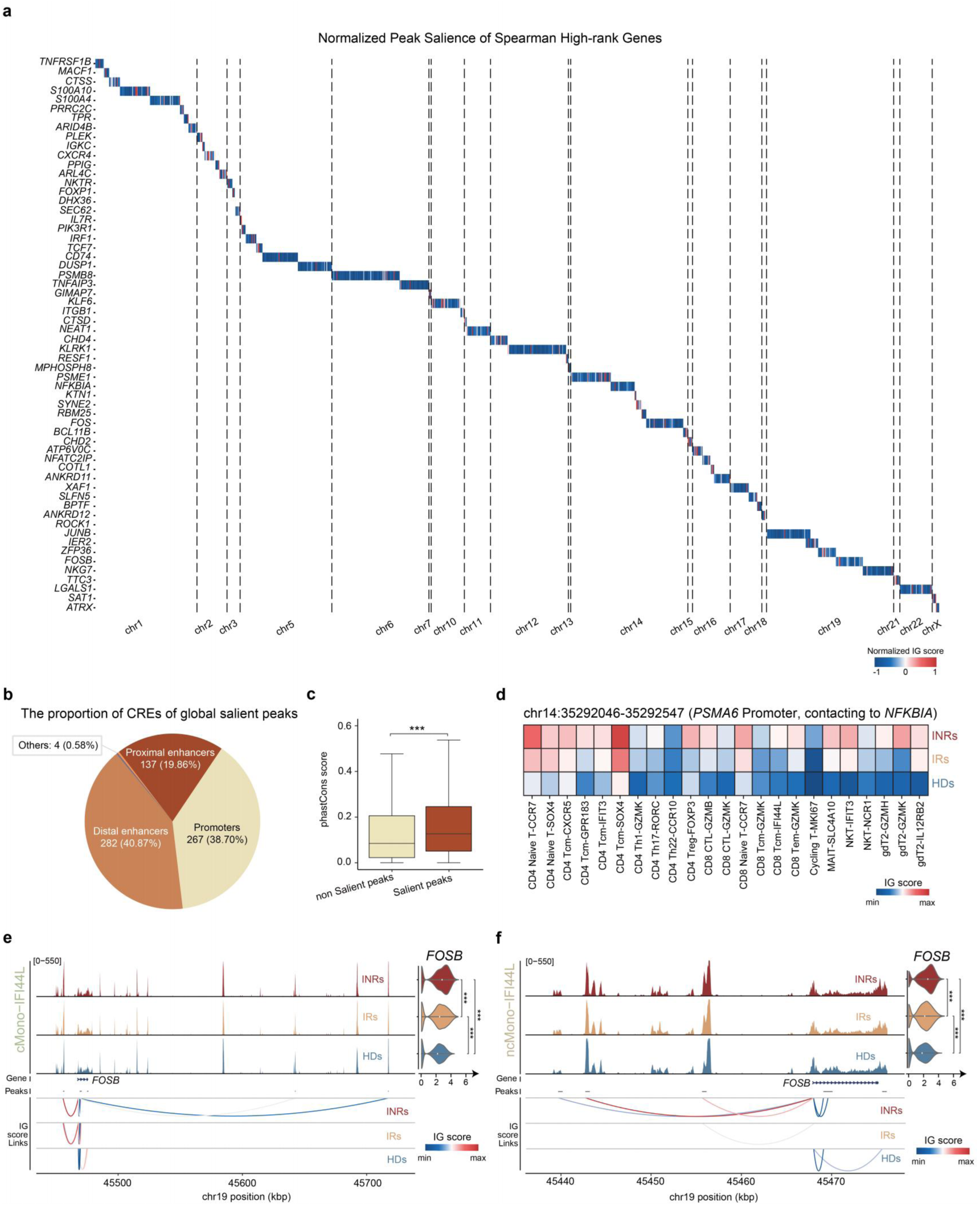
Analysis of genes and regulatory elements important for HIV disease states. **a**, Heatmap of peak importance scores, calculated using Integrated Gradients (IG), for the highly predictable genes (n = 60). The IG scores for all peaks associated with a given gene are min-max scaled to a range of -1 to 1. **b**, Genomic annotation of peaks. **c**, Comparison of evolutionary conservation (phastCons scores) between salient peaks (defined as peaks with IG scores > 0.005) and non-salient peaks. Significance was determined by a two-sided Mann-Whitney U-test with Benjamini-Hochberg correction, *P < 0.05, **P < 0.01, ***P < 0.001. **d**, Heatmap showing the IG scores of a peak located at chr14:35292046-35292547 (the *PSMA6* promoter) contacting to *NFKBIA* gene. The scores are displayed across different patient cohorts and various T and NK cell subtypes. **e**-**f**, Cell type-specific chromatin peaks interacting with *FOSB* in cMono-IFI44L for (**e**) and ncMono-IFI44L for (**f**). In each panel, arc plots visualize significant peak-to-gene interactions, while violin plots compare *FOSB* expression across patient cohorts. Inset box plots show the median, the 25th-75th percentiles, and whiskers extending to 1.5 times the interquartile range. Significance was determined by a two-sided Mann-Whitney U test, ***P < 0.001, **P < 0.01, and *P < 0.05.

**Supplementary Figure 15.**
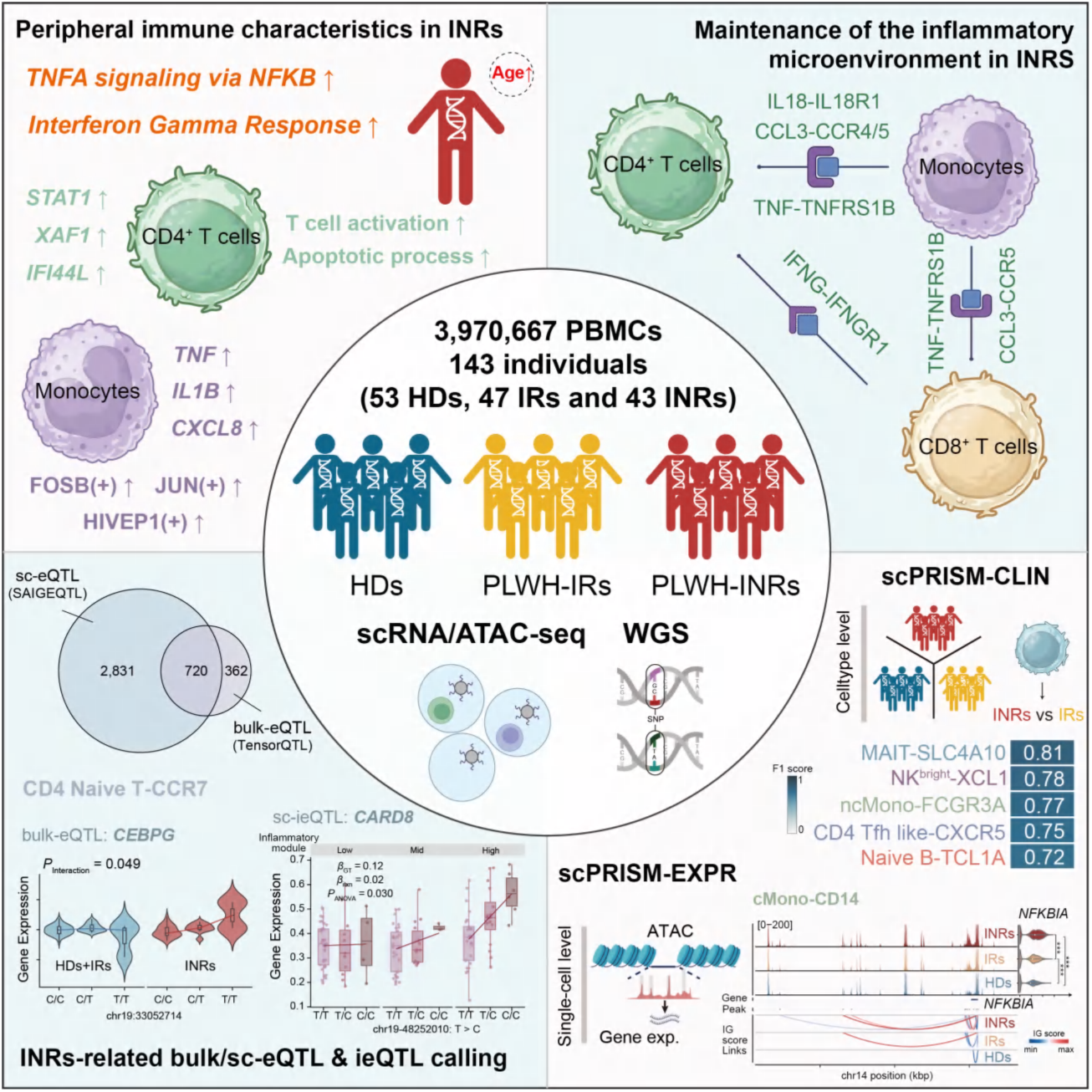
Graphical abstract of our study.

## Supplementary Tables

Supplementary Table 1. Sample clinic information

Supplementary Table 2. Immune cell annotation marker genes

Supplementary Table 3. Immune cell differentially expressed genes in HDs, IRs and INRs

Supplementary Table 4. Immune cell differentially accessible peaks in HDs, IRs and INRs

Supplementary Table 5. Pathway gene sets

Supplementary Table 6. Immune cell differential TFs activity in HDs, IRs and INRs

Supplementary Table 7. Immune cell ligand receptor enrichment in HDs, IRs and INRs

Supplementary Table 8. Summary of lead nominal *cis*-xQTLs in immune subtypes

Supplementary Table 9. Summary of lead interaction *cis*-xQTLs in immune subtypes

Supplementary Table 10. Summary of sc-eQTLs in CD4^+^ T cells

Supplementary Table 11. Summary of sc-ieQTLs in CD4^+^ T cells

